# Enzyme-driven phase separation of synthetic condensates enables self-organizing compartments and protective microenvironments

**DOI:** 10.64898/2025.12.24.696465

**Authors:** Archishman Ghosh, Advait Thatte, Surased Suraritdechachai, Roman Rattunde, Christoph A. Weber, T-Y Dora Tang

**Author notes:** Correspondence should be addressed to Dora Tang and Christoph A. Weber. Authors contributed equally (Archishman Ghosh, Advait Thatte).

## Abstract

Compartmentalization via phase separation is increasingly recognized as a dynamic regulator of cellular biochemistry, yet the consequences of coupling enzymatic reactions to compartmentalization remain poorly understood. We combine a minimal, cell-free expression system with kinetic modeling to show how coupling enzymatic activity to biomolecular condensates drives emergent self-regulation via droplet formation and dissolution. In-vitro mRNA transcription induces phase separation with an intrinsically disordered protein (mutant G3BP1), forming condensates that modulate transcription and degradation kinetics. Our kinetic model shows that phase separation reduces rate constants, with slower degradation within condensates than in the mRNA-protein-poor phase. This leads to more mRNA with a prolonged lifetime of mRNA relative to the case without condensates. Extending the model to sustained and oscillatory resource supply reveals that condensates elevate mean mRNA levels and buffer deviations from the mean compared to the non-condensate scenario. These findings provide a general mechanism of cross-regulation and feedback between phase separation and enzymatic networks, highlighting condensates as active regulators of biochemical flux rather than as passive organizers.

**GRAPHICAL ABSTRACT:** 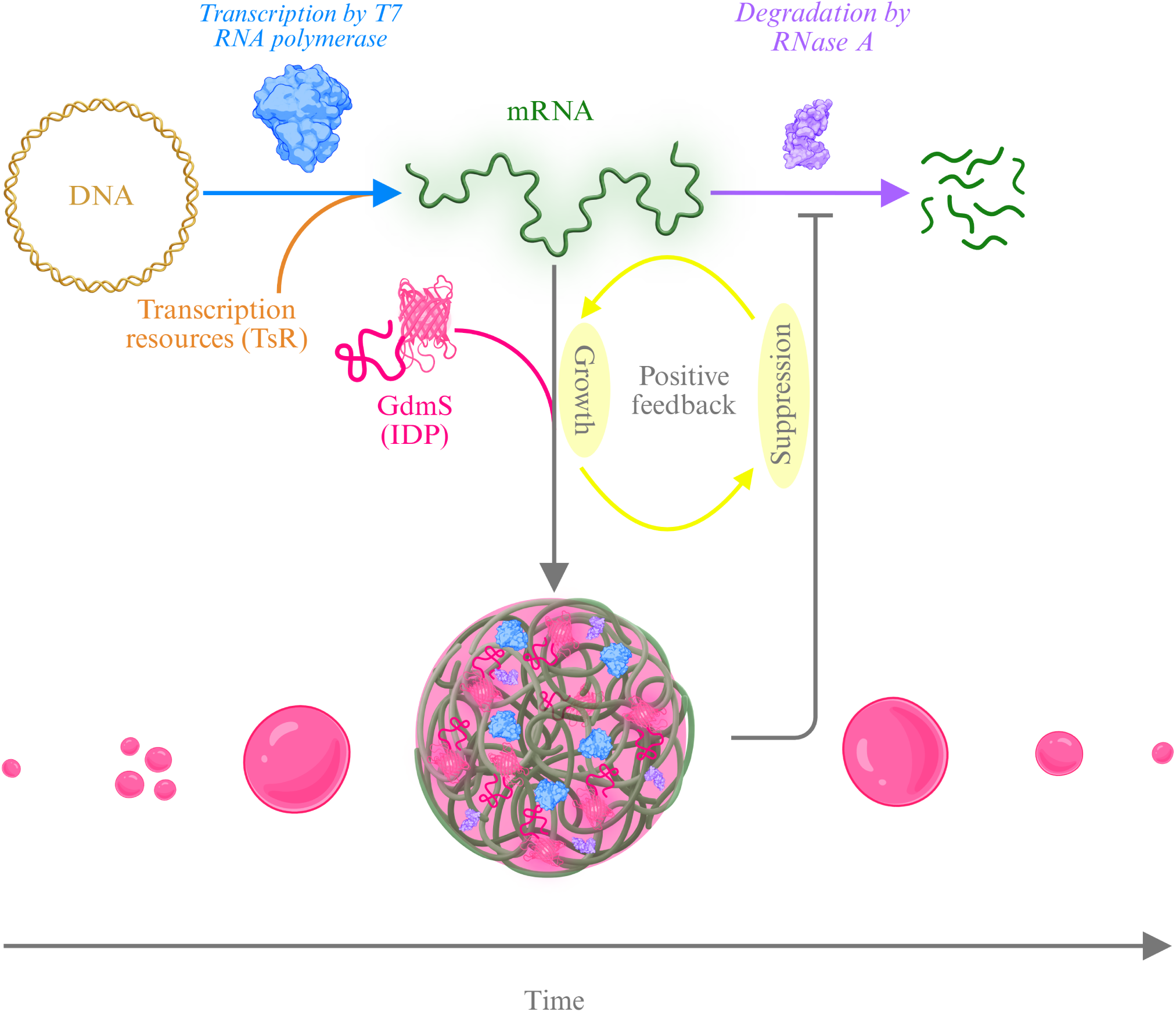

## INTRODUCTION

Living systems are defined by their ability to sustain out-of-equilibrium behavior, a property that is underpinned by dynamic cellular functions such as metabolism, transcription, and translation [1, 2, 3]. From microbes to mammals, these dynamic functions are further governed by complex, interwoven reaction networks that respond to environmental stimuli and regulate biochemical activity [4]. While molecular complexity and reaction networks enable the persistence of nonequilibrium states, the role of compartmentalization in modulating these networks is still poorly understood.

Biomolecular condensates formed via liquid-liquid phase separation offer a promising framework to explore feedback mechanisms that might arise from the coupling between enzyme reactions and compartmentalisation [5, 6, 7]. These mesoscale compartments generate heterogeneous environments within the cytoplasm and have been shown to influence reaction kinetics by concentrating biomolecules [7, 8, 9], reduce noise [10, 11], altering diffusion dynamics [12, 13], and by the selective partitioning of solutes [14, 15, 16, 17]. Recent advances have proposed a molecular grammar driving condensate formation and linked the onset and dissolution of condensates to enzymatic processes including transcription and translation [18], proteolysis [19], and phosphorylation cycles [20, 21]. In these cases, cycles of formation and degradation have been driven by external input at regular time intervals [21]. Functionally, condensates have been implicated in diverse cellular roles – from nucleating cytoskeletal structures [22, 23] to regulating translation via mRNA sequestration [24, 25].

Synthetic condensates composed of intrinsically disordered proteins, polyelectrolytes, and nucleic acids serve as minimal models for studying membrane-free compartmentalization [26, 27, 28, 29, 30]. These systems allow precise control over chemical composition and material properties, and are amenable to quantitative measurements of enzyme kinetics, molecular concentrations, and viscosity. Moreover, synthetic condensates have been extended to nonenzymatic reactions, positioning them as key tools in origin-of-life and systems chemistry research [31, 32, 33, 34, 35]. Despite these advances, a general framework for feedback regulation via coupling between enzyme reactions and compartmentalization has not yet been demonstrated nor quantitatively characterized.

In this work, we use bottom-up design to construct a minimal synthetic system under cell free conditions in which synthetic condensates form and dissolve autonomously, without external intervention. This behavior is mediated by the compartment, creating a feedback on the enzyme kinetics. Using cell-free expression systems, we produce mRNA that phase separates with G3BP1, a well-known scaffold for mesoscale organization [36, 20]. We develop a two-phase model that accounts for resource-limited transcription and translation, relying on recently developed theoretical frameworks [37, 38, 6]. Fitting the model to our experimental data, we extract phase-dependent degradation and transcription rate coefficients. A key finding is that phase separation suppresses mRNA degradation by lowering the degradation rate inside the condensates. The consequence is that, when transcriptional resources are depleted, condensates dissolve, but on a much longer timescale than in the absence of phase separation. Thus, phase separation mediates a distinct feedback mechanism between condensation, mRNA transcription, and degradation. When using our model to account for periodic resource input, we observe oscillations in mRNA levels and condensate volume, with condensates mediating a buffering effect that reduces oscillations in mRNA concentrations. In summary, by combining theory and experiments and integrating feedback between enzymatic reactions and phase separation into a unified quantitative model, we demonstrate self-regulated mesoscale organization in minimal cell free environments. While our system employs condensate-based confinement reminiscent of protocells, the underlying principles extend to other compartmentalized architectures, including lipid vesicles.

## RESULTS

### Experimental cell-free system

We established a cell-free expression system that autonomously generates and dissolves biomolecular condensates triggered by mRNA transcription and degradation (Figure 1 and Supplementary figure 1). The system is based on a commercially available insect cell-free transcription–translation extract derived from *Spodoptera frugiperda* (Sf21) cells (TNT extract). The extract was programmed to synthesize mRNA that phase separates with a mutant form of the intrinsically disordered protein G3BP1 (ΔE1/ΔE2), fused to the red fluorescent protein mScarlet-I (hereafter referred to as GdmS) (Supplementary methods 9.2). To drive coupled condensate formation and degradation, TNT extract was supplemented with 5.6 nM plasmid DNA encoding GdmS and 5 *µ*M purified GdmS protein. 1 *µ*M T7 RNA polymerase, and 0.1 *µ*M RNase A were also added to enhance transcription and degradation rates. A complete list of materials is provided in supplementary methods section 1, and the associated mRNA sequence and GdmS sequence in section 2.For plasmid design and preparation, see section 3, and for purification and labeling protocol of mRNA and GdmS, refer to section 4, section 5, section 6 and table 2.

**Figure 1:**
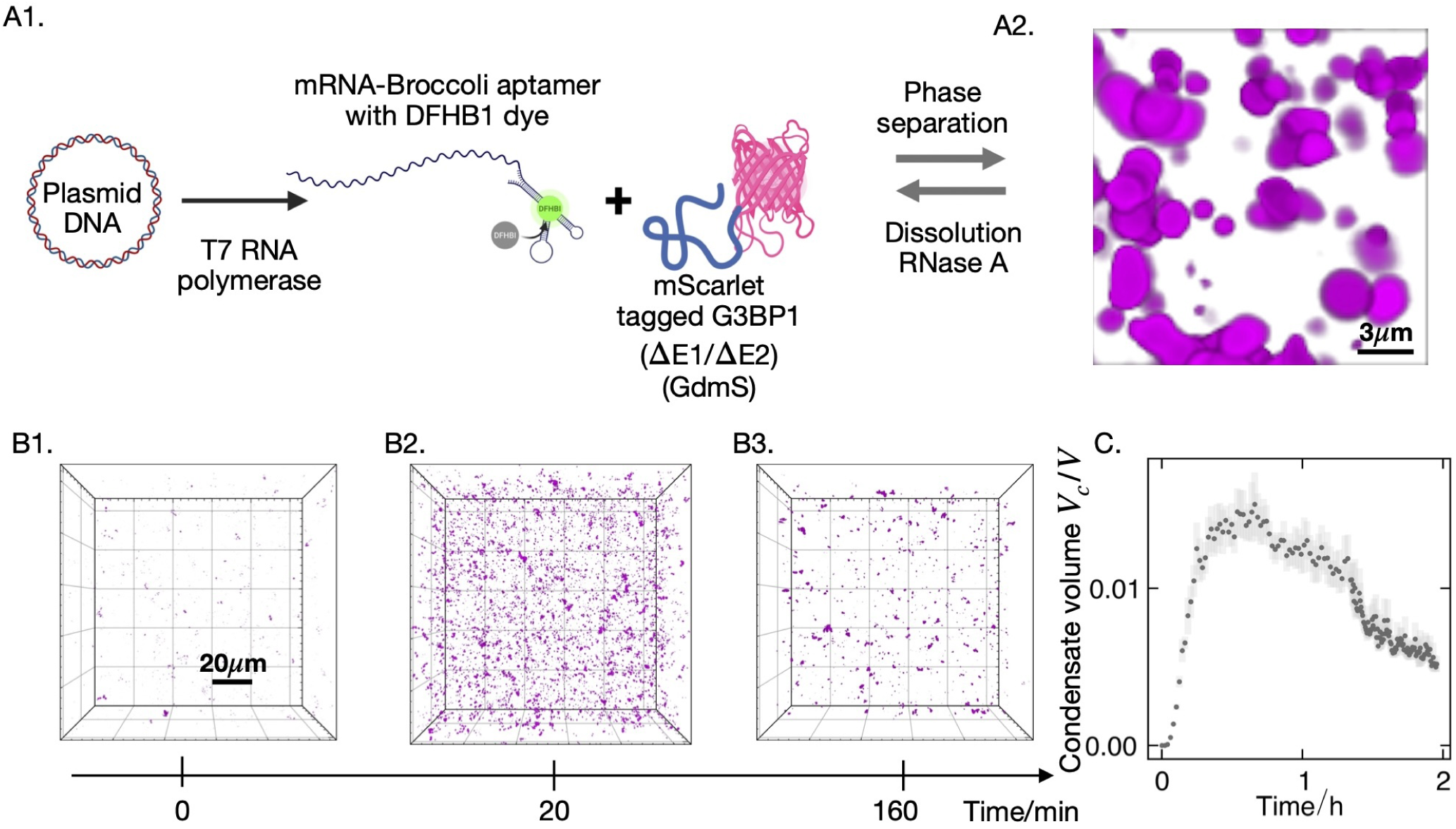
Condensates form and dissolve as mRNA is transcribed and subsequently degraded. (A1) Schematic of cell-free transcription and phase separation of mRNA with protein, followed by condensate dissolution by RNase A. (A2) Three-dimensional rendering of confocal *z*-stacks for 6 *µ*M GdmS, 1 *µ*M T7 polymerase, and 0.1 *µ*M RNase A in 80% TNT extract at 30 *^◦^*C. (B1–B3) Confocal three-dimensional micrographs over time showing cell-free expression–driven formation and dissolution of mRNA–GdmS condensates. The system was doped with 6 *µ*M GdmS, 1 *µ*M T7 polymerase, and 0.1 *µ*M RNase A. (C) Kinetic trace of condensate volume fraction over time for the same confocal time-lapse experiment.

Time-resolved 3D confocal microscopy stacks (see Supplementary methods section 7 and section 10) revealed in situ condensate formation and subsequent dissolution over 1.5 h, with maximal condensate formation observed approximately 0.5 h after the onset of the reaction (Figure 1C). Using 3D surface reconstruction as described in Ref. [15], we converted time-resolved 3D confocal stacks into traces of condensate volume fraction (*V_f_* ) , defined as the ratio of the total condensate volume (*V_c_*) within the observation field to the total volume of that field (*V* ), *V_f_* = *V_c_/V* . These time traces revealed the temporal dynamics of condensate growth and dissolution (Figure 1B1-B3,C)

Confocal dual-channel fluorescence micrographs of GdmS condensates and mRNA confirmed co-localization of mRNA and protein within the condensates. The mRNA, containing the Broccoli aptamer and bound to DFHBI, was detected in the GFP channel, whereas GdmS was detected in the RFP channel (Figure 2A,B).

**Figure 2:**
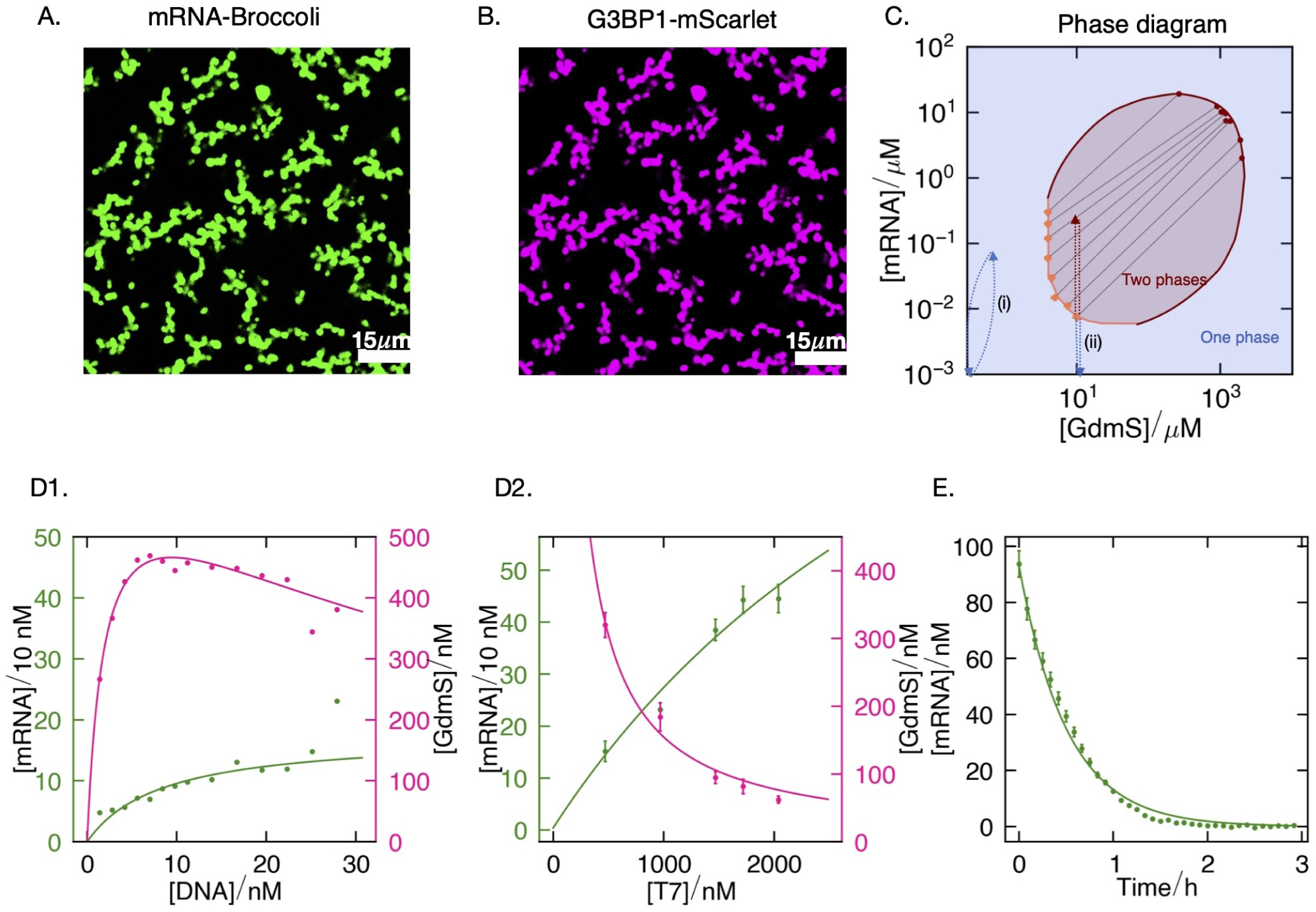
Evolution of condensate volume is governed by the phase diagram and transcription and degradation rates. (A) Fluorescence micrographs of GdmS condensates showing mRNA fluorescence from the Broccoli aptamer bound to DFHB1 dye. (B) mScarlet-I fluorescence from the GdmS fusion protein. Samples were prepared with 100 nM mRNA, 10 *µ*M GdmS, and 20 *µ*M DFHB1 dye. (C) Phase boundaries/binodals between the one-phase (blue) and two-phase (pink) regions of purified GdmS and mRNA in TNT extract determined by confocal microscopy. Errors in the binodals are detailed in Supplementary methods sections 8.1 and 8.2. A phase space trajectory of the system in the one phase region is illustrated in (i). Upon doping with GdmS, the system enters the two phase region, leading to a higher maximal mRNA concentration for the same initial TsR concentration (ii). (D1–D2) Plots showing the maximum yield of mRNA and GdmS produced by the cell-free expression system as a function of (1) T7 polymerase concentration and (2) DNA concentration. (E) Time course of mRNA degradation driven by endogenous RNase A activity in TNT cell-free extract. Reactions contained 80% TNT extract supplemented with 100 nM mRNA and 20 *µ*M DFHB1, and fluorescence was monitored over time. All experiments were conducted at 30 *^◦^*C. Data points show mean values from triplicate measurements.

To find the correct conditions for spontaneous formation and dissolution of condensates we determined the concentrations for in situ phase separation between GdmS and mRNA by mapping the GdmS poor arm of the phase diagram by optical microscopy (Supplementary methods section 9.1). This was achieved by mixing purified mRNA and GdmS at defined concentrations in 80% TNT extract (in the absence of DNA) (see Supplementary methods section 8.1, Figure 5, and Figure 1). For completeness, we obtained the GdmS rich branch by quantifying condensate volumes from 3D confocal stacks (Supplementary methods section 8.2, Supplementary Figure 5) and applying principles of mass conservation (Figure 1B1-B3,Figure 2C). The minimal concentrations of RNA and GdmS to form condensates were determined to be [mRNA]_tot_ = 0.012 *µ*M and [GdmS]_tot_ = 4.5 *µ*M. The corresponding concentrations in the GdmS-rich phase were 10 *µ*M mRNA and 1000 *µ*M GdmS. These concentrations define the edges of the system’s closed-loop phase boundary, beyond which no condensates are observed. These measurements delineate the GdmS-mRNA binodal (GdmS poor and GdmS rich phases) and define the concentrations used for subsequent *in situ* phase-separation experiments.

Given that GdmS phase separates with its cognate mRNA in TNT extract, we next quantified mRNA and protein production to assess whether their levels are sufficient to drive *in situ* phase separation. We determined the maximal concentrations of mRNA and GdmS achievable in the cell-free system by varying the initial concentrations of T7 RNA polymerase (T7) (Figure 2D2) and plasmid DNA (Figure 2D1). Kinetic fluorescence measurements of mRNA synthesis, monitored via DFHBI–Broccoli fluorescence, showed that the mRNA concentration increased from 12.5 nM to 45 nM as T7 concentration was raised from 470 nM to 2040 nM. In contrast, the translated GdmS concentration decreased from 320 nM to below 10 nM over the same T7 range. This inverse relationship between mRNA and GdmS yields is consistent with competition for shared transcription–translation resources, whereby enhanced transcription at high T7 polymerase levels limits protein synthesis.

Plots of the maximum attainable mRNA and protein concentrations as a function of DNA template and T7 polymerase (supplementary note 9.3) demonstrate that production of both components is tunable by these inputs. Crucially, however, the cell-free expression system could not generate sufficient GdmS to reach the phase-separation threshold (4.5 *µ*M), even under conditions where ample mRNA was produced. To circumvent this limitation, we supplemented the reactions with purified GdmS before the onset (“doping”), and relied purely on *in situ* synthesis of mRNA to drive phase separation.

We next sought to establish the degradation part of the dynamic system by characterising the ability of the reaction mixture to degrade mRNA and thereby drive condensate dissolution. To test this, we added 100 nM purified mRNA to the TNT extract and monitored the decrease in mRNA concentration over time by fluorescence spectroscopy. The mRNA concentration, calibrated from DFHBI fluorescence, decayed with apparent first-order kinetics, with a decay constant of 0.007 ± 0.001 h^−1^. Indeed, pseudo–first-order decay analysis with RNase A titrations yielded an effective endogenous RNase A concentration of 270 ± 0.05 nM in the extract (Supplementary Figure 15). Having demonstrated that the system produces mRNA above the phase-separation threshold and that this mRNA undergoes spontaneous degradation in the extract, we established a rational experimental framework for a minimal system in which transcription and compartment feedback drive the spontaneous formation and dissolution of condensates.

### Model for transcription-degradation kinetics with and without condensates

We propose that condensate formation and dissolution are driven by transcription and degradation of mRNA (Figure 1). Further, the formation of the condensate provides a feedback mechanism that spatially and differentially modulates mRNA transcription and degradation within the protein-rich phase and the surrounding protein-poor phase, leading to condensate dissolution. To test this hypothesis, we developed a resource-limited mass action model to estimate the rates of transcription, translation, and mRNA degradation in both the single-phase (homogeneous) and phase-separated regimes of the system (for details, see supplementary methods section 11.1 to section 11.6.).

The model is composed of the following components: mRNA (a transcribed and a degraded species), GdmS (the translated IDP), TsR (the transcription resources), TlR (the translation resources), and DNA, along with the mRNA production and degradation enzymes, T7 RNA polymerase (T7) and RNase A (Figure 3(A)). For simplicity, all components except mRNA and GdmS were considered to be dilute, meaning that their effects on phase separation are neglected.

**Figure 3:**
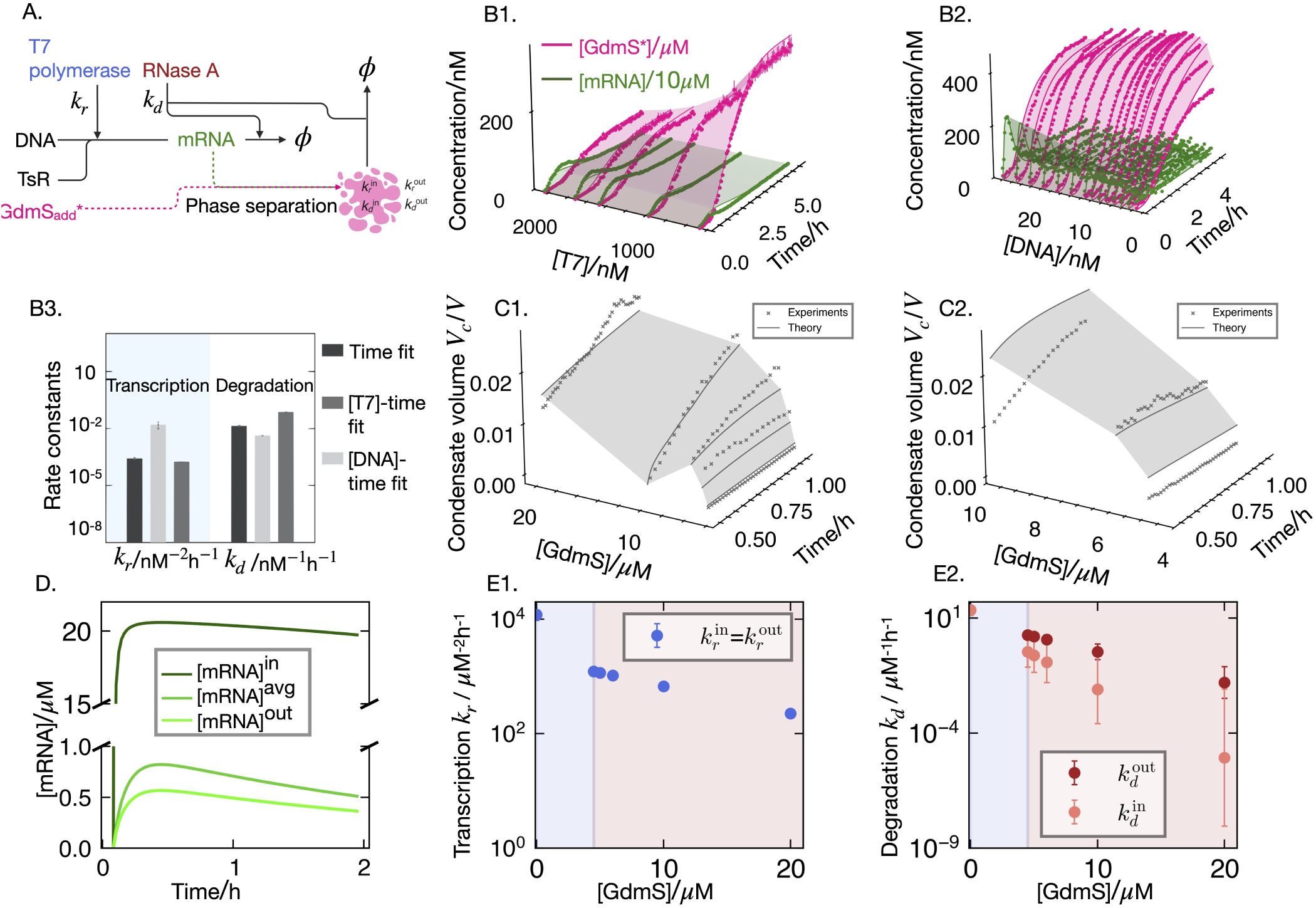
Rate constants extracted from the theoretical model indicate that RNA degradation is suppressed within the droplets. (A) Simplified reaction scheme for transcription-translation with rate constants and components for the two phase kinetic model. Condensation is denoted by the pink region. **(B)** Kinetic profiles of mRNA (green) and GdmS (pink) production over time for (B1) increasing T7 polymerase concentration and (B2) increasing DNA template concentration, obtained from multi-well plate experiments conducted at 30 *^◦^*C. Dots with error bars indicate experimental data, and the shaded surfaces represent global fits of the combined T7 and DNA titration datasets to a single-phase kinetic model. (B3) The transcription (*k_r_*) and degradation (*k_d_*) rate constants obtained from the global fit of the one phase kinetic model to the kinetic data. (C) Condensate volume fraction data (dots with error bars) obtained from analysis of 3D confocal timelapse as a function of GdmS concentration (0–20 *µ*M) in the absence (C1) and presence of 1 *µ*M T7 polymerase and 0.1 *µ*M RNase A (C2). Global fits to a two-phase kinetic model are shown as solid lines with a shaded gray surface. Data points show mean values and errors from triplicate measurements. (D) Example concentration–time trajectories of mRNA production and degradation inside and outside the condensates, and their average, obtained from our kinetic model. (E) Estimates of the transcription (E1) and degradation (E2) rate constants with increasing doped GdmS concentration obtained from simulation of the two phase kinetic model on the experimental data. The blue and shaded pink regions depict the one and two phase regions respectively. The transcription rate constants are identical in both phases of the condensate, whereas the degradation rate constants show distinct trends inside (orange) and outside (red). Error bars represent the fitting error in estimating the rate constants *k_r_* and *k_d_*.

The experiment and the model system are initially prepared without mRNA and contain a fixed, pre-doped concentration of GdmS. Transcription is triggered by adding DNA and T7 RNA polymerase at defined concentrations, such that mRNA is produced at a rate proportional to the transcription rate constant *k_r_* and the concentrations of transcription resources [TsR], transcription enzyme [T7], and the template [DNA]. As soon as mRNA is transcribed, it starts getting degraded by the action of RNase A, at a rate proportional to [mRNA] and [RNase]. The rate constant associated with degradation is denoted as *k_d_*. In the case where the system phase separated, GdmS was provided in excess relative to the total translation yield, and the translation dynamics were ignored. As the mRNA concentration increases and reaches the saturation concentration for phase separation, defined by the GdmS-poor binodal of the phase diagram, the system crosses into the two-phase region and undergoes phase separation, see Figure 2C for illustration of mRNA concentration trajectories. In this regime, the average concentrations of mRNA, GdmS, TsR, and TlR are partitioned between a GdmS-rich (droplet) phase and a GdmS-poor phase. Figure 2A,B show the up-concentration of mRNA and GdmS inside condensates, where mRNA is labeled using Broccoli-DFHB1 aptamer (green) and G3BP1 using m-Scarlet (magenta). Each phase is characterized by distinct transcription, degradation, and translation kinetics, with phase-specific rate constants.

An important feature of the model is that the average mRNA concentration is coupled to the condensate volume fraction through the phase diagram (Figure 2 and supplementary methods section 11.5). As transcription proceeds and the total mRNA concentration increases, the system moves along the binodals of phase diagram, leading to an increase in the condensate volume fraction. At the same time, the transcription and degradation rates of mRNA, governed by the kinetic rate constants *k_r_* and *k_d_*, in general differ between the GdmS-rich and GdmS-poor phases. Consequently, changes in the condensate volume fraction feed back onto the effective kinetics of mRNA production and decay, establishing a feedback loop between phase behaviour and mRNA dynamics.

In our model, the following assumptions were made: (1) Transcription and degradation occur in both phases. (2) mRNA and GdmS constitute the scaffold of the condensate phase, while dilute components (clients) may partition between phases according to their intrinsic properties but are approximated to have no influence on phase separation. Although transcription and translation resources such as nucleotide triphosphates and amino acids are not strictly dilute, they are treated as such for expediency in modeling. (3) The system is assumed to remain at phase equilibrium at all times (see Supplementary methods for more information section 11.2). We use the kinetic model to obtain the transcription *k_r_* and degradation *k_d_* rate constants that give the best fit to the experimental data, shown in Figure 3.

### Determination of transcription and degradation rate constants without condensates

In the case where mRNA and GdmS concentrations lie outside the binodal region (see Figure 2C), the system is composed of a single phase. The reaction scheme and reaction rates for the one-phase system is decribed in detail in Supplementary methods section 11.1. The optimal values for different parameters were obtained by globally fitting the model to mRNA and GdmS concentration profiles for 5 different transcription enzyme (T7) concentrations (470 nM, 970 nM, 1470 nM, 1720 nM and 2040 nM) and fourteen different DNA concentrations ranging from 0-30 nM (Figure 3, Supplementary methods section 12.4).

The translation process was modeled as conversion of TlR into GdmS with mRNA as the template and an initial TlR concentration [TlR]_t=0_, the translation rate being proportional to the translation rate constant *k_p_*, and the concentrations of mRNA, transcription and translation resources [mRNA], [TsR] and [TlR], respectively (Supplementary methods section 11.1). Finally, GdmS or GdmS* was not considered to be degradable in the experimental timescales (See Supplementary methods, section 11.3). Note, hereafter GdmS* is referred to as GdmS for simplicity. Fitting our model to mRNA and GdmS kinetic data gives values for *k_r_* (0.012 ± 0.005 nM^−2^*h*^−1^ ) and *k_d_*(0.022 ± 0.018 nM^−1^h^−1^); details see Supplementary methods sections 12.1, 12.2 and 12.3.

Since the condensate volume is a measurable output of the experimental system, we used phase diagrams to relate mRNA concentrations to the volume of the condensates. Experimentally measured phase boundaries and tie-lines for purified mRNA and GdmS in TNT extract (Figure 2C) were used as the basis for a constructed phase diagram (see Supplementary methods section 11.5). This gave information on the concentration of mRNA and GdmS, where phase separation will occur and the tie-lines give the volume fraction of the GdmS-rich phase.

### Determination of transcription and degradation rate constants in the presence of condensates

To determine how dynamic compartmentalization modulates transcription and degradation to produce measurable changes in the condensate volume, we estimated the transcription and degradation rates in the GdmS-poor (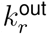 and 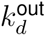) and GdmS-rich phases, (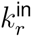 and 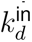) phases by fitting a two-phase kinetic model that explicitly incorporates phase separation of mRNA and GdmS to the experimental data. To improve the robustness of these estimates, we fitted the model to the condensate volume fraction as a function of time, measured by confocal microscopy, for GdmS doping concentrations of 4.5, 5, 6, 10, and 20 *µ*M without additional T7 or RNase A (Figure 3C1), and for concentrations of 5, 6, and 10 *µ*M with an additional 1 *µ*M T7 and 0.1 *µ*M RNase A (Figure 3C2). Results showed that increasing the initial GdmS concentration results in a higher maximum volume fraction of the GdmS-rich phase. This trend is consistent with the Lever rule applied to the phase diagram, which states that increasing the total concentration of the condensate components will lead to an increase in the condensate volume fraction (see Supplementary methods Section 11.5, Eq. (28)).

Given the mRNA–GdmS phase diagram and the experimentally determined condensate volume traces, we extracted the transcription and degradation rate constants in the GdmS-poor and GdmS-rich phases by fitting the condensate volume fraction dynamics using the two-phase kinetic model. We employed a fitting routine that iteratively solved the two-phase kinetic model for different values of the dilute- and GdmS-rich-phase transcription rate constants, 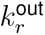 and 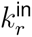, and degradation rate constants, 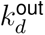 and 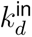, in order to match the volume traces generated by the model to the experimental data. The rate constants 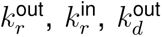, and 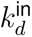 were empirically shown to depend exponentially on the doped GdmS concentration [GdmS]_add_, as described in the Supplementary methods section 11.6.

The fits obtained with the two-phase kinetic model provide estimates of the intrinsic degradation rate constants in the one-phase and two-phase regimes (Figure 3G,H), as well as the transcription rate constants before and after phase separation. However, the model could not reliably resolve distinct transcription rate constants for the GdmS-rich and GdmS-poor phases. This likely reflects the relatively small contribution of condensate-localised transcription to the overall transcription rate.

The plot of the intrinsic transcription and degradation rate constants as a function of doped GdmS concentration shows that crossing the phase boundary (at 4.5 *µ*M of GdmS) to a twophase region leads to a significant decrease in both the transcription and degradation rate constants. For the transcription rate constant there is a two order of magnitude decrease from the one phase region ((1.608 ± 0.639) *×* 10^−2^) nM^−2^h^−1^ to the two phase region ((1.487 ± 0.210) *×* 10^−4^) nM^−2^h^−1^ at 5 *µ*M GdmS. For the degradation rate constant, there is an order of magnitude decrease in the one phase region ((4.29 ± 0.05) *×* 10^−3^)nM^−1^h^−1^ to ((2.66 ± 1.05) *×* 10^−4^)nM^−1^h^−1^ in the two-phase region at a doping concentration of 5 *µ*M GdmS. The overall rate of degradation (*k_d_×* [RNase A] *×* [mRNA]) is reduced despite the upconcentration of RNase A inside the condensates.

Upon increasing the GdmS concentration to 20 *µ*M, we observed a further decrease in the effective rates of transcription and degradation in both the GdmS rich and GdmS poor phases. The transcription rate constant and decreases to *k_r_* = ((1.487 ± 0.210) *×* 10^−4^) nM^−2^h^−1^ and degradation rate constant, *k_d_*, to ((6.06 ± 6.05) *×* 10^−9^) nM^−1^h^−1^. These values are so low that they can be considered negligible. One explanation for this is that the condensate environment suppresses transcription and degradation by sequestering T7 and RNase A (See Supplementary methods section 12.5). With increasing volume fraction of condensates, more enzyme is partitioned into the condensate, almost completely attenuating transcription and degradation.

The observed dependence of the rate constants on GdmS concentration can be motivated using the theory of chemical kinetics for non-dilute mixtures, as explained in the reference [38] and discussed in Supplementary method section 11.6. While it is not possible to determine precisely how the condensate environment suppresses enzyme activity, the observed reduction could be attributed to factors including reduced diffusion length scales of enzymes, substrate, and co-factors (see Supplementary methods text), changes in enzyme–substrate binding constants, and/or inhibition of the enzyme active site. In addition, it cannot be ruled out that GdmS directly interferes with T7 and RNase A activity. Despite this, it is clear that the formation of condensates regulates enzyme kinetics in a manner that is commensurate with condensate dissolution. Given that transcription and degradation are suppressed within the condensate, condensate dissolution is likely driven predominantly by degradation in the GdmS-poor phase of the dispersion.

### Condensate dissolution driven by mRNA degradation in the GdmS-poor phase

To test the prediction that the decrease in condensate volume is driven by RNase A-mediated degradation in the GdmS-poor phase, we performed a series of control experiments to confirm that mRNA can indeed be degraded in the GdmS poor phase. First, we directly measured the effect of RNase A on condensate volume. The condensates were prepared in TNT extract by mixing 10 *µ*M GdmS and 100 nM mRNA. RNase A was added with increasing concentrations of 1, 2, 5, and 10 *µ*M. The volume fraction of the condensates was tracked as a function of time by confocal microscopy as previously described (see Figure 4B). The volume fractions were then plotted against the reduced time (*τ* = time *×* [RNase A]). These plots were overlaid and fit with a single exponential function (Supplementary methods section 13) to obtain the degradation rate *k*^avg^ = (1.50 ± 0.40) *×* 10^−3^ nM^−1^h^−1^. It is noted that this value is comparable to the intrinsic degradation rate constant for mRNA in the GdmS-poor phase at the same doped GdmS concentration (10 *µ*M), viz. *k*^out^ = (1.39 ± 0.31) *×* 10^−3^ nM^−1^h^−1^. This agreement helped validate the degradation rates obtained from the two-phase model.

**Figure 4:**
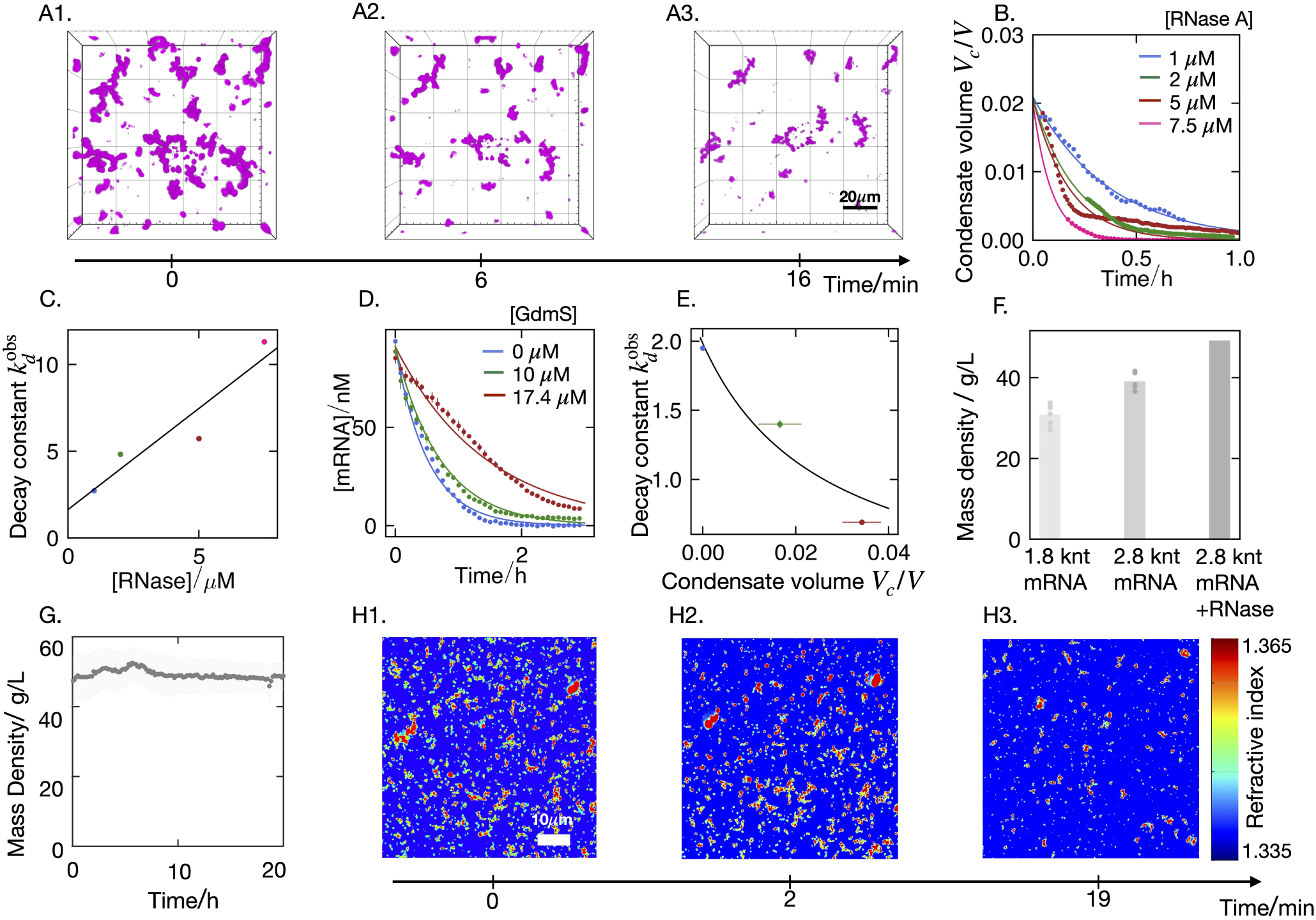
RNase activity is negatively correlated with condensate volume. (A1-A3) Example 3D confocal time-lapse showing the degradation of condensates formed from a mixture of 10 *µ*M GdmS,0.1 *µ*M mRNA, and 5 *µ*M RNase A. (B) Plots showing the change in condensate volume fraction over time for 10 *µ*M GdmS and 0.1 *µ*M mRNA at different RNase A concentrations: 1 *µ*M (blue), 2 *µ*M (green), 5 *µ*M (red), and 7.5 *µ*M (pink), obtained from analysis of 3D confocal timelapses. (C) The decay constants obtained by fitting the time traces in (B) to a first-order decay model plotted as a function of RNase A concentration, fit to a pseudo-first order model. (D) Time traces report mRNA degradation in the isolated GdmS-poor phase after condensate removal by centrifugation, with an additional 0.10 *µ*M mRNA and 20 *µ*M DFHBI added. The removed condensates were originally prepared with 0.10 *µ*M mRNA and either 0 *µ*M GdmS (blue circles), 10 *µ*M GdmS (green circles), or 17.4 *µ*M GdmS (red circles). Error bars indicate experimental triplicates. Solid lines show fits of the data to a first-order decay model to obtain the observed decay constants, 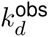. (E) 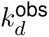 was plot against the volume fraction of the condensates. which were obtained by analysis of 3D confocal micrographs of the condensates as described above fit to a model explaining the correlation between condensation and suppression of decay rate constants (Supplementary section 2.8). (F) Bar chart showing the mass density inside condensates prepared by mixing 10 *µ*M GdmS with 0.15 *µ*M 1.8 knt mRNA, 0.10 *µ*M 2.8 knt mRNA, and the time averaged mass density of condensates prepared by mixing 10 *µ*M GdmS, 0.10 *µ*M 2.8 knt mRNA, and 0.10 *µ*M RNase A, obtained from holotomographic microscopy (G) Plot of the mass density of the latter condensates as a function of time shows that the density inside the condensates remains constant. (H1-H3) Snapshots of 2D holotomographic micrographs from a representative time-resolved experiment showing volume and refractive index changes for the same condensates. The images reveal a decrease in condensate area (and therefore, volume fraction) with no detectable change in refractive index. The color bar indicates the refractive index scale. All condensates were prepared in 80% TNT extract, and experiments were performed at 30*^◦^*C.

**Figure 5:**
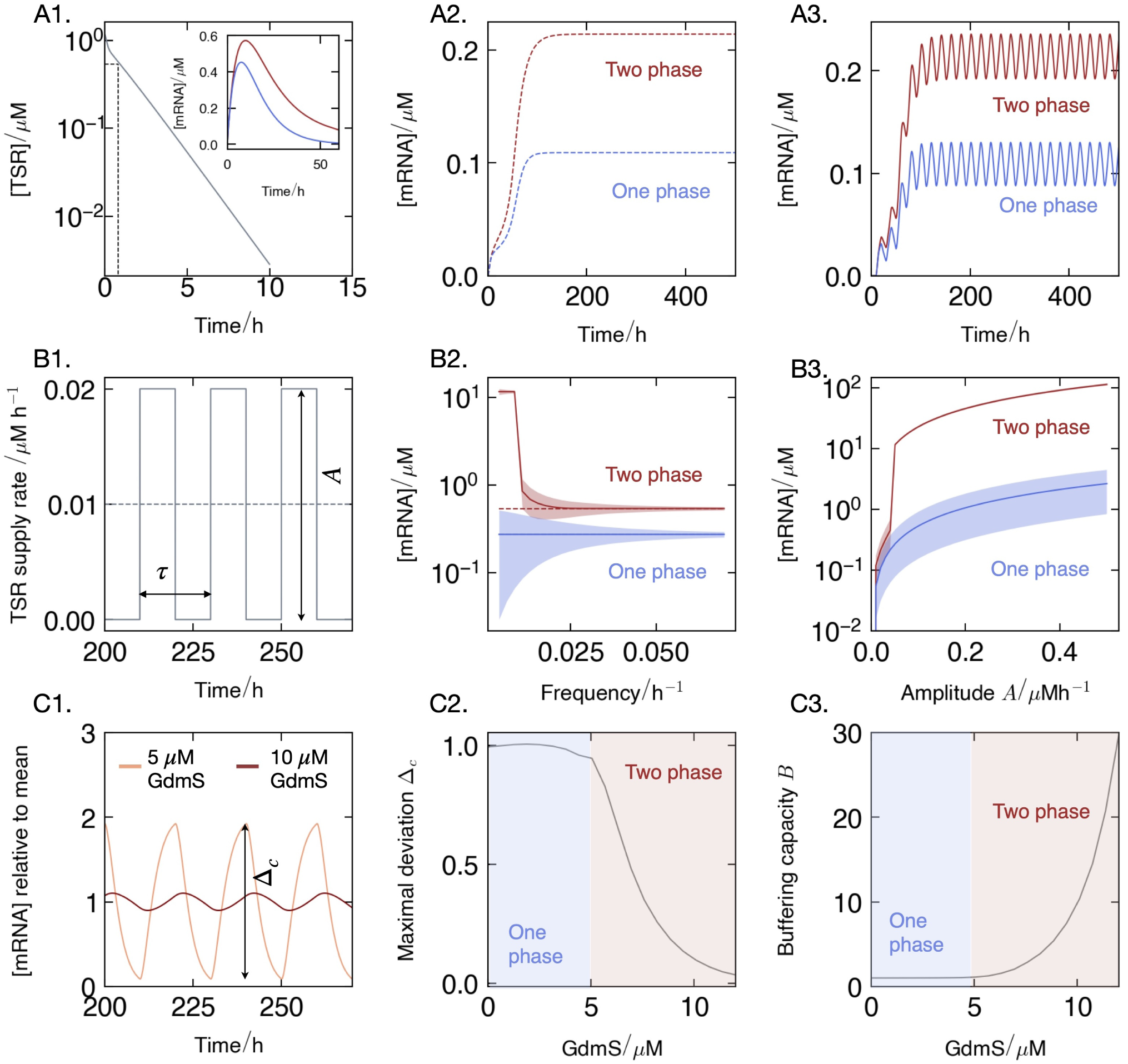
Phase separation and oscillatory supply of TsR enhances mRNA concentration: (A) Time evolution of mRNA concentration in the one-phase (blue) and two-phase scenarios (red) depending on TsR supply. While for a finite initial amount of TsR (A1), both mRNA and TsR decay in time (inset of A1), a constant supply of TsR (A2) leads to a stationary state in mRNA levels. (A3) When oscillating the TsR supply with frequency 1*/τ* and amplitude *A*, mRNA shows oscillations as well. The comparison between constant (dotted line in B1) and oscillating supply of TsR (solid line in B1) shows a higher mRNA concentration for oscillatory supply (solid red line in B2) compared to the constant case in the two-phase system (dotted red line in B2), particularly for low frequencies. In (B3), we see that upon varying the supply amplitude *A* of TsR at constant supply frequency 1*/τ* = 0.01 h^−1^, the two-phase system gives a higher mRNA concentration, which increases non-linearly with *A*. (C) Deviations in mRNA concentration are smaller for a higher GdmS concentration. (C1) shows the mRNA concentration relative to a mean value (relative amplitude Δ_mRNA_) calculated for [GdmS]=10 *µ*M (dark curve) and at [GdmS]=5 *µ*M (light curve) at an average TsR supply rate of 0.01 *µ*M h^−1^, oscillating at a frequency 1*/τ* =0.05 h^−1^, with an amplitude *A* = 0.005*µ*Mh^−1^. (C2) The maximal deviation of mRNA relative to the mean RNA concentration, Δ_mRNA_, decays with GdmS concentration. The decay is more pronounced in the two-phase region (red-shaded area) than in the one-phase region (blue-shaded area), showing that condensates buffer mRNA levels. (C3) The increase of the buffering capacity *B* (Eq. (2)) with [GdmS] confirms the condensate-mediated buffering effect.

Next, given that RNase A was known to partition into the condensate, the presence and activity of RNase in the GdmS-poor phase were examined. Three different dispersions of condensates in TNT extract were prepared: 0.10 *µ*M mRNA and 0.10 *µ*M RNase A with either 0, 10, or 17.4 *µ*M GdmS (see Materials and Methods and Supplementary methods section 14.1). The condensates were centrifuged to pellet the GdmS-rich phase. For each solution, the GdmS-poor phase was removed conservatively to prevent disturbance of the pellet, and a further 0.10 *µ*M mRNA and 20 *µ*M DFHBI dye were added to the GdmS-poor phase. Kinetic data shows degradation of mRNA in the GdmS-poor phase for each of the samples, and this was taken to confirm that sufficient RNase A was present in the GdmS-poor phase to drive mRNA degradation. To further corroborate this behaviour, a first-order decay curve was fit to the decrease in mRNA concentration to obtain the pseudo-first-order observed degradation rate constant (*k*^obs^). This rate constant was then plotted against condensate volume fraction *V_f_* , and a strong negative correlation was observed as depicted in Figure 4D,E. This behaviour indicates that an increased volume fraction of the GdmS-rich phase is associated with suppression of mRNA degradation and suggests a protection mechanism for mRNA upon condensation (for further discussion, please see Supplementary Material section 11.6 and section 14.1). The intercept of this fit Eq. (60) gave a predicted intrinsic degradation rate constant of = (5.33 ± 0.01) *×* 10^−3^ nM^−1^h^−1^. This value is comparable to the degradation rate constant of (4.29 ± 0.05) *×* 10^−3^ nM^−1^h^−1^ obtained by fitting with the one-phase kinetic model (Figure 3) that confirms that estimates from mass action models provide reasonable values.

To directly assess any change in condensate density arising from mRNA degradation, holotomographic microscopy was used to measure the density within condensates during mRNA decay. Holotomographic microscopy reports local mass density by quantifying refractive index variations on a pixel-by-pixel basis and converting these to density via calibration (Supplementary Methods section 15). To confirm that mRNA degradation and the accumulation of shorter mRNA fragments within condensates would produce a measurable decrease in refractive index, and thus mass density, condensates were prepared with GdmS and either 1.8-kilonucleotide (knt) mRNA or 2.8-knt mRNA (the full-length mRNA encoding GdmS) at equivalent total charge (0.15 *µ*M and 0.10 *µ*M respectively) (see Supplementary methods section 15).

These results show that condensates formed from 1.8 knt mRNA have a lower mass density of 30.90 ± 1.041 gL^−1^ compared to the full length mRNA mass density of 39.12 ± 1.091 gL^−1^ (see Figure 4F). This indicates that shorter length mRNA within condensates would show a measurable reduction in condensate density. Thus, we tracked the change in density of the condensates prepared with 10 *µ*M GdmS, 0.1 *µ*M of mRNA in the presence of 5 *µ*M RNase A over 20 minutes (Figure 4(G-H)). Time-dependent holotomographic imaging showed a constant mass density of 49.17 ± 5.020 gL^−1^ over 20 mins despite the complete loss of visible condensate volume (Figure 4(G)). This could be attributed to a negligible mRNA degradation rate constant in the condensate phase. However, it cannot be entirely ruled out that mRNA degradation still takes place within the condensate. For example, a constant mass density within the droplets could alternatively be maintained by the diffusive efflux of degraded mRNA balanced by the influx of mRNA across the interface. Nevertheless, within the resolution of the differences in refractive index detected by the microscope, we were unable to detect any change in the mass density of the dissolving condensate. Together, our results confirm that degradation can take place in the GdmS-poor phase, which provides some evidence that condensate dissolution could be driven by mRNA degradation in the GdmS poor phase.

### Enhanced protection of mRNA against degradation when supplying transcription resources

We have developed a kinetic model governing the formation and dissolution of condensates, which is driven by transcription and degradation of mRNA in a cell-free expression system. A key feature of the cell-free expression system is that it contains a finite amount of resources. Consequently, during the time course of mRNA production, the resources TsR are consumed, tending towards zero (Figure 5A1). Numerical solution to our model of TsR shows a decay in the resources from 1000 to 500 nM in about 50 minutes for the case of 470 nM T7 polymerase. Once the mRNA concentration exceeds the mRNA saturation concentration, condensates form that grow in volume over time (Figure 1). After one hour, once TsR is sufficiently depleted, the degradation kinetics dominates, leading to a decay of mRNA and thus shrinkage of condensates (see inset of Figure 5A1). As mRNA levels drop below the mRNA saturation concentration (for fixed GdmS), condensates dissolve.

In cellular systems, transcriptional resources, including nucleotide triphosphate levels (NTPs), are either constant or fluctuate in time [39, 40]. Thus, a biologically relevant case is to study our model of the cell-free expression system for conditions where resources are supplied. Specifically, it is interesting to distinguish between constant and oscillatory supplies that mimic a fluctuating environment. In the following, we use our two-phase model with the parameters obtained from the experimental closed-cell free expression systems to study the effect of condensate feedback in an open system where the resources are under constant supply or input with an oscillatory input with frequency 1*/τ* and an amplitude *A*; see Figure 5B1. The GdmS concentration is fixed to 10 *µ*M, and our kinetic model was set to consider both (i) a two-phase (phase-separated system) and (ii) a one-phase (homogeneous) system. The latter is analogous to a system where nucleation is continually suppressed, preventing condensate formation and maintaining a supersaturated homogeneous phase (see Supplementary methods section 16).

Numerical solutions to the kinetic model show that when resource supply is constant, mRNA concentration increases and reaches a steady state. This steady state reflects that mRNA transcription and degradation, in both the one and two-phase systems, are balanced. When TsR is supplied in an oscillatory manner, mRNA concentration increases at early time points. At long times, mRNA concentration oscillates around a mean value. Such oscillations result from a competition between transcription and degradation. In each TsR supply cycle (Figure 5B1), the mRNA transcription rate eventually exceeds the degradation rate, leading to an increase in the mRNA concentration. In the second half of the cycle, when no TsR is supplied, degradation eventually dominates over transcription, making mRNA levels decay over time. Over many TsR supply cycles, the competition between transcription and degradation gives rise to smooth, sinusoidal-like oscillations in mRNA concentration.

Interestingly, in both cases where the TsR resources are supplied as a constant or pulsed input, we observe that phase separation (two-phase system) leads to a higher mRNA concentration than in the homogeneous (one-phase) system (Figure 5A2,A3). This finding holds across all TsR oscillation frequencies and amplitudes (Figure 5B2,B3). Since the degradation rate constant is lower in the condensates than in the protein-poor phase, the mRNA is effectively protected within the condensates. The result is a higher total mRNA concentration in the presence of condensates compared to the one-phase system.

The supply frequency is an important parameter that regulates mRNA levels in the system. We find that reducing the supply frequency 1*/τ* in the two-phase system causes a significant deviation in the average mRNA concentration (red solid line) from that in the constant supply case (red dotted line); see Figure 5B2. A longer feeding duration *τ* , (low frequency) provides more resources to the system compared to high frequency input to produce more mRNA. A higher mRNA concentration corresponds to a larger condensate volume fraction, which is already reflected in the phase diagram (Figure 1). Such larger condensates can protect more mRNA from degradation, showing that oscillatory TsR supply provides an even stronger protection mechanism. Remarkably, below a certain frequency, the condensates start occupying most of the system volume, leading to a rapid increase in mRNA concentration with decreasing frequency. This increase saturates because the condensates then occupy the entire system volume at all times, making the mRNA concentration independent of the oscillation frequency below 1*/τ* = 0.008 h^−1^. In addition, lowering the frequency results in larger oscillation amplitudes of mRNA concentrations.

We next studied the effect of the supply amplitude *A* on the phase-averaged mRNA concentration [mRNA]^avg^. In this case, we fixed the TsR supply oscillation frequency to 1*/τ* = 0.01 h^−1^ and varied the supply amplitude, *A*, for the one and two-phase systems. With increasing supply amplitude, *A*, we observe an increase in average mRNA concentration that is commensurate with larger TsR levels leading to more mRNA. For the one-phase system, the mRNA concentration linearly increases with oscillation amplitude. In contrast, the two-phase system shows a non-linear increase, particularly for larger TsR supply amplitude, *A*. This non-linear trend results from the condensates slowing down the degradation of mRNA compared to the one-phase system. Moreover, above a certain supply amplitude, we see a rapid increase in mRNA concentration, see Figure 5B3. This is because above this supply amplitude, the condensates occupy most of the system volume leading to a drastic slow down of degradation. Once condensates occupy the entire system volume, the mRNA concentration continues to increase linearly with *A*. This is expected because the condensate volume cannot change any further, so the increase in mRNA concentration is solely due to increase in TsR. Taken together, these results show that when more resources are provided by a smaller oscillation frequency 1*/τ* or a larger TsR resource amplitude, *A*, mRNA concentration as well as the condensate volume increase. In other words, since condensates protect mRNA from degradation, they provide positive feedback on mRNA concentration.

To quantify this feedback, we consider the mRNA concentration relative to the average concentration. Figure 5C1 shows this quantity for 5 *µ*M and 10 *µ*M GdmS as a function of time. We see that larger protein concentration leads to smaller deviations from the mean, and thus a smaller maximal deviation of mRNA concentration (Figure 5C1,C2). This maximal deviation is defined as:

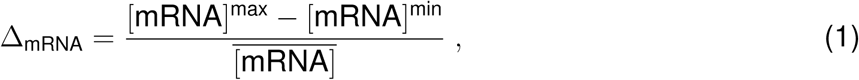

where [mRNA]^max^, [mRNA]^min^ and [mRNA] are the maximum, minimum and time-averaged mRNA concentrations during a cycle, respectively. Moreover, the Δ_mRNA_ is much lower in the phase-separated compared to the one-phase system (figure 5C2) . This trend suggests that phase separation can reduce overall mRNA oscillations. In other words, condensates buffer mRNA variations against changes in TsR concentrations, analogous to the classical buffering capacity of salts and amphoteric compounds upon acid-base titration. The propensity of condensates to buffer mRNA variations can be characterized by the buffering capacity

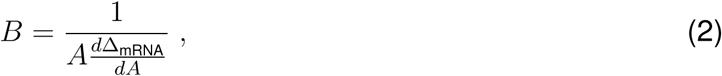

where *A* is the amplitude of TsR oscillations (Figure 5B1). According to Eq. (2), a large buffering capacity *B* corresponds to minimal changes of the deviations in mRNA concentration, Δ_mRNA_, upon changes in the TsR oscillation amplitude *A*.

Our model shows that the buffering capacity *B* is much higher in the two-phase region compared to the one-phase region. Upon phase separation, the buffering capacity increases nonlinearly with GdmS protein concentration [GdmS] (Figure 5C3), and thus, the volume of the condensates. This non-linearity results from condensates providing a feedback mechanism on mRNA degradation.

Taken together, our kinetic model confirms that a phase-separated system exhibits additional properties compared to a homogeneous or one-phase system. In this instance, condensates suppress mRNA degradation, thereby protecting mRNA, leading to an overall higher concentration of mRNA in a two-phase system compared to a one-phase system. Condensates can be sustained by periodically supplying resources to the system. Due to this supply of transcriptional resources, mRNA concentration oscillates, resulting from a competition between transcription and degradation. Further increasing the average amount of the system’s resources, either by lowering the oscillation frequency of the supply or increasing the input amount (oscillation amplitude), leads to higher mRNA concentrations and thus a higher condensate volume. This enhanced condensed volume can protect more mRNA, corresponding to a positive feedback loop. Finally, we show that condensates can buffer the relative mRNA concentration in a condensate-dependent manner. Overall, our results point to a significant protective mechanism of condensates, which can also buffer deviations to mRNA concentration, driven by a positive feedback between condensate volume and mRNA concentration.

## Conclusion

Kinetic modeling with minimal cell-free systems offers a unique opportunity to explore emergent properties arising from coupling enzyme reactions with compartmentalization. We show that in vitro mRNA production drives in situ phase separation with a mutant form of G3BP1, where condensates modulate transcription and degradation rate constants. This creates a self-regulating, phase-separating system evidenced by the spontaneous formation and dissolution of synthetic condensates.

Our kinetic model quantifies how compartmentalization governs its own dynamics by estimating transcription and degradation rate constants inside and outside droplets. This can be extended to any phase-separating system, including lipid membranes. Our model shows that phase separation suppresses both rate constants, with degradation occurring more slowly within droplets, suggesting that droplet dissolution is driven by mRNA decay in the protein-poor phase. The molecular basis of enzyme suppression remains unclear and is difficult to elucidate, as they likely involve multiple contributing factors. For example, it may involve interactions between GdmS and T7 RNA polymerase or RNase A that reduce activity; or partitioning of enzymes into condensates that lower effective concentrations outside droplets and therefore the overall rate. Additionally, charged, crowded, and viscous environments can affect rate constants by affecting molecular diffusion (see SI and references). Indeed, control and engineering of condensate material properties could offer a rational route to modulating enzymatic reactions in cross-regulatory systems [41, 42, 43, 44]

Our findings highlight the functional role of compartmentalization in dynamic, resource-dependent systems where enzyme suppression enables droplet regulation, molecular protection, and buffering (see Fig. 5). It is plausible that compartments can provide a general cross-regulatory role within reaction networks that influence different functional outcomes. For example, compartments accelerate reaction rates, creating a positive feedback mechanism, or the correct balance of degradation and formation could stabilise droplet size. In addition, self-regulatory feedback mechanisms can be driven by other reaction networks such metabolism where it has been shown that small molecule metabolites, such as NADPH, ATP can drive compartment formation and affect material properties [45, 46, 47, 48]. Given that condensates selectively enrich certain mRNAs [49, 50, 51], modulation of enzyme reactions hint towards an evolutionary selection pressure for the protection of specific transcripts in a prebiotic test-tube setting. Finally, our work positions condensates as central hubs for coordinating enzymatic networks and emergent dynamic cellular functions and paves the way for the design of minimal life-like systems with sustained out-of-equilibrium behaviour.

## Methods

Details regarding the materials, experimental methods and procedures can be found in supplemental methods.

## Resource availability

### Lead contact

Requests for further information and resources should be directed to Christoph Weber (christoph.weber@ augsburg.de) or Dora Tang

### Materials availability

Plasmids generated in this study have been deposited to [Addgene, name and catalog number or unique identifier].

## Data and code availability

## Acknowledgments

Funding by the European Union (ERC, MinSynCell, 101088834, T-YDT) and (ITN, DARCHEMDN Grant number 101119956, T-YDT). Views and opinions expressed are, however, those of the author(s) only and do not necessarily reflect those of the European Union or the European Research Council; neither the European Union nor the granting authority can be held responsible for them. We acknowledge financial support from the University of Saarland and the Max Planck Society (T-Y,D.T.). A.G. was funded by the ELBE post-doc programme from the MPI-PKS; A.G. and S.S. was funded through ERC MinSynCell, 101088834. We thank the PharmaScienceHub for financial support (T-Y,D.T.).

We thank the Light Microscopy Facility (LMF), Advanced Imaging Facility (AIF), and Protein expression, purification and characterisation Facility (PEPC) of the Max Planck Institute of Molecular Cell Biology and Genetics (MPI-CBG) for their technical support and useful discussions. We thank in particular Jan Peychl of the LMF for use of the holotomography microscopy and Michael Bugiel of the AIF for help with writing the pMOT analysis code. We further thank the light microscopy core facility at the University of Saarland. We thank the Hyman Lab for purified proteins and the Alberti Lab for protein plasmid. We thank Ivar Haugerud, Gaetano Granatelli and Samuel Gomez for fruitful discussions on the model. C.A.W. acknowledges the European Research Council (ERC) for financial support under the European Union’s Horizon 2020 research and innovation programme (“Fuelled Life”, Grant agreement No. 949021).

## Author contributions

T-Y.D.T, A.G., A.T., C.W. contributed to the design and concept of the study. R.R., A.G., S.S., A.T., C.W., T-Y.D.T contributed to experimental design, methodology and formal analysis; R.R. A.G. S.S. undertook the experiments; A.G. A.T and C.W. performed the theoretical calculations. A.G., A.T., C.W., T-Y D.T contributed to the writing of the manuscript. T-Y D.T and C.W. acquired funding.

## Declaration of generative AI and AI-assisted technologies

During the preparation of this work, the author(s) used Copilot/Grammarly in order to polish text and ChatGPT enhancing the efficiency of coding. After using this tool or service, the author(s) reviewed and edited the content as needed and take(s) full responsibility for the content of the publication.

