## supplementary information for "Enzyme-driven phase separation of synthetic condensates enables self-organizing compartments and protective microenvironments"

### Contents

**1 List of materials** 2

**2 mRNA and GdmS sequences** 3

**3 Plasmid Design and preparation** 5

**4 Transcription and purification of mRNA** 6

**5 Expression and purification of GdmS** 7

**6 Fluorescent labeling of T7 RNA polymerase and RNase A** 8

**7 Confocal microscopy** 9

**8 Construction of Phase Diagrams for mRNA-GdmS Condensates** 9

**15 Refractive Index Imaging of Condensate Degradation by Holotomography 35**

**16 Transcription and degradation kinetics with unlimited transcriptional resource (TsR) supply 36**

**17 Supplementary note: Determination of GdmS-mRNA condensate material properties using optical tweezer based passive microrheology 38**

**18 Supplementary note: Diffusion coefficient determination by Fluorescence Recovery after Photobleaching (FRAP) 41**

### 1 List of materials

| Reagent | Supplier | Catalogue no. |
|---|---|---|
| Plasmids for protein and mRNA expression | MPI-CBG, Dresden (Germany) | in-house |
| T7 RNA polymerase | Protein Expression and Purification (PEPC) facility, MPI-CBG (Germany) | in-house |
| H3C protease | Protein Expression and Purification (PEPC) facility, MPI-CBG (Germany) | in-house |
| FUS-GFP protein | Hyman laboratory, MPI-CBG (Germany) | gift (no catalogue no.) |
| MBPTrap HP column, 5 mL | Cytiva (Germany) | 28-9187-79 |
| HiLoad 16/600 Superdex 200 pg column | Cytiva (Germany) | 28-9893-35 |
| Amicon Ultra-15 centrifugal filter unit, Ultracel-PL, 30 kDa | Merck-Millipore (Germany) | UFC903024 |
| Amicon Ultra-4 centrifugal filter unit, Ultracel-PL, 30 kDa | Merck-Millipore (Germany) | UFC803024 |
| Amicon Ultra-0.5 centrifugal filter unit, Ultracel-PL, 30 kDa | Merck-Millipore (Germany) | UFC500324 |
| Microscope slides | Paul Marienfeld GmbH & Co. KG (Germany) | 1000000 |

| Reagent | Supplier | Catalogue no. |
|---|---|---|
| Cover slips, 24 × 60 mm, No. 1.5H | Paul Marienfeld GmbH & Co. KG (Germany) | 0107242 |
| Cover slips, 24 × 24 mm, No. 1.5 | Paul Marienfeld GmbH & Co. KG (Germany) | 0102062 |
| Flat-bottom 384-well polypropylene microplates | Greiner Bio-One (Germany) | 781209 |
| $\mu$-Slide 8-well chambered coverslips | Ibidi GmbH (Germany) | 80826 |
| Anti-evaporation silicone oil, 125 mL | Ibidi GmbH (Germany) | 50051 |
| Double-sided adhesive tape, 19 mm × 33 m | Scotch (3M Deutschland GmbH, Germany) | 665 (3M ID 7100169984) |
| Poly(ethylene glycol) (PEG) silane | abcr GmbH (Germany) | AB111226 |
| NruI restriction enzyme | New England Biolabs (USA) | R3192 |
| HiScribe T7 High Yield RNA Synthesis Kit | New England Biolabs (USA) | E2040S |
| QIAquick PCR Purification Kit | QIAGEN GmbH (Germany) | 28104 |
| RNeasy Kit | QIAGEN GmbH (Germany) | 74134 |
| RNase-free water | Merck/Sigma-Aldrich (Germany) | 7732-18-5 |
| RNase A | Merck/Sigma-Aldrich (Germany) | R4875 |
| DFHB1 ((Z)-4-(3,5-difluoro-4-hydroxybenzylidene)-1,2-dimethyl-1H-imidazol-5(4H)-one) | Merck/Sigma-Aldrich (Germany) | SML1627-5MG |
| ATTO 488 NHS ester | Merck/Sigma-Aldrich (Germany) | NC1868714 |
| Alexa Fluor 680 NHS ester | Merck/Sigma-Aldrich (Germany) | A20008 |
| Poly-L-lysine hydrobromide (150–300 kDa) | Merck/Sigma-Aldrich (Germany) | P4832 |
| TnT T7 Insect Cell Extract Protein Expression System | Promega (USA) | L1101 |
| Tris-HCl | Merck/Sigma-Aldrich (Germany) | T5941 |
| NaCl | Avantor/VWR (USA) | 27810.295 |
| DTT | Thermo Scientific (USA) | R0862 |
| Glycerol | Avantor/VWR (USA) | 24388.295 |
| Arginine-HCl | Merck/Sigma-Aldrich (Germany) | A5131 |
| PBS | Merck/Sigma-Aldrich (Germany) | P3813 |
| Maltose | Merck/Sigma-Aldrich (Germany) | M5885 |
| Phosphate buffer component ($Na_2HPO_4$, dibasic, anhydrous) | Grüssing GmbH (Germany) | 121471000 |
| Phosphate buffer component ($NaH_2PO_4$, monobasic, anhydrous) | Thermo Scientific (USA) | 389870025 |

Supplementary table 1: Reagents and materials

### 2 mRNA and GdmS sequences

mRNA sequence:

AUGGUGAGCAAAGGGGAAGCAGUGAUCAAGGAGUUUAUGAGGUUCAAGGUCCA
CAUGGAGGGGUCAAUGAAUGGGCACGAGUUCGAAAUCGAGGGAGAGGGAGAGG
GCAGGCCCUACGAGGGCACCCAGACAGCCAAGCUGAAGGUGACCAAGGGAGGA
CCACUGCCCUUCAGCUGGGACAUCCUGUCCCCUCAGUUCAUGUAUGGCUCUCG

GGCCUUUAUCAAGCACCCUGCCGACAUCCCAGAUUACUAUAAGCAGAGCUUCC
CAGAGGGCUUUAAGUGGGAGAGAGUGAUGAACUUCGAGGACGGAGGAGCAGUG
ACCGUGACACAGGACACCUCCCUGGAGGAUGGCACACUGAUCUACAAGGUGAA
GCUGAGGGGCACAAAUUUCCCCCUGAUGGCCCAGUGAUGCAGAAGAAGACAA
UGGGCUGGGAGGCCUCUACAGAGCGCCUGUAUCCCGAGGACGGCGUGCUGAAG
GGCGAUAUCAAGAUGGCACUGCGGCUGAAGGACGGCGGCAGAUACCUGGCCGA
CUUCAAGACCACAUAUAAGGCCAAGAAGCCCGUGCAGAUGCCUGGCGCCUACA
ACGUGGACAGAAAGCUGGAUAUCACCAGCCACAAUGAGGAUUAUACAGUGGUG
GAGCAGUAUGAGAGGUCCGAAGGGAGACAUUCAACCGGGGGGAUGGAUGAGCU
GUAUAAG GGAUCCGCUGGCUCCGCUGCUGGUUCUGGCGCGGCCGCG
AUGGUCAUGGAAAAACCCUCGCCUUUGCUCGUCGGAAGAGAGUUCGUGCGUCA
AUACUACACACUGCUCAACCAAGCCCCAGAUAUGCUGCACCGUUUCUACGGCA
AGAACUCCAGCUACGUGCACGGUGGCCUGGACUCUAACGGCAAGCCAGCUGAC
GCCGUCUACGGACAGAAGGAGAUCCACCGUAAGGUCAUGUCACAGAACUUCAC
CAACUGCCACACUAAGAUCAGGCACGUCGACGCUCACGCCACUCUGAACGACG
GAGUGGUCGUGCAGGUCAUGGGUCUGCUGUCCAACAACAACCAGGCUCUGCGC
CGUUUCAUGCAGACCUUCGUGCUGGCUCCUGAGGGAAGCGUCGCCAACAAGUU
CUACGUGCACAACGACAUCUUCCGUUACCAGGACGAAGUCUUCGGAGGUUUCG
UGACUAGUAGGCAGCAGACUCCCGAGGUCGUGCCAGACGACUCUGGUACCUUC
UACGACCAGGCUGUCGUGUCAAACGCUAGCGCCCAGAAGUCUUCAUCCCCAGC
UCCUGCCGACAUCGCUCAGACCGUCCAGGAGGACCUGAGGACUUUCAGCUGGG
CUUCUGUGACCUCAAAGAACCUGCCUCCCUCUGGUGCUGUCCCAGUGACUGGA
AUCCCACCUCACGUCGUGAAGGUCCCUGCCAGCCAGCCCAGACCAGAAUCCAA
GCCAGAGAGCCAGAUCCCUCCUCAGCGCCCUCAGAGAGACCAGAGGGUGAGAG
AACAACGCAUCAACAUCCCUCCCCAGAGGGGACCUAGGCCAAUCAGGGAAGCC
GGCGAGCAGGGAGACAUCGAGCCUAGGAGAAUGGUCCGCCACCCCGACUCUCA
CCAGCUGUUCAUCGGAAACCUGCCUCACGAAGUGGACAAGUCUGAGCUGAAGG
ACUUCUUCCAGUCAUACGGAAACGUCGUGGAACUGAGAAUCAACUCAGGCGGA
AAGCUGCCCAACUUCGGUUUCGUCGUGUUCGACGACUCCGAGCCCGUCCAGAA
GGUGCUGAGCAACCGUCCAAUCAUGUUCCGUGGUGAAGUCAGGCUGAACGUGG
AAGAGAAGAAGACCAGAGCUGCUAGGGAGGGUGACAGGAGGGACAACAGGCUG
AGGGGUCCAGGUGGCCCUAGAGGAGGUCUGGGCGGAGGUAUGCGCGGCCCACC
UCGUGGUGGUAUGGUCCAGAAACCCGGAUUCGGAGUCGGUAGAGGAUUGGCCC
CCAGACAA GGCGCGCCGUAAUAG
UUGCCAUGUGUAUGUGGGAGACGGUCGGGUCCAUCUGAGACGGUCGGGUCCAG
AUAUUCGUAUCUGUCGAGUAGAGUGUGGGCUCAGAUGUCGAGUAGAGUGUGGG
CUCCCACAUACUCUGAUGAUCCAGACGGUCGGGUCCAUCUGAGACGGUCGGGU
CCAGAUAUUCGUAUCUGUCGAGUAGAGUGUGGGCUCAGAUGUCGAGUAGAGUG
UGGGCUGGAUCAUUCAUGGCAA

Legend: mScarlet-I, Linker, G3BP1(ΔE1/ΔE2), 2× Broccoli aptamer.

GdmS sequence:

MVSKGEAVIKEFMRFKVHMEGSMNGHEFEIEGEGEGRPYEGTQTAKLKVTKGG
PLPFSWDILSPQFMYGSRAFIKHPADIPDYYKQSFPEGFKWERVMNFEDGGAVT
VTQDTSLEDGTLIYKVKLRGTNFPPDGPVMQKKTMGWEASTERLYPEDGVLKG
DIKMALRLKDGGRYLADFKTTYKAKKPVQMPGAYNVDRKLDITSHNEDYTVVEQ
YERSEGRHSTGGMDELYKG SAGSAAGSGAAA MVMEKPSPLLVGREFVRQYY
TLLNQAPDMLHRFYGKNSSYVHGGLDSNGKPADAVYGQKEIHRKVMSQNFTN

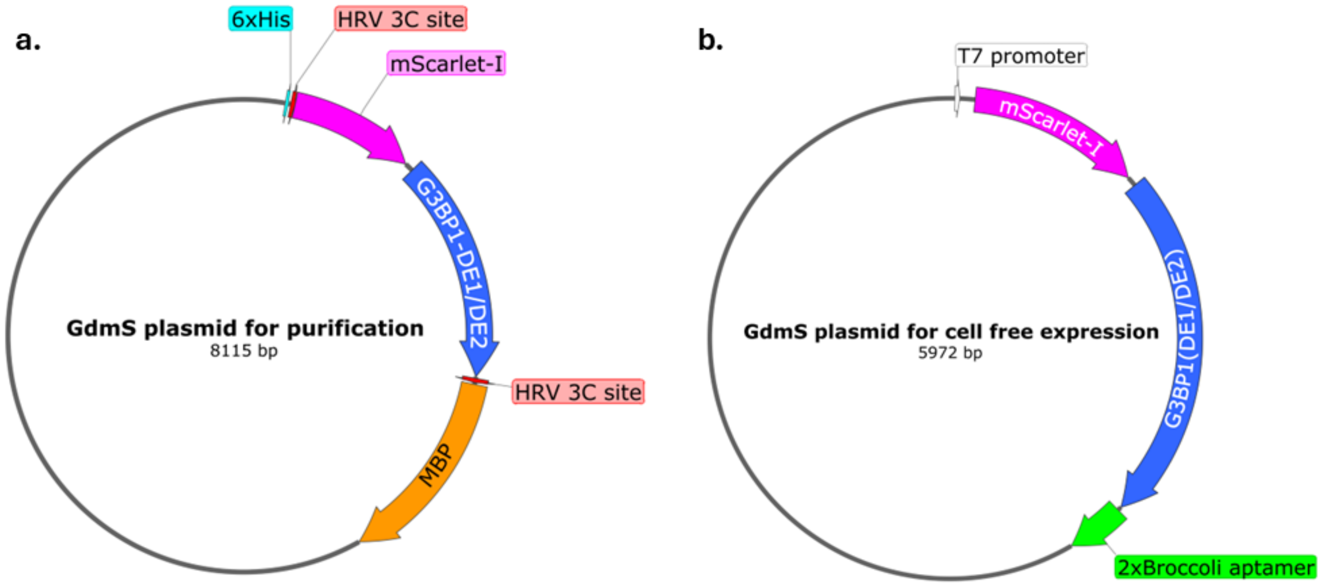

Supplementary figure 1: Plasmid constructs for a) protein expression in and purification from *T.ni.* cells and b) cell-free expression in TNT insect cell extract.

CHTKIRHVDAHATLNDGVVVQVMGLLSNNNQALRRFMQTFVLAPEGSVANKFY
VHNDIFRYQDEVFGGFV TsRQQTPEVVPDDSGTFYDQAVVSNASAQKSSSPAP
ADIAQTVQEDLRTFSWASVTSKNLPPSGAVPVTGIPPHVVKVPASQPRPESKPE
SQIPPQRPQRDQRVREQRINIPPQRGPRPIREAGEQGDIEPRRMVRHPDSHQLF
IGNLPHEVDKSELKDFFQSYGNVVELRINSGGKLPNFGFVVFDDSEPVQKVLSNR
PIMFRGEVRLNVEEKKTRAAREGDRRDNRLRGPGGPRGGLGGGMRGPPRGG
MVQKPGFGVGRGLAPRQGAP

Legend: mScarlet-I, Linker, NTF2L domain, Intrinsically disordered region.

### 3 Plasmid Design and preparation

A pOCC plasmid backbone was obtained from the Protein Expression and Purification (PEPC) Facility at the Max Planck Institute for Molecular Cell Biology and Genetics (MPI-CBG) that was optimized for expression in the TNT extract. The pOCC vectors were modified to contain AscI (5'-GG/CGCGCC-3') and NotI (5'-GC/GGCCGC-3') enzymatic recognition sites downstream of the T7 promoter for modular restriction cloning with various inserts. The vector contained the sequence for the mScarlet-I fluorescent tag upsteam to the cloning site and two consecutive Broccoli mRNA aptamers (as used in [1]). The gene used in this project was the deletion mutant of the IDP G3BP1, referred to as G3BP1(ΔE1/ΔE2) or Gd. This was sourced from the vector pOCC189-G3BP1(ΔE1/ΔE2)-L-667 (gift from Alberti lab). Cloning via restriction enzyme digestion of the G3BP1(ΔE1/ΔE2) gene using the NotI and AscI enzymes and ligation with T4 DNA ligase produced the cell-free expression ready plasmid pOCC499-mScarlet-I-G3BP1(ΔE1/ΔE2) expressing mScarlet-I-G3BP1(ΔE1/ΔE2), or GdmS.

Additionally, for purification of GdmS, the entire sequence of the fusion construct was cloned into a pOCC vector optimised for Baculovirus based infection and expression in Trichoplusia ni (*T.ni*) cells. The final construct had an hexa-histidine site upstream, two HRV 3C cleavage sites sandwiching the mScarlet-I-G3BP1(ΔE1/ΔE2) gene, and an MBP fusion gene downstream. The construct was called pOCC435-mScarlet-I-G3BP1($\Delta$E1/$\Delta$E2). The plasmid constructs for both insect cell extract ( TNT) and insect cell (*T.ni.*) expression are provided in supplementary figure 1.

| Buffer / solution | composition (pH) | Application |
|---|---|---|
| Lysis buffer (insect cells) | 50 mM Tris–HCl, 1 M NaCl, 5% (v/v) glycerol, pH 7.4 | Resuspension and lysis of *T. ni* insect cell pellets; pre-microfluidization. |
| MBP affinity buffer (Buffer A) | 20 mM Tris–HCl, 200 mM NaCl, pH 7.4 | Equilibration, loading, and low-salt washing of MBPTrap column. |
| High-salt wash buffer | 6$\times$ PBS, 1 mM DTT (pH $\sim$7.4) | High-salt wash (10 CV) of MBPTrap column to remove non-specific binders. |
| MBP elution buffer (Buffer B) | Buffer A + 10 mM maltose (20 mM Tris–HCl, 200 mM NaCl, 10 mM maltose, pH 7.4) | Isocratic elution of GdmS–MBP from MBPTrap column. |
| Cleavage / solubility buffer (post-MBP elution) | Buffer B adjusted to 400 mM NaCl and 500 mM arginine–HCl (pH 7.4) | H3C protease cleavage of MBP tag; prevention of aggregation of cleaved GdmS. |
| SEC / GdmS storage buffer | 50 mM phosphate buffer ($Na_2HPO_4$/$NaH_2PO_4$), 300 mM NaCl, 5% (v/v) glycerol, pH 7.0 | Superdex 200 16/60 SEC running buffer and final storage buffer for GdmS. |
| TNT extract reaction mixture | 80% (v/v) TNT T7 Insect Cell Extract (Promega L1101), 20% RNase-free water | Cell-free expression reactions in TNT extract. |
| Labeling buffer ($NaHCO_3$) | 100 mM sodium bicarbonate ($NaHCO_3$), pH 8.5 (freshly prepared) | NHS-ester labeling of T7 RNA polymerase and RNase A with ATTO-488 / Alexa Fluor 680. |
| Post-labeling storage buffer (labeled proteins) | Same as SEC / GdmS storage buffer: 50 mM phosphate buffer, 300 mM NaCl, 5% (v/v) glycerol, pH 7.0 | Final buffer after desalting labeled T7 RNA polymerase and RNase A; used in GdmS assays. |

Supplementary table 2: Buffers used for GdmS expression, purification, labeling, and cell-free experiments.

### 4 Transcription and purification of mRNA

The constructed plasmid containing the T7 promoter was linearized by NruI restriction enzyme at 37 ℃ and purified by QIAGEN PCR purification kit as described by the manufacturer's instructions. The mRNA was transcribed using the HiScribe T7 High yield RNA synthesis kit using the linearized plasmid at 37 ℃ for at least 4 hours. DNase I in DNase buffer, from the HiScribe

| Name | Sequence |
|---|---|
| SSO-030 | TAATACGACTCACTATAGGACGCCACTCTGAACGAC |
| SSO-031 | CCTATAGTGAGTCGTATTAATTTCGC |

Supplementary table 3: Primer sequences for generating the 1.8 knt mRNA.

kit, was added to the reaction and incubated at 37 ℃ for 30 minutes to degrade the template DNA. Two transcription reactions of 20 µL aliquots were pooled and the mRNA was purified with QIAGEN RNeasy kit as described by the manufacturers instructions and eluted with 40 µL of nuclease free water to achieve concentrations higher than 2000 ng/µL for the full length 2.8 knt mRNA.

For synthesis of the 1.8 knt mRNA, the plasmid pOCC499-mScarlet-I-G3BP1($\Delta$E1/E2) was used as the PCR template with primers SSO-030 and SSO-031 (see Table 3). After linearization using the NruI enzyme as described above, the primers were used to generate truncated copies of the plasmid using PCR. Following this, the RNA synthesis process was repeated as described above to generate the shorter 1.8 knt mRNA.

### 5 Expression and purification of GdmS

The GdmS protein was expressed in insect cells using a baculoviral expression system. GdmS has a deleted acidic region allowing for a lower phase separation threshold compared to the wildtype [2]. For each 500 mL culture of Trichoplusia ni (*T. ni.*) insect cells, 5 ml of baculoviral stock containing the plasmid encoding the GdmS fusion protein, pOCC435-mScarlet-I-G3BP1($\Delta$E1/$\Delta$E2), was added. All procedures were performed under sterile conditions to prevent bacterial contamination, which could compromise the entire culture [3]. Cultures were incubated at 30 ℃ for 72 hours. Successful expression was indicated by a pink hue in the medium, and microscopy revealed intact, enlarged cells, fluorescent under the RFP channel (Supplementary Figure 2). Lysis at this stage was avoided to prevent premature protein degradation. Cultures were harvested by centrifugation at 2000 × g for 5–7 minutes, and the resulting cell pellets were collected.

Pellets were resuspended in lysis buffer containing 50 mM Tris-HCl, 1 M NaCl, and 5% glycerol (pH 7.4). Notably, excessive glycerol concentrations may impair MBP-affinity interactions. The resuspended lysate was diluted at a 2:1 volume ratio with additional lysis buffer to facilitate passage through the LM20 microfluidizer (Microfluidics, USA), which was operated at 1000-1200 bars for three passes on ice to ensure thorough cell disruption. Following lysis, benzonase nuclease was added to degrade residual DNA to a final concentration of 100 U/ml, which could otherwise interfere with downstream phase separation assays.

Clarification of the cell lysis suspension was achieved by centrifugation at 50,000 × g for 40 minutes at 4 ℃. The supernatant was then filtered through a 0.22 $\mu$m sterile membrane filter to remove any remaining particulates. The filtered lysate was loaded slowly (1 ml/min) onto an MBP-Trap column pre-equilibrated with Buffer A (20 mM Tris-HCl, 200 mM NaCl, pH 7.4). The column was washed with 10 column volumes (CV) of Buffer A, followed by 10 CV of high-salt wash buffer (6x PBS, 1 mM DTT), and finally another 10 CV of Buffer A. Elution was performed isocratically with Buffer B (Buffer A supplemented with 10 mM maltose (maltose). To promote solubility and prepare for MBP tag removal, the eluate was adjusted to final concentrations of 400 mM NaCl and 500 mM Arginine-HCl. H3C protease was then added (1:50 (w/w)) to the GdmS-MBP eluate from MBPTrap column, and the mixture incubated for 1 hour at room temperature or overnight at 4 ℃. These additives were essential to prevent aggregation of the cleaved protein. Post-cleavage, the sample was concentrated to a final volume of approximately 5 ml using a 10

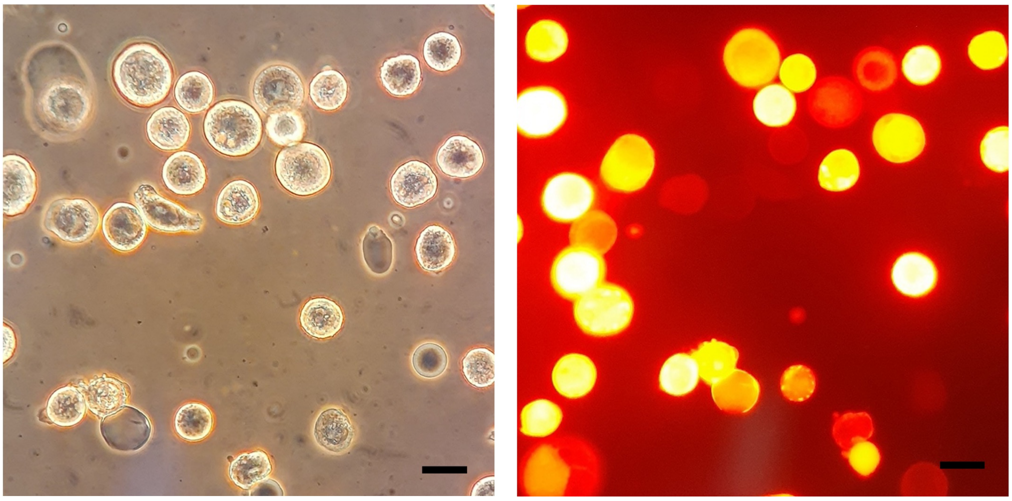

Supplementary figure 2: Left, brightfield micrograph of healthy *Tn.i.* cells that have been infected with baculoviral titre. Right, widefield fluoresence micrograph in the RFP channel, of baculovirus infected *T.ni.* cells after 72 hrs of incubation at 30 C showing expression of GdmS as indicated by fluoresence in the RFP channel. Scale bars denote 10 $\mu$m.

kDa molecular weight cutoff centrifugal concentrator. The device was operated gently at 2000 × g, with supernatant removed every 15 minutes via pipetting to minimize mechanical stress.

The sample was subsequently purified by size-exclusion chromatography using an FPLC system (Äkta Pure) equipped with a Superdex 200 16/60 column, pre-equilibrated in the final buffer that was 50 mM phosphate buffer, 300 mM NaCl, 5% glycerol, pH 7.0). Fractions were collected and monitored at 280 and 569 nm, and those corresponding to the major peak were pooled.

The protein was then concentrated to a final volume of 200–500 µL. Typical preparations yielded a final protein concentration of approximately 150-200 µM, allowing for efficient doping into cell free reactions without significant interference from the storage buffer.

The final product was aliquoted into 20 µL portions, flash-frozen in liquid nitrogen, and stored at –80 °C for long-term use. SDS-PAGE was performed at every step of the purification as shown in Supplementary Figure 3. A schematic of the entire purification protocol can be found in Supplementary Figure 4.

### 6 Fluorescent labeling of T7 RNA polymerase and RNase A

T7 RNA polymerase and RNase A were labeled with ATTO-488-NHS ester and Alexa Fluor 680-NHS ester, respectively. Both dyes were purchased as pre-activated NHS esters. For labeling, 10 $\mu$M of each protein was incubated with a 10-fold molar excess of the corresponding dye in 100 mM sodium bicarbonate buffer ($NaHCO_3$) that was freshly prepared and adjusted to pH 8.5. Reactions were performed in the dark at room temperature for 1 hour to allow covalent attachment of the dye to the enzymes.

Unreacted dye was removed by size exclusion using a 7 kDa molecular weight cut-off Zeba Spin Desalting Column with a 500 $\mu$L sample bed. To ensure compatibility with subsequent assays, an additional desalting step was performed to exchange the buffer into the GdmS-compatible storage buffer (50 mM phosphate buffer, 300 mM NaCl, 5% glycerol, pH 7.0). The labelled proteins were characterized by measuring the degree of labeling (DOL). This was quantified by UV–Vis spectrophotometry using the standard equation DOL=$A_{\lambda max}/\varepsilon_{dye}(A_{280}\text{-}CF\ A_{\lambda max})/\varepsilon_{protein}$ where $A_{\lambda max}$ and $A_{280}$are the absorbances at the dye's $\lambda_{max}$and at 280 nm, $\varepsilon_{dye}$ and $\varepsilon_{protein}$ are

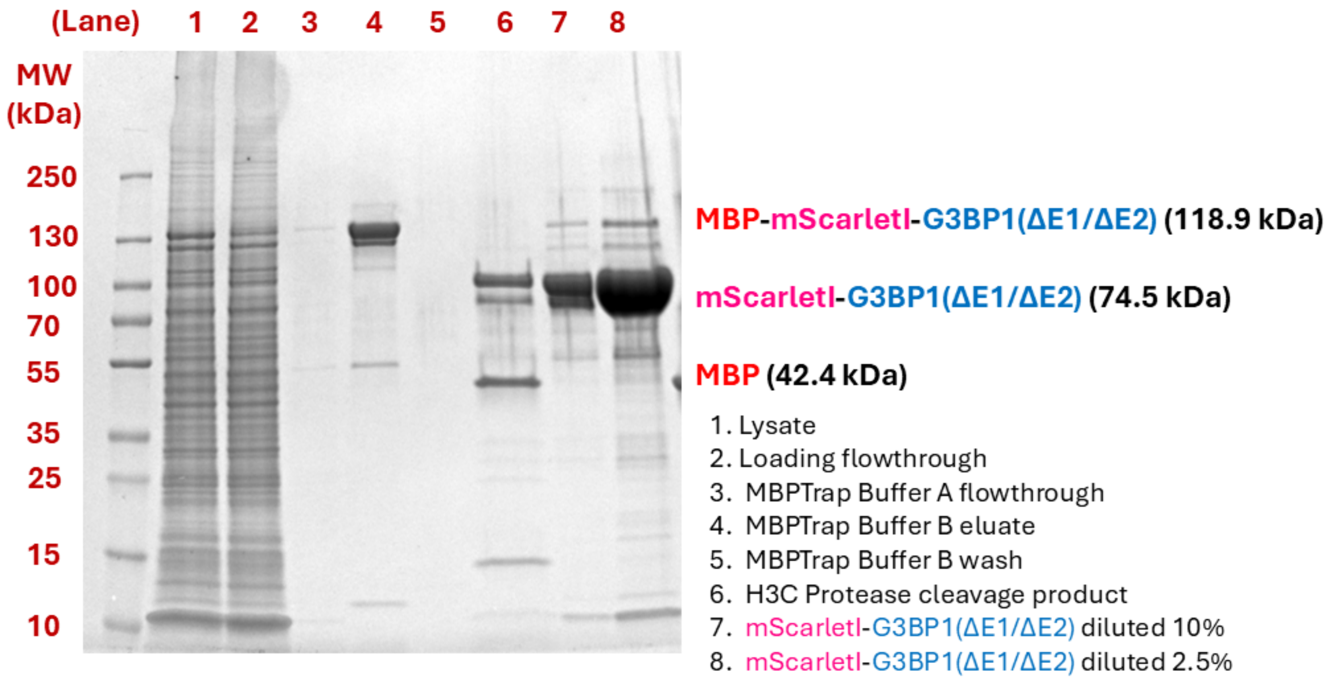

Supplementary figure 3: SDS-PAGE of GdmS (mScarletI-G3BP1(ΔE1/ΔE2) expressed in *T.ni.* cells samples across different steps of the purification process.

the molar extinction coefficients of the free dye and unlabeled protein, and CF corrects for dye absorbance at 280 nm. Dye-specific $\varepsilon$ and CF values from the manufacturers were used. The labeling efficiencies were 63% for T7 RNA polymerase-ATTO-488 and 67% for RNase A-Alexa Fluor 680. The protocol was adapted from Ref. [4].

### 7 Confocal microscopy

All confocal imaging was undertaken using a Zeiss LSM 880 confocal microscope equipped with an AiryScan detector and a C-Apochromat 40×/1.2 W objective, corrected for 0.17 mm coverslip thickness unless otherwise stated. Fluorescence imaging was conducted using a 561 nm laser for excitation, and emitted light was detected in the 579-650 nm window for mScarletI and 488 nm laser for excitation, and emitted light was detected in the 500-550 nm window for the Broccoli aptamer. Z-stacks were acquired in 12-bit bidirectional laser scanning mode over a 106.27 × 106.27 $\mu$m² field of view (512 × 512 pixels), averaged 4 times, with a Z-step size of 0.5 $\mu$m across a vertical height of approximately 40 $\mu$m. All experiments were undertaken at 30°C by equilibrating the temperature in a sample chamber prior to loading of the sample with a approximately 2 minutes of temperature equilibration for the sample.

### 8 Construction of Phase Diagrams for mRNA-GdmS Condensates

Phase diagrams for mRNA and GdmS protein mixtures in the TNT extract were constructed using confocal fluorescence microscopy and volumetric analysis, as detailed below.

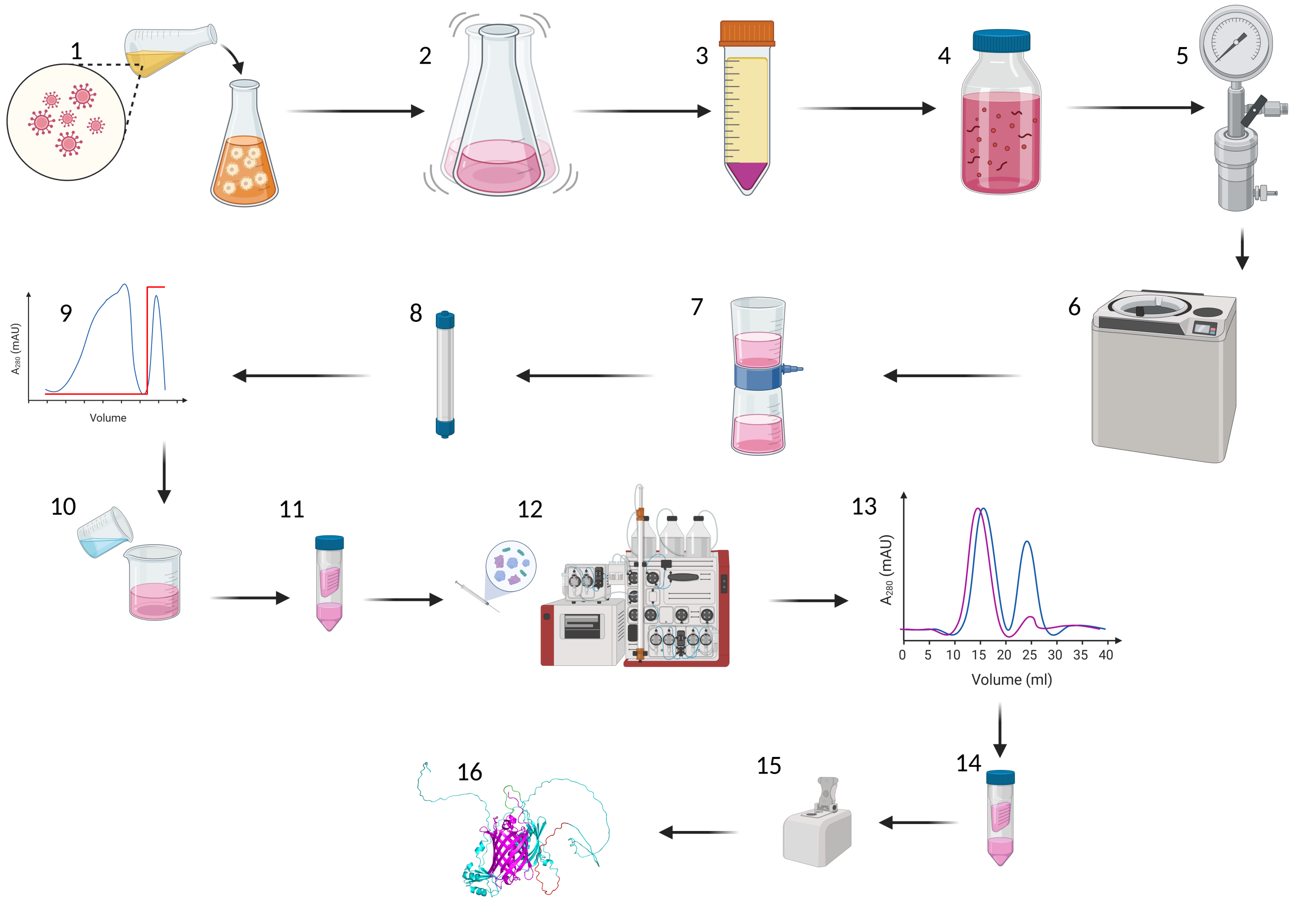

Supplementary figure 4: Purification steps for GdmS. The steps denoted are: 1. Baculoviral titre infection into T.ni. cell culture, 2. Shaking and incubation at 30 C for 72 hours, 3. Centrifugation to collect cell pellet, 4. Resuspension of cell pellet in lysis buffer, 5. Homogenization of the resuspension, 6. Centrifugation to clarify lysate, 7. Filtration of supernatant, 8. Flow through MBPTrap affinity column, 9. Isocratic elution using maltose buffer, 10. Addition of cleavage buffer with H3C protease, 11. Centrifugal concentration, 12. Loading onto size exclusion column, 13. Fluorescent peak fractions selected, 14. Centrifugal concentration, 15. Spectrophotometric concentration measurement, 16. Pure protein aliquoted and stored.

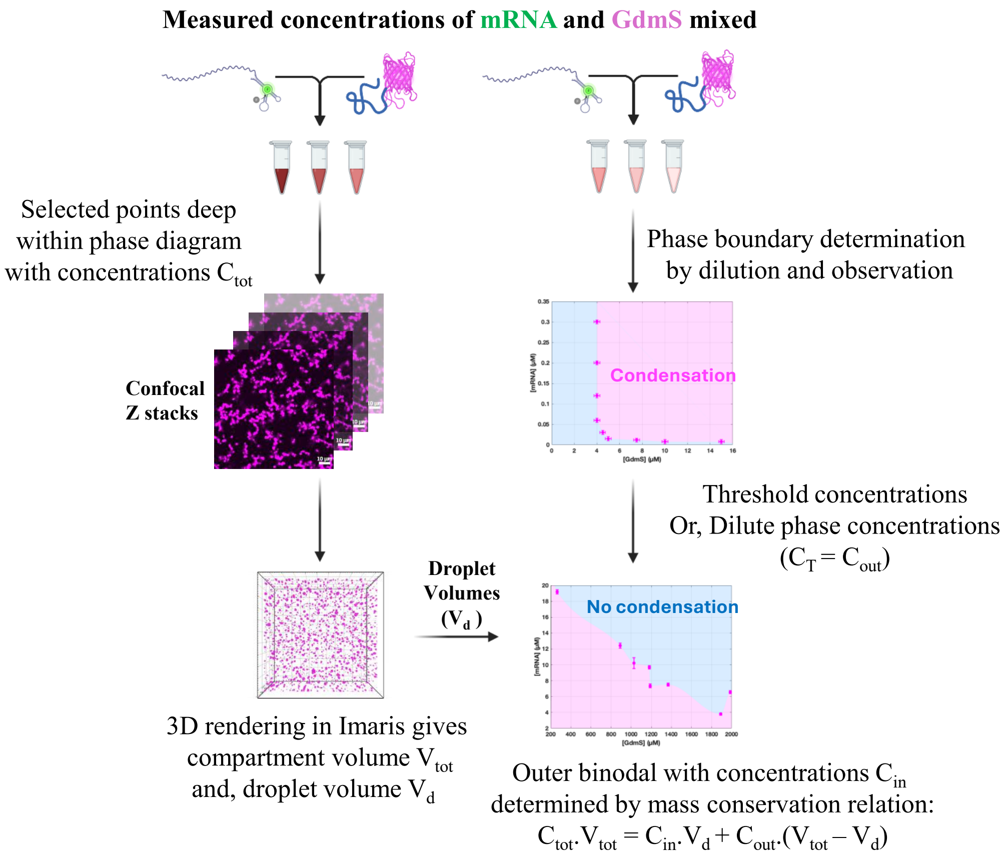

Supplementary figure 5: Schematic describing the protocol for determining the inner and outer binodals of the phase diagram experimentally.

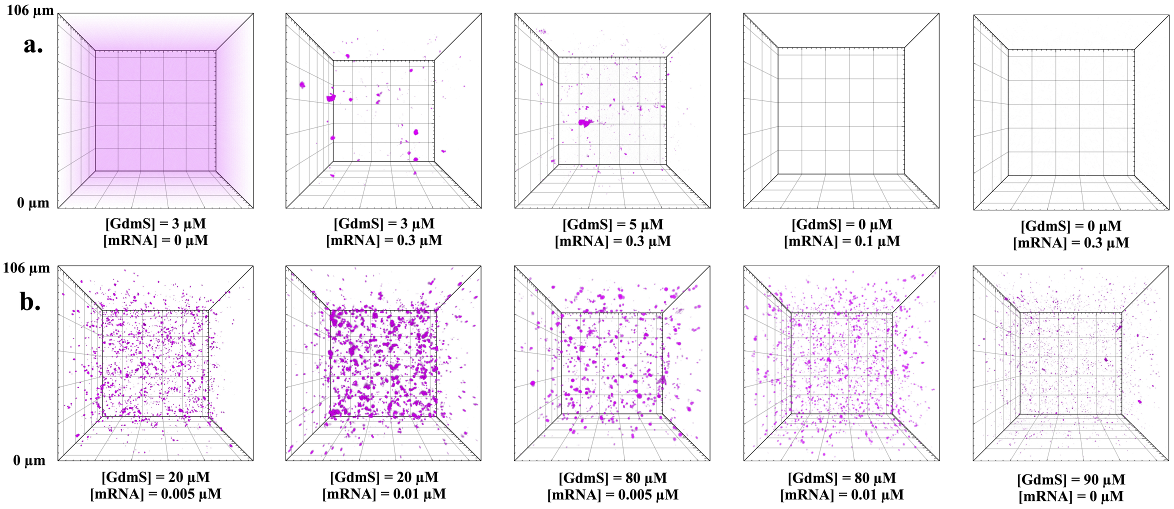

Supplementary figure 6: 3D confocal micrographs of GdmS-mRNA condensates used to determine the inner binodal with a) showing the binodal parallel to the mRNA axis and b) showing the inner binodal parallel to the GdmS axis. GdmS was found to self-condense at approximately 90 $\mu$M. The dimensions of each micrograph is $106\,\mu$m $\times$ $106\,\mu$m.

| [GdmS] / $\mu$M | [mRNA] / $\mu$M |
|---|---|
| $15.0 \pm 0.25$ | $0.0075 \pm 0.0050$ |
| $10.0 \pm 0.25$ | $0.0075 \pm 0.0050$ |
| $7.50 \pm 0.25$ | $0.0113 \pm 0.0050$ |
| $5.00 \pm 0.25$ | $0.0150 \pm 0.0050$ |
| $4.50 \pm 0.25$ | $0.0300 \pm 0.0050$ |
| $4.00 \pm 0.25$ | $0.0600 \pm 0.0050$ |
| $4.00 \pm 0.25$ | $0.1200 \pm 0.0050$ |
| $4.00 \pm 0.25$ | $0.2000 \pm 0.0050$ |

Supplementary table 4: Concentrations obtained for the GdmS poor-phase binodal. Errors correspond to the least count of visual distinguishability between two samples at different concentrations under the microscope.

#### 8.1 Determination of the GdmS poor phase binodal

To determine the GdmS poor phase (inner) binodal , a range of mRNA and GdmS concentrations were prepared by mixing defined amounts of purified mRNA and GdmS into 80% TnT extract. Each mixture was immediately transferred to an imaging chamber and incubated for a brief period at 30 °C before imaging by confocal microscopy (see supplementary information 7). The presence or absence of condensates was visually assessed from the acquired Z-stacks. An approximate phase boundary was defined from the transition between concentrations of mixtures for which condensation occurred vs when it did not. The concentrations that formed the GdmS poor phase binodal are provided in Table 4. The errors for the GdmS poor binodal came from the least count of visual distinguishability for two samples of different concentrations under the microscope.

#### 8.2 Determination of the GdmS rich phase binodal and tie lines

To identify the outer binodal of the phase diagram, mixtures were prepared with higher total concentrations of mRNA and GdmS to favor phase separation. After incubation in TNT extract,

| [GdmS] / $\mu$M | [mRNA] / $\mu$M |
|---|---|
| $1891.78 \pm 3.79$ | $4.45 \pm 0.13$ |
| $1987.20 \pm 6.55$ | $1.23 \pm 0.18$ |
| $1187.37 \pm 6.55$ | $0.76 \pm 0.23$ |
| $1367.19 \pm 7.50$ | $1.23 \pm 0.19$ |
| $1178.91 \pm 9.69$ | $1.29 \pm 0.19$ |
| $1025.49 \pm 10.22$ | $2.30 \pm 0.69$ |
| $890.64 \pm 12.41$ | $0.98 \pm 0.30$ |
| $264.27 \pm 19.20$ | $0.87 \pm 0.26$ |

Supplementary table 5: Concentrations obtained for the GdmS concentrated-phase binodal. Errors correspond to variability across multiple images taken from different regions of the same sample.

full 3D confocal Z-stacks were acquired under conditions as stated above. The Z-stack images were imported into Imaris 10.1 (Bitplane, Switzerland ) for quantitative 3D segmentation. Using the "Surfaces" module, condensates were detected via intensity thresholding and size-based filtering. The total condensate volume ($V_c$) was calculated as the sum of all segmented surfaces across the Z-stack. The volume of the entire imaging compartment (V) was computed from the x, y, and z dimensions of the confocal volume of the imaging field.

To determine the GdmS poor phase concentration ($C^{out}$) for both mRNA and GdmS, the total input concentration ($C^{avg}$) for mRNA and GdmS for each sample was mapped to its closest point on the previously determined inner binodal boundary. Using this value and the measured condensate and compartment volumes, the concentration inside the droplets ($C^{in}$) for mRNA and GdmS was calculated by applying the principle of mass conservation:

$$C^{\text{avg}}V = C^{\text{in}}V_c + C^{\text{out}}(V - V_c)\,. \tag{1}$$

This applied to both mRNA and GdmS so the relation for the individual concentration of mRNA inside the droplets was therefore:

$$[\text{mRNA}]^{\text{in}} = (([\text{mRNA}]^{\text{avg}} - [\text{mRNA}]^{\text{out}})V + [\text{mRNA}]^{\text{out}}V_c)/V_c\,. \tag{2}$$

And likewise, for GdmS,

$$[\text{GdmS}]^{\text{in}} = (([\text{GdmS}]^{\text{avg}} - [\text{GdmS}]^{\text{out}})V + [\text{GdmS}]^{\text{out}}V_c)/V_c\,. \tag{3}$$

This relationship allowed for an approximate estimation of the GdmS rich phase concentration $C_{in}$ under conditions where macroscopic phase separation was evident. These concentrations were defined as the outer binodal of the phase diagram. These concentrations calculated from the relations above are given in Table 5. The errors for the GdmS rich phase binodal came from multiple images taken from different areas of the same sample. The experimental tie-lines were drawn using the points from the outer binodal to the points on the inner binodal that were used to calculate the points on the outer binodal.

### 9 Cell free expression and quantification of mRNA and GdmS.

#### 9.1 Calibration of Fluorescence Signals for well plate reader experiments.

Fluorescence intensities for both mScarlet-I protein and Broccoli-labelled mRNA were converted into concentration values using independent calibration curves. For protein calibration, purified

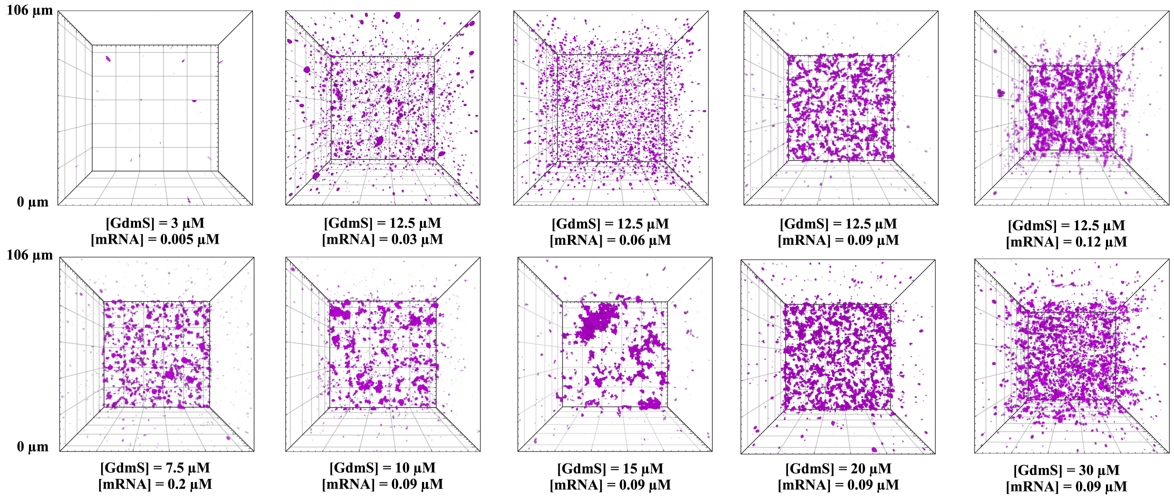

Supplementary figure 7: 3D confocal micrographs of gel-like GdmS-mRNA condensates in 80% TnT extract for various combinations involved in the process of generating experimental tie lines. These mixtures show regions deep inside the GdmS rich region of the phase diagram. The dimensions of each micrograph is $106\,\mu$m $\times$ $106\,\mu$m.

mScarlet-I protein was serially diluted across a range that bracketed the highest experimental signal. The concentrations were 0.039875, 0.07975, 0.1595, 0.319, 0.638, and 1.276 $\mu$M. These standards were prepared in the same buffer and TNT conditions as the test reactions. The fluorescence intensity values were fit to a hyperbolic curve with the equation:

$$I_{\text{GdmS}}^{\lambda=594\text{nm}} = (a[\text{GdmS}])/(b - [\text{GdmS}])\,. \tag{4}$$

mRNA calibration was performed by maintaining DFHB1 at a constant concentration of 20 $\mu$M (This concentration was chosen to make sure an excess amount of free dye is present in the solution to drive the binding to mRNA) and varying the concentration of the transcribed and purified mRNA from the pOCC499-mScarlet-I-G3BP1($\Delta$E1/E2) plasmid, containing the Broccoli aptamer. The concentrations of mRNA used were 0, 25, 50, 100, 200, 300, and 500 nM. The resulting fluorescence values were fit to a linear calibration curve with equation:

$$I_{[\text{mRNA}]}^{\lambda=450\text{nm}} = a + b[\text{mRNA}]\,. \tag{5}$$

MATLAB was used to fit both calibration datasets to their respective models.

#### 9.2 Cell free expression of mRNA and mScarlet-I-G3BP1($\Delta$E1/$\Delta$E2) (GdmS).

To monitor transcription and translation in a cell-free environment, reactions were carried out using commercially available insect cell TNT extract following the manufacturer's protocol. Each 10 $\mu$L reaction mixture consisted of 80% (v/v) TNT extract diluted in nuclease-free water, supplemented with 5.6 nM plasmid DNA encoding GdmS, pOCC499-mScarlet-I-G3BP1($\Delta$E1/$\Delta$E2). For experiments requiring additional enzymatic doping, 1 $\mu$M T7 RNA polymerase and 0.1 $\mu$M RNase A were included. The mRNA produced contained a Broccoli aptamer, enabling fluorescence-based quantification upon binding with the fluorophore DFHBI, which was added to each reaction at a final concentration of 20 $\mu$M.

Reactions were assembled in triplicate and pipetted into a black 384-well flat-bottom microplate (Greiner, Cat. No.: 784076), sealed with an adhesive film, and transferred immediately to a TECAN Spark 20M microplate reader. The reader was pre-equilibrated to 30 ℃ until a stable

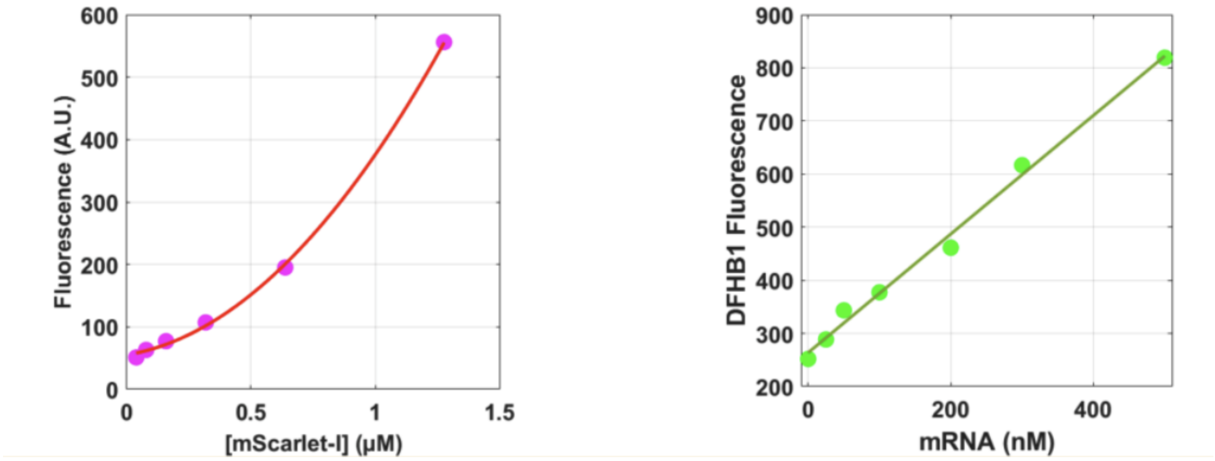

Supplementary figure 8: Calibration curves for well-plate reader experiments for (left) mScarlet-I ($\lambda_{\text{exc}} = 569$ nm, $\lambda_{\text{em}} = 594$ nm) and (right) mRNA ($\lambda_{\text{exc}} = 450$ nm, $\lambda_{\text{em}} = 501$ nm).

temperature reading was obtained. Fluorescence was monitored simultaneously for mRNA and protein production over a 4-hour period, with data collected every 5 minutes. Prior to each read, the plate was shaken at 330 rpm for 3 seconds. The mScarlet-I signal, indicative of protein expression, was recorded using an excitation wavelength of 569 nm and an emission wavelength of 594 nm, with a 10 nm bandwidth, gain setting of 70, and three flashes per measurement. mRNA was measured via the fluorescence from DFHBI-Broccoli complex, using an excitation wavelength of 450 nm and emission at 501 nm under identical bandwidth, gain, and flash settings. All readings were conducted with the Z-position set to 17,500 $\mu$m for optimal focal alignment during scanning.

#### 9.3 Supplementary note: Modelling peak mRNA and GdmS concentrations with respect to titrants

The observed titration behavior of peak mRNA concentration ($\text{mRNA}_{\text{max}}$) and GdmS concentration at equilibrium ($\text{GdmS}_{\text{max}}$) as a function of T7 RNA polymerase and DNA template concentrations was fit to an approximated analytical derivation from the one phase model to explain the titration behaviors as follows:

Although mass action kinetics were primarily used to build the one phase model, here we assume the transcription rate is governed by Michaelis–Menten kinetics due to limited transcription resources (TsR, T7):

$$\frac{d[\text{mRNA}]}{dt} = \frac{k_r[\text{DNA}][\text{TsR}][\text{T7}]}{(K_m + [\text{T7}])} - k_d[\text{RNase}][\text{mRNA}]\,. \tag{6}$$

At steady state $\frac{d[\text{mRNA}]}{dt} = 0$ , therefore,

$$\frac{k_r[\text{DNA}][\text{TsR}][\text{T7}]}{(K_m + [\text{T7}])} = k_d[\text{RNase}][\text{mRNA}]_{\text{max}}\,, \tag{7}$$

which has a saturable form that fit well with the $[\text{mRNA}]_{\text{max}}$ vs $[\text{T7}]_{\text{data}}$:

$$[\text{mRNA}]_{\text{max}} = \frac{C_1[\text{T7}]}{C_2(K_m + [\text{T7}])}\,. \tag{8}$$

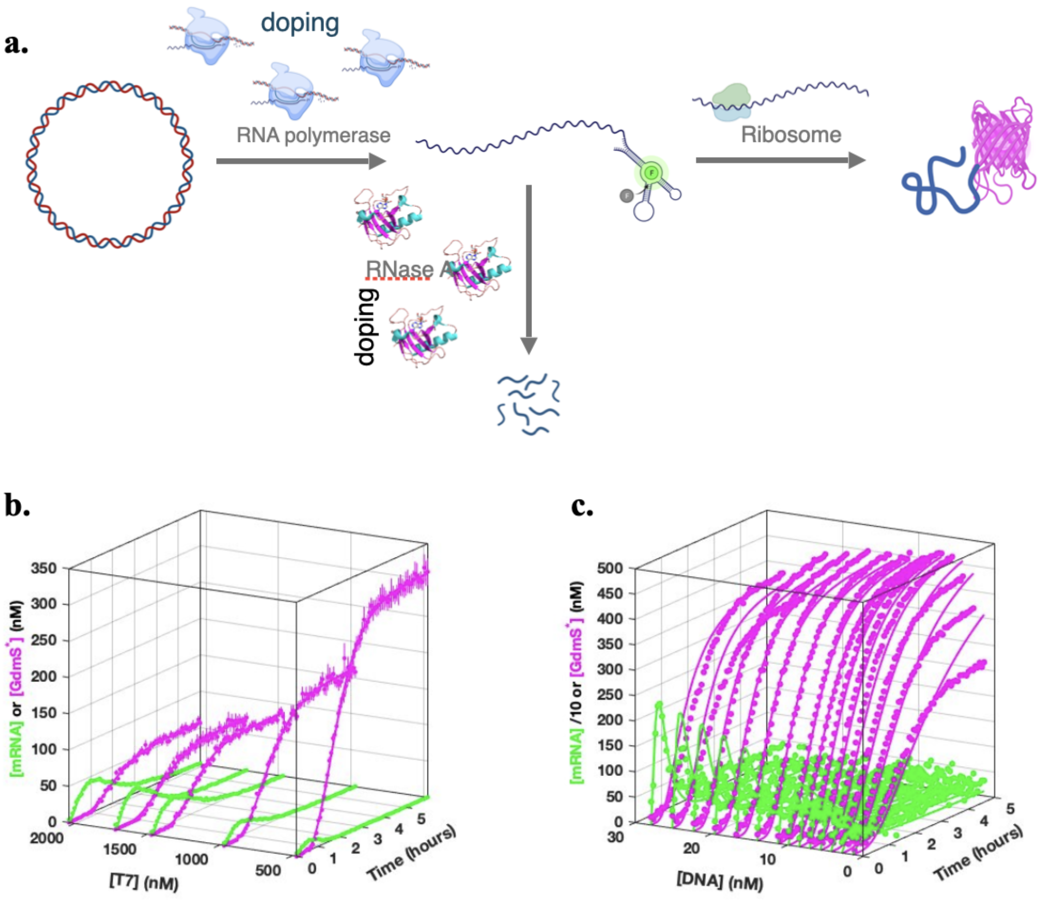

Supplementary figure 9: Cell-free extract based transcription translation under homogeneous conditions (no phase separation). a) A schematic shows the production and degradation scheme mediated by T7 RNA polymerase and RNase the concentrations for which can be controlled by doping. Kinetic curves for mRNA (green) and GdmS (pink) with b) changing T7 polymerase condentrations and c) DNA concentrations.

Ignoring maturation kinetics, the rate of GdmS production is approximately given by:

$$\frac{d[\text{GdmS}]}{dt} = k_p[\text{mRNA}][\text{TIR}]\frac{[\text{TsR}]}{2}\,. \tag{9}$$

Doping in excessive T7 RNA polymerase is predicted to deplete TsR which can hinder GdmS production at the expense of mRNA production. Thus, we can model maximum mRNA and GdmS concentrations as inversely proportional to each other:

$$[\text{mRNA}]_{\text{max}} = \frac{C}{[\text{GdmS}]_{\text{max}}}\,. \tag{10}$$

Substituting the expression for $[\text{mRNA}]_{\text{max}}$ into this relation:

$$[\text{GdmS}]_{\text{max}} = \frac{C_3(K_m + [\text{T7}])}{C_1[\text{T7}]}\,, \tag{11}$$

which has a decaying asymptotic behavior that fits well to the $[\text{GdmS}]_{\text{max}}$ vs [T7] data. This behavior of translation suppression due to enhancement of transcription was predicted by the model which provides an empirical validation of the model.

Similar to the T7 RNA polymerase dependence of peak mRNA concentrations, an equivalent expression for a DNA titrant can be represented as:

$$[\text{mRNA}]_{\text{max}} = \frac{C_1[\text{DNA}]}{C_2(K_m + [\text{DNA}])}\,. \tag{12}$$

For the peak GdmS concentration we get the same expression as before only that this time we add an inhibitory term with respect to excessive addition of DNA which can deplete TsR:

$$[\text{GdmS}]_{\text{max}} = \frac{C_3(K_m + [\text{DNA}])}{C_1[\text{DNA}]}\frac{1}{1 + [\text{DNA}]/K_i}\,, \tag{13}$$

which gives a non monotonic curve with peak GdmS concentration rising at low DNA concentrations because the first term dominates, and it decreasing at higher DNA concentrations resulting in an optimal DNA concentration in agreement with the $[\text{GdmS}]_{\text{max}}$ vs [DNA] data. Both TsR and TIR, while time dependent, were absorbed into the constants as a time-averaged effective value around the narrow time window at which mRNA peaks.

Fitting the data to this model in which transcription resources were coupled to both transcription and translation rates (Figure 2D1-D2) shows that increasing DNA concentration led to a rise in mRNA levels reaching a plateau of 15 nM, and GdmS reached a maximum concentration of 450 nM at 9.4 nM DNA Figure 2D1), not far from the commercial recommendation for DNA concentration of 5.6 nM for the TNT extract which we use for our experiments. (See supplementary methods table 2).

### 10 Observation of dynamic behaviour of GdmS and mRNA

To measure the dynamics of condensate growth and dissolution over time, time-lapse 3D confocal Z-stacks were acquired using settings described in the earlier section. Experiments were conducted in Ibidi µ-Slide VI 0.1 chambers pre-equilibrated at 30℃ using a microscope box chamber incubator. 10 µL samples were prepared by diluting TNT extract to 80% (v/v) in nuclease-free water, supplemented with dopants as in 4.5–20$\mu$M GdmS, 0 or 1 $\mu$M T7 RNA polymerase, and 0 or 0.1 $\mu$M RNase A. The plasmid DNA was added as a final concentration

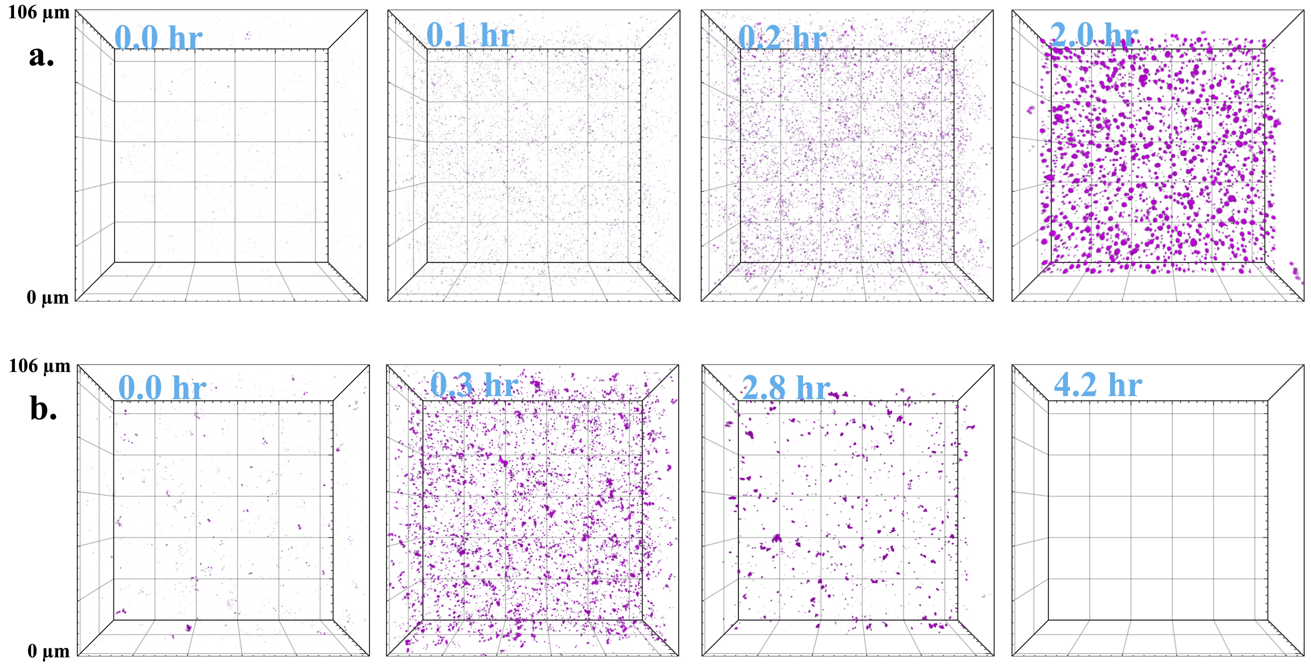

Supplementary figure 10: Timelapse 3D confocal micrographs of mRNA production and degradation driven formation and dissolution of GdmS-mRNA condensates with a) 5 $\mu$M GdmS doped and b) 5 $\mu$M GdmS, 1 $\mu$M T7 polymerase and 0.1 $\mu$M RNase A doped.

of 5.6 nM immediately prior to pipetting into the chambered slide. Both inlets were sealed with 40 µL of anti-evaporation oil to minimize evaporation. Operational delays between mixing and imaging were minimized. Z-stacks were acquired continuously for at least 2-4 hours. The timelapse function was added upon the Z-stack acquisition settings duplicated from that used for the construction of the phase diagram, except that the averaging number was reduced to 2 for faster acquisition. The method was adapted from the reference [4].

All Z-stack timelapses were processed in Imaris 10.1 (Bitplane, Switzerland) and total condensate volumes for each frame were obtained as described previously in the phase diagram construction section. The volumes were divided by the compartment volume (typically, for 80 slices with a slice width of 0.5 $\mu$m, and a cropped area of 106 x 106 $\mu m^2$, the compartment volume was 449.4 picoliters). This resulting volume fraction data over time were plotted with the simulation data.

### 11 Theoretical model and definition of parameters

Here we describe the theoretical framework used to describe the system, including the full reaction scheme, definition of chemical reaction rates, determination of phase equilibrium and finally the extraction of rate constants associated with transcription and degradation inside and outside the droplets.

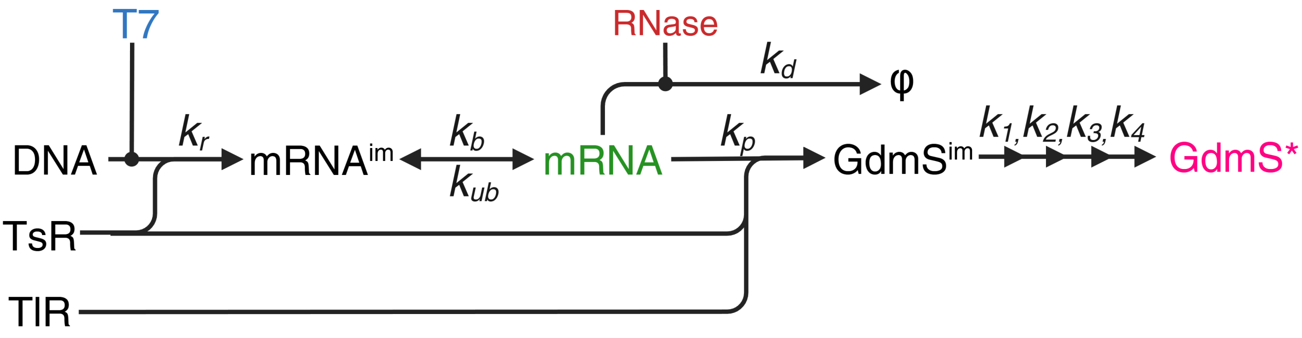

Supplementary figure 11: Schematic of the rate equations for the one phase kinetic model which includes contiguous maturation steps $k_1$, $k_2$, $k_3$, $k_4$ from the immature GdmS to fluorescent GdmS.

#### 11.1 Transcription-translation scheme

A resource-limited, mass action model to describe transcription and translation is stated as follows:

$$\text{TsR} \xrightarrow{\text{DNA,T7}} \text{mRNA}_{\text{im}} \rightleftharpoons \text{mRNA} \tag{14a}$$

$$\text{mRNA} \xrightarrow{\text{RNase}} \phi \tag{14b}$$

$$\text{TIR} + \text{TsR} \xrightarrow{\text{mRNA}} \text{GdmS}_0 \tag{14c}$$

$$\text{GdmS}_0 \rightarrow \text{GdmS}_1 \rightarrow \text{GdmS}_2 \rightarrow \text{GdmS}_3 \rightarrow \text{GdmS}^* \,. \tag{14d}$$

Here, [$\text{mRNA}_{\text{im}}$] refers to the immature mRNA which has not yet bound to the DFHBI dye, $[\text{GdmS}]_0$ is the concentration of non-fluorescent protein, and $[\text{GdmS}]^*$ is the final mature form of the protein.
The schematic is given in Supplementary figure 11 and the corresponding differential equations are given as follows:
The production of immature mRNA (mRNA with Broccoli aptamer unbound to DFHBI dye) depends on the concentration of DNA, the transcription resources TsR and the T7 RNA polymerase T7. The removal of the immature mRNA occurs by maturation to mRNA by DFHBI binding to the broccoli aptamer once it has undergone folding.

$$\frac{d[\text{mRNA}_{\text{im}}]}{dt} = k_r[\text{DNA}][\text{TsR}][\text{T7}] - k_b[\text{mRNA}_{\text{im}}] + k_{ub}[\text{mRNA}] \,. \tag{15}$$

The production of dye bound mRNA depends on the concentration of immature mRNA. The depletion of mRNA depends on degradation of mRNA by RNase as well as the dye unbinding from it. We ignore the possibility of the immature mRNA degrading as well since the binding kinetics is relatively fast.

$$\frac{d[\text{mRNA}]}{dt} = -k_d[\text{mRNA}][\text{RNase}] + k_b[\text{mRNA}_{\text{im}}] - k_{ub}[\text{mRNA}] \,. \tag{16}$$

The production of immature non-fluorescent GdmS depends on mRNA as well as the translation resources and half of the transcription resources. This is rationalized by the understanding that TsR is primarily the four nucleotides and translation processes require half of that, viz. GTP and ATP, wherein ATP is used up by aminoacyl-tRNA synthetases to charge tRNAs with amino acids, and GTP is used up by translation factors for initiation, elongation and termination. A cascade of processes is incorporated further to model the delay in the production of the final fluorescent protein, GdmS*.

$$\frac{d[\text{GdmS}_0]}{dt} = \frac{k_p[\text{mRNA}][\text{TsR}][\text{TIR}]}{2} - k_1[\text{GdmS}_0]\,, \tag{17a}$$

$$\frac{d[\text{GdmS}_1]}{dt} = k_1[\text{GdmS}_0] - k_2[\text{GdmS}_1]\,, \tag{17b}$$

$$\frac{d[\text{GdmS}_2]}{dt} = k_2[\text{GdmS}_1] - k_3[\text{GdmS}_2]\,, \tag{17c}$$

$$\frac{d[\text{GdmS}_3]}{dt} = k_3[\text{GdmS}_2] - k_4[\text{GdmS}_3]\,, \tag{17d}$$

$$\frac{d[\text{GdmS}^*]}{dt} = k_4[\text{GdmS}_3]\,. \tag{17e}$$

The transcription and translation resource decay are modeled as:

$$\frac{d[\text{TsR}]}{dt} = -k_r[\text{DNA}][\text{TsR}][T7] - \frac{k_p[\text{mRNA}][\text{TIR}][\text{TsR}]}{2}\,, \tag{18a}$$

$$\frac{d[\text{TIR}]}{dt} = -\frac{k_p[\text{mRNA}][\text{TIR}][\text{TsR}]}{2}\,. \tag{18b}$$

The overall translation rate constant was defined as the harmonic mean of $k_p$ and $k_1$ through $k_4$:

$$k_p^{\text{overall}} = \frac{1}{\frac{1}{k_p} + \frac{1}{k_1} + \frac{1}{k_2} + \frac{1}{k_3} + \frac{1}{k_4}}\,. \tag{19}$$

The maturation half-life was calculated as:

$$\tau_{\text{mat}}^{1/2} = \ln 2 \left( \frac{1}{k_1} + \frac{1}{k_2} + \frac{1}{k_3} + \frac{1}{k_4} \right)\,. \tag{20}$$

For ease of annotation in the future we refer GdmS* simply as GdmS, denoting it is the mature fluorescent form.

#### 11.2 Transcription-translation kinetics at phase equilibrium

When studying the interplay between phase separation and transcription-degradation, we made some simplifications in the above reaction scheme: the step describing the mRNA binding to DFHBI and the GdmS maturation steps were ignored. The set of reactions considered in this case was:

$$\text{TsR} \xrightarrow{\text{DNA,T7}} \text{mRNA} \tag{21}$$

$$\text{mRNA} \xrightarrow{\text{RNase}} \phi \tag{22}$$

$$\text{TIR} + \text{TsR} \xrightarrow{\text{mRNA}} \text{GdmS} \tag{23}$$

We consider that the same reactions take place in each phase, wherein concentrations and rate constants are designated with a superscript $\alpha$ standing for either the GdmS rich phase ($\alpha$ = in) or the GdmS poor phase ($\alpha$ = out). Here, mRNA and GdmS form the scaffold for the condensate phase, and the dilute components or clients can partition into each of the phases depending on an intrinsic partition coefficient.

All components except mRNA and GdmS were considered to be dilute so they were approximated to not have an effect on the phase separation. The mRNA-GdmS droplets are considered to be at phase equilibrium at all times, which is a valid assumption in the fast diffusion limit where the reaction diffusion length scales given by $\sqrt{D^\alpha/k^\alpha_{r,\text{obs}}}$ and $\sqrt{D^\alpha/k^\alpha_{r,\text{obs}}}$ are much larger compared to the condensate size [5]. Here D represents diffusion coefficient of the components, $k_{r,\text{obs}}$ and $k_{d,\text{obs}}$ are the observed transcription and degradation rate constants respectively. The value of D for GdmS in a condensate as calculated by FRAP was 0.6 $\mu$m$^2$/min (see section 18) and the observed rate constant for both transcription and degradation was of the order of 0.01 hr$^{-1}$ (Figure 3(C)). The reaction diffusion lengthscale calculated from these values came to be around 60 $\mu$m which was an order of magnitude larger than a typical size for the largest condensate at 5 $\mu$m. This allows us to ignore spatial gradients in concentration, and solve ODEs for the GdmS rich and GdmS poor phase concentrations.

#### 11.3 Reaction rates in each phase

As mentioned in section 11.1, the reaction rates of each component were assumed to follow mass action kinetics. Since reaction rates depend on the concentration of mRNA, reactions in general occur at different rates in each phase. In addition, the reaction rate constants can also depend on the concentrations of mRNA and GdmS, and therefore on the phase. We assume that the transcription and translation resources partition equally in each phase, while as described in the main text, we found the enzymes to preferentially partition inside the droplets. Altogether, we can write the reaction rates in each phase for all components as

$$r^\alpha_{\text{m-RNA}} = k^\alpha_r[\text{DNA}][\text{TsR}][\text{T7}]^\alpha - k^\alpha_d[\text{mRNA}]^\alpha[\text{RNase}]^\alpha\,, \tag{24a}$$

$$r^\alpha_{\text{GdmS}} = \frac{k^\alpha_p[\text{mRNA}]^\alpha[\text{TsR}][\text{TLR}]}{2}\,, \tag{24b}$$

$$r^\alpha_{\text{TsR}} = -k^\alpha_r[\text{DNA}][\text{TsR}][\text{T7}]^\alpha - \frac{k^\alpha_p[\text{mRNA}]^\alpha[\text{TsR}][\text{TLR}]}{2}\,, \tag{24c}$$

$$r^\alpha_{\text{TLR}} = -k^\alpha_p[\text{mRNA}]^\alpha[\text{TsR}][\text{TLR}]\,. \tag{24d}$$

In the two-phase scenario, we ignore the change in [GdmS] due to translation since the doped concentration is much higher.

#### 11.4 Kinetic equations for average mRNA concentration

We will now obtain an expression for the evolution of average concentration of mRNA for a two phase system due to transcription and degradation of mRNA in each phase at phase equilibrium.

As shown by Bauermann et al. [5], we can express the rate of concentration change of mRNA in each phase in terms of the chemical flux $r_{\text{mRNA}}$, the exchange flux $j_{\text{mRNA}}$ between phases and the rate of change of volume of the GdmS rich phase:

$$\frac{d[\text{mRNA}]^\alpha}{dt} = r^\alpha_{\text{mRNA}} - j^\alpha_{\text{mRNA}} - \frac{[\text{mRNA}]^\alpha}{V^\alpha}\frac{dV^\alpha}{dt}\,, \tag{25}$$

where $r^\alpha_{\text{mRNA}}$ is the net rate of production of mRNA by chemical processes, and $j^\alpha_{\text{m-RNA}}$ is the rate diffusive exchange of mRNA molecules between the phases. This allows us to write

$$\frac{d[\text{mRNA}]^{\text{avg}}}{dt} = \frac{V_c(r^{\text{in}}_{\text{mRNA}} - j^{\text{in}}_{\text{mRNA}}) + (V-V_c)(r^{\text{out}}_{\text{mRNA}} - j^{\text{out}}_{\text{mRNA}})}{V}\,. \tag{26}$$

Since diffusive exchange conserves the total number of mRNA molecules, $V_c\, j^{\text{in}} = -(V-V_c)j^{\text{out}}$, which leaves us with

$$\frac{d[\text{mRNA}]^{\text{avg}}}{dt} = \frac{V_c r^{\text{in}}_{\text{mRNA}} + (V-V_c)r^{\text{out}}_{\text{mRNA}}}{V}\,. \tag{27}$$

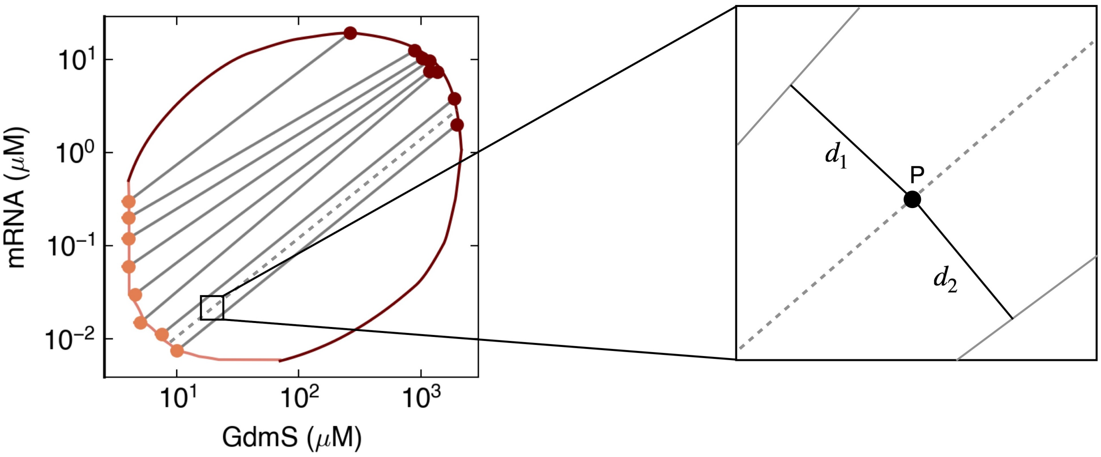

Supplementary figure 12: Determination of GdmS rich and GdmS poor phase concentrations. For a point P lying between two experimental obtained tie-lines with slopes $m_1$ and $m_2$, the slope of the tie-line passing through P is the linear interpolation between the values $m_1$ and $m_2$, weighted by the distance of P from the tie-lines. More specifically $m_P = m_1 \frac{d_2}{d_1+d_2} + m_2 \frac{d_1}{d_1+d_2}$.

Thus, given the reaction rates in each phase, the rate of change of [mRNA]$^{\text{avg}}$ can be evaluated. Given [mRNA]$^{\text{avg}}$(t), the volume of the condensates over time can be determined as described in the next section.

#### 11.5 Thermodynamic phase diagram for determining equilibrium concentrations and condensate volume

For a given average mRNA concentration, we determine the corresponding equilibrium concentrations for each phase and as a result, the condensate volume fraction. Experimentally measured equilibrium concentrations described in section 8 were used to define the GdmS poor and GdmS rich phase binodals. For the GdmS poor phase binodal, pairwise linear interpolation was used between experimental data points in the GdmS poor phase region. Similarly, to define the GdmS rich phase binodal pairwise linear interpolation was used for the corresponding GdmS rich phase data points. The dense binodal thus obtained was quadratically extrapolated to complete the binodal manifold in the form of a hull, see supplementary figure 12.

Any point ([GdmS]$^{\text{add}}$,[mRNA]$^{\text{avg}}$) in the binodal region is connected to the GdmS rich and GdmS poor phase binodals with a unique tie-line. To determine the slope of this tie-line, we linearly interpolated the tie-line slopes of experimentally determined tie-lines, and linearly extrapolated them beyond the experimentally measured region. See supplementary figure 12. The intersection of the tie line with the binodal gives [mRNA]$^{\text{in/out}}$.

The average concentration [mRNA]$^{\text{avg}}$ can be expressed as

$$[\text{mRNA}]^{\text{avg}} = \frac{V_c[\text{mRNA}]^{\text{in}} + (V - V_c)[\text{mRNA}]^{\text{out}}}{V} . \tag{28}$$

Eq. (28) can be rearranged to give an expression for the condensate volume fraction $V_f$:

$$V_f = \frac{V_c}{V} = \frac{[\text{mRNA}]^{\text{avg}} - [\text{mRNA}]^{\text{out}}}{[\text{mRNA}]^{\text{in}} - [\text{mRNA}]^{\text{out}}} . \tag{29}$$

Thus for any point ([GdmS]$^{\text{add}}$,[mRNA]$^{\text{avg}}$) in the binodal region, we can calculate the $V_f$ from the phase diagram as described above. By knowing the equilibrium concentrations of mRNA and GdmS, we can evaluate the reaction rates $r_i^\alpha$ in each phase $\alpha$ (section 11.2), determine the rate of change of $[\text{mRNA}]^{\text{avg}}$ (section 11.4), and thus, the evolution of $V_f$ for subsequent time steps (section 11.5). We used $V_f$ obtained from the model to extract reaction rate constants, by comparing the model prediction to experimental data.

#### 11.6 Phase dependence of rate constants

When writing mass action laws for non-dilute components, the corresponding rate constants are in general concentration dependent [5]. Therefore $k_r$ and $k_d$ in Eq.(24a) in each phase depend on the respective [mRNA] and [GdmS] concentrations in each phase. We estimated this dependence empirically, given the complexity of the system. We used experimental data for condensate volume fraction for 8 different experimental conditions; more specifically different values of $[\text{GdmS}]^{\text{add}}$, $[\text{T7}]^{\text{add}}$ and $[\text{RNase}]^{\text{add}}$, which are the doped concentrations of GdmS, T7 RNA polymerase and RNase A. We obtained $k_r^{\text{in/out}}$ and $k_d^{\text{in/out}}$ for each experimental condition independently by fitting our model to experimental volume traces. We strikingly found that the rate constants $k_r^{\text{out}}$ and $k_d^{\text{out}}$ obtained by from the fits decay exponentially with $[\text{GdmS}]^{\text{add}}$, see supplementary figure 13. This dependence on GdmS concentration can be possibly explained as follows.

Let us consider the degradation reaction of mRNA in each phase. The degradation rate in terms of its chemical potential $\mu_{\text{mRNA}}$ is given as

$$r^\alpha_{\text{mRNA}} = -k^\alpha \exp\left(\beta \mu^\alpha_{\text{mRNA}}\right) , \tag{30}$$

where $k$ is a kinetic constant, $\beta = 1/k_B T$ and $\alpha$ is the phase index, see the reference [5]. The chemical potential $\mu_{\text{mRNA}}$ can be further written as

$$\mu^\alpha_{\text{mRNA}} = \mu^0_{\text{mRNA}} + k_B T \log([\text{mRNA}]^\alpha \gamma^\alpha_{\text{mRNA}}) , \tag{31}$$

where $\mu^0_{\text{mRNA}}$ is the reference chemical potential of mRNA and $\gamma^\alpha_{\text{mRNA}}$ is the chemical activity coefficient, as defined using the Flory-Huggins free energy density in the reference [5]. Substituting $\mu^\alpha_{\text{mRNA}}$ in Eq. (30), we get

$$r^\alpha_{\text{mRNA}} = -k^\alpha \exp\left(\beta \mu^0_{\text{mRNA}}\right) [\text{mRNA}]^\alpha \gamma^\alpha_{\text{mRNA}} . \tag{32}$$

The chemical activity coefficient $\gamma^\alpha_{\text{mRNA}}$ is given by

$$\begin{aligned}\gamma^\alpha_{\text{mRNA}} = \frac{C}{[\text{mRNA}]^\alpha}\Bigg[\exp\Bigg(&\frac{\nu_{\text{GdmS}}\cdot\nu_{\text{mRNA}}}{\nu_{\text{solvent}}}[\text{GdmS}]^\alpha\left(\chi_{\text{mRNA-GdmS}} - \chi_{\text{mRNA-solvent}} - \chi_{\text{GdmS-solvent}}\right) \\ &- \frac{\nu^2_{\text{mRNA}}}{\nu_{\text{solvent}}}[\text{mRNA}]^\alpha \chi_{\text{mRNA-solvent}}\Bigg)\Bigg]\cdot\left(\frac{\nu_{\text{mRNA}}[\text{mRNA}]^\alpha}{1-\nu_{\text{mRNA}}[\text{mRNA}]^\alpha - \nu_{\text{GdmS}}[\text{GdmS}]^\alpha}\right) ,\end{aligned} \tag{33}$$

where $\chi_{i-j}$ denote the Flory-Huggins interaction parameters between components $i$ and $j$, $\nu_i$ denote molecular volumes, and $C$ is a dimensionless constant. The equation (32) can then be recast in the following form:

$$\begin{aligned} r^\alpha_{\text{mRNA}} = -k'^\alpha \exp&\left(\frac{\nu_{\text{GdmS}}\cdot\nu_{\text{mRNA}}}{\nu_{\text{solvent}}}[\text{GdmS}]^\alpha\left(\chi_{\text{mRNA-GdmS}} - \chi_{\text{mRNA-solvent}} - \chi_{\text{GdmS-solvent}}\right)\right)\cdot \\ &\cdot\left(\frac{\nu_{\text{mRNA}}[\text{mRNA}]^\alpha}{1-\nu_{\text{mRNA}}[\text{mRNA}]^\alpha - \nu_{\text{GdmS}}[\text{GdmS}]^\alpha}\right) ,\end{aligned} \tag{34}$$

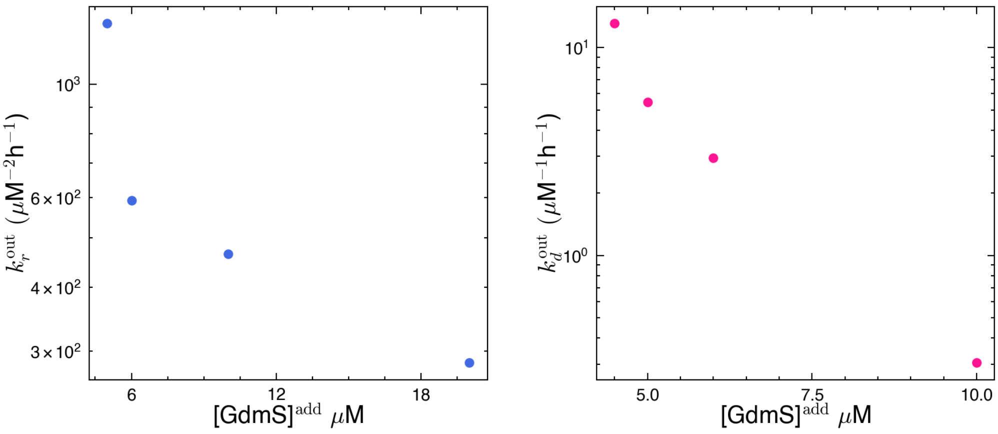

Supplementary figure 13: The rate constants in GdmS poor phase $k_r^{\text{out}}$ and $k_d^{\text{out}}$ show a roughly linear decrease when plotted on a log scale against the doped GdmS concentration.

where $k'^{\alpha}$ is a constant. Comparing Eq. (34) with the mass action form of the degradation rate $r^{\alpha}_{\text{mRNA}} = -k_d^{\alpha}[\text{RNA}]^{\alpha}[\text{RNase}]^{\alpha}$, which we defined in section 11.1, we get

$$k_d^{\alpha} = \frac{k'^{\alpha}}{[\text{RNase}]^{\alpha}} \exp\left(\frac{\nu_{\text{GdmS}} \cdot \nu_{\text{mRNA}}}{\nu_{\text{solvent}}}[\text{GdmS}]^{\alpha}\left(\chi_{\text{mRNA-GdmS}} - \chi_{\text{mRNA-solvent}} - \chi_{\text{GdmS-solvent}}\right)\right) \cdot \quad (35)$$

$$\cdot \left(\frac{\nu_{\text{mRNA}}}{1 - \nu_{\text{mRNA}}[\text{mRNA}]^{\alpha} - \nu_{\text{GdmS}}[\text{GdmS}]^{\alpha}}\right) \quad . \quad (36)$$

We can already see that $k_d^{\alpha}$ is of the form $\exp\left(p[\text{GdmS}]^{\alpha}\right)$ with

$$p = \frac{\nu_{\text{GdmS}} \cdot \nu_{\text{mRNA}}}{\nu_{\text{solvent}}}\left(\chi_{\text{mRNA-GdmS}} - \chi_{\text{mRNA-solvent}} - \chi_{\text{GdmS-solvent}}\right) \ , \quad (37)$$

given a constant volume fraction of the solvent $(1 - \nu_{\text{mRNA}}[\text{mRNA}]^{\alpha} - \nu_{\text{GdmS}}[\text{GdmS}]^{\alpha})$. In the experiments the condensate volume $V_c \ll V$, the container volume, so we can write $[\text{GdmS}]^{\text{out}} \approx [\text{GdmS}]^{\text{add}}$. Since the partition coefficient for GdmS is defined as $P = \frac{[\text{GdmS}]^{\text{in}}}{[\text{GdmS}]^{\text{out}}}$, for small condensate volumes we can write $[\text{GdmS}]^{\text{in}} \simeq P[\text{GdmS}]^{\text{add}}$. Thus, $k_d^{\alpha}$ can be effectively expressed in terms of $[\text{GdmS}]^{\text{add}}$ for each of the phases. We assume a similar form for $k_r^{\alpha}$ using a similar motivation as above.

Having estimated an exponential dependence of the rate constants on $[\text{GdmS}]^{\text{add}}$, we used the following functional form:

$$k_r^{\text{in}} = k_r^0 \exp\left(-p_1 \cdot [\text{GdmS}]^{\text{add}}\right) , \quad (38a)$$

$$k_d^{\text{in}} = k_d^0 \exp\left(-p_2 \cdot [\text{GdmS}]^{\text{add}}\right) , \quad (38b)$$

$$k_r^{\text{out}} = k_r^0 \exp\left(-p_3 \cdot [\text{GdmS}]^{\text{add}}\right) , \quad (38c)$$

$$k_d^{\text{out}} = k_d^0 exp\left(-p_4 \cdot [\text{GdmS}]^{\text{add}}\right) . \quad (38d)$$

Here $k_r^0$ and $k_d^0$ are the intrinsic rate constants for transcription and degradation in the dilute limit in the absence of $[\text{GdmS}]^{\text{add}}$. The functional form in Eq.(38a) allowed us to use $p_1$, $p_2$, $p_3$ and $p_4$ as fitting parameters across all experimental conditions together. The resulting $p_i$ were used to calculate $k_r^{\text{in/out}}$ and $k_d^{\text{in/out}}$ using Eq. (38a).

### 12 Determination of rate constants

In order to fit the kinetic traces of mRNA and GdmS expression to a fully descriptive kinetic model, represented by a set of coupled ordinary differential equations, an initial simple model

| Symbol | Parameter | Unit |
|---|---|---|
| [DNA] | DNA concentration | $\mu M$ |
| $[\text{mRNA}]_0$ | Initial mRNA concentration | $\mu M$ |
| $[\text{TsR}]_0$ | Initial transcription resource concentration | $\mu M$ |
| $[\text{TIR}]_0$ | Initial translation resource concentration | $\mu M$ |
| $k_r$ | Transcription rate constant | $\mu M^{-2}h^{-1}$ |
| $k_d$ | Degradation rate constant | $\mu M^{-1}h^{-1}$ |
| $k_p$ | Translation rate constant | $\mu M^{-2}h^{-1}$ |
| $k_{\text{mat}}$ | Maturation rate constant | $h^{-1}$ |

Supplementary table 6: Parameters used in the description of the one-phase kinetic model.

| Symbol | Variable | Unit |
|---|---|---|
| [mRNA] | mRNA concentration | $\mu$M |
| [GdmS] | Protein concentration | $\mu$M |
| $[\text{GdmS}]^*$ | Mature protein concentration | $\mu$M |
| [TsR] | Transcription resource concentration | $\mu$M |
| [TIR] | Translation resource concentration | $\mu$M |

Supplementary table 7: Concentration variables that change with time in the one-phase kinetic model.

with analytical solutions were used to fit sections of the traces to obtain initial guesses for a list of parameters that would go into the one-phase model. The parameters and variables used in the one phase kinetic model are given in table 6 and table 7.

The mRNA trace was seen to initially increase due to transcription, reach a maximum followed by decay due to degradation and depletion of resources.

#### 12.1 Determination of native T7 polymerase concentration and intrinsic transcription rate

To get initial guesses for the rate constants for transcription and degradation, each part was modelled separately. Additionally, it was assumed that the TNT extract had a native concentration of T7 RNA polymerase and RNase enzymes. These were also calculated by building a simple model for each part. For the T7 RNA polymerase titration data, the peak mRNA was determined and the section of the curve from beginning to the peak mRNA timepoint was fit to the following model. Note that we only model the rise of mRNA given by the following chemical equation:

$$\text{TsR} \xrightarrow{\text{DNA,T7}} \text{mRNA}\,. \tag{39}$$

The rate of mRNA production depends on the rate constant, DNA concentration, transcription resources and the T7 RNA polymerase.

$$\frac{d[\text{mRNA}]}{dt} = k_r[\text{DNA}][\text{TsR}][\text{T7}]\,. \tag{40}$$

Since the system has a limited amount of resources ($[\text{TsR}]_0$), the rate of TsR decay will be:

$$\frac{d[\text{TsR}]}{dt} = -k_r[\text{DNA}][\text{TsR}][\text{T7}]\,. \tag{41}$$

Let set all the constant parameters into an observed transcription rate constant $k_r^{\text{obs}} = k_r[\text{DNA}][\text{T7}]$ and proceed with analytical solutions:

$$\frac{d[\text{TsR}]}{dt} = -k_r^{\text{obs}}[\text{TsR}] \ , \tag{42}$$

which gives

$$\text{TsR}(t) = [\text{TsR}]_0 \exp\left(-k_r^{\text{obs}} t\right) \ . \tag{43}$$

Plugging this into the mRNA rate we get:

$$\frac{d[\text{mRNA}]}{dt} = k_r^{\text{obs}}[\text{TsR}]_0 \exp\left(-k_r^{\text{obs}} t\right) \ , \tag{44}$$

giving

$$[\text{mRNA}](t) = [\text{TsR}]_0(1 - \exp\left(-k_r^{\text{obs}} t\right)) \ , \tag{45}$$

assuming zero initial mRNA concentration. Note that this equation is valid only in the initial stages of the experiment when the mRNA concentration and hence the degradation rate is very small. This form was used to fit the truncated mRNA traces for different T7 RNA polymerase concentration. Owing to the truncation of the mRNA traces there was an uncertainty to the plateau of these curves, which generally would give the $[\text{TsR}]_0$ parameter. For this reason four different values of this parameter was used to fit the curves (Supplementary figure 14 (a)) and the $k_r^{\text{obs}}$ was plotted against the varying T7 RNA polymerase concentrations that were provided externally. Given that the native T7 concentration is $[\text{T7}]_{\text{native}}$ and the doped T7 polymerase is $[\text{T7}]_{\text{add}}$ . These were then fit to the equation (Supplementary figure 14 (b))

$$k_r^{\text{obs}} = k_r[\text{DNA}]([\text{T7}]_{\text{native}} + [\text{T7}]_{\text{add}}) = k_r[\text{DNA}][\text{T7}]_{\text{native}} + k_r[\text{DNA}][\text{T7}]_{\text{add}} \, . \tag{46}$$

The slope and intercept of the above equation give us $k_r$ and $[\text{T7}]_{\text{native}}$. The obtained values were : $[\text{T7}]_{\text{native}} = 470 \pm 0.07nM$, $k_r = 0.013 \pm 0.002nM^{-2}hr^{-1}$.

#### 12.2 Determination of native RNase A and intrinsic degradation rate

To get an initial guess for the degradation rate and the native RNase concentration, the mRNA degradation was modelled as:

$$\text{mRNA} \xrightarrow{\text{RNase}} \phi \, . \tag{47}$$

The rate of mRNA degradation depends on the rate constant, mRNA concentration, and the RNase concentration :

$$\frac{d[\text{mRNA}]}{dt} = -k_d[\text{mRNA}][\text{RNase}] \ . \tag{48}$$

Let the observed degradation rate constant $k_d^{\text{obs}} = k_d[\text{RNase}]$. The solution for the rate equation is:

$$\text{mRNA}(t) = [\text{mRNA}]_0 \exp\left(-k_d^{\text{obs}} t\right) \ . \tag{49}$$

Titrations of 4 different doped concentrations of RNase ($[\text{RNase}]_{\text{add}}$) on 100 nM purified mRNA added to the extract were fit to this equation and subsequently $k_d^{\text{obs}}$ was plotted against $[\text{RNase}]_{\text{add}}$ and fit to the equation $k_d^{\text{obs}} = k_d([\text{RNase}]_{\text{native}} + [\text{RNase}]_{\text{add}})$. The average fitted values of native RNase and intrinsic degradation rate constant $k_d$ across five titrations were $[\text{RNAse}]_{\text{native}}$ = $270 \pm 0.05$ nM and $k_d = 0.007 \pm 0.001 \ nM^{-1}h^{-1}$.

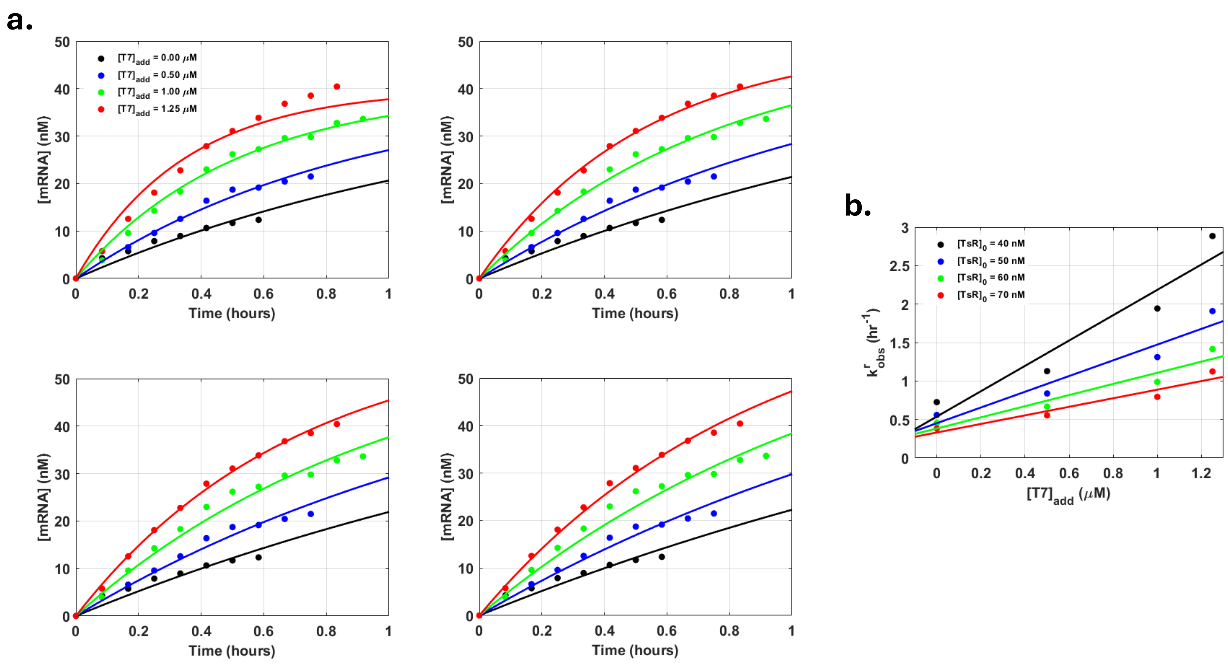

Supplementary figure 14: Determination of native T7 polymerase concentration that is already present in the TNT extract by fitting the a) rise section of the mRNA kinetic curves titrated by T7 RNA polymerase to a first order kinetic model. The initial TsR concentrations were guessed four times. b) Fitting the dependence of the added T7 RNA concentration vs the observed rate constants using a pseudo first order model gave the native T7 concentration.

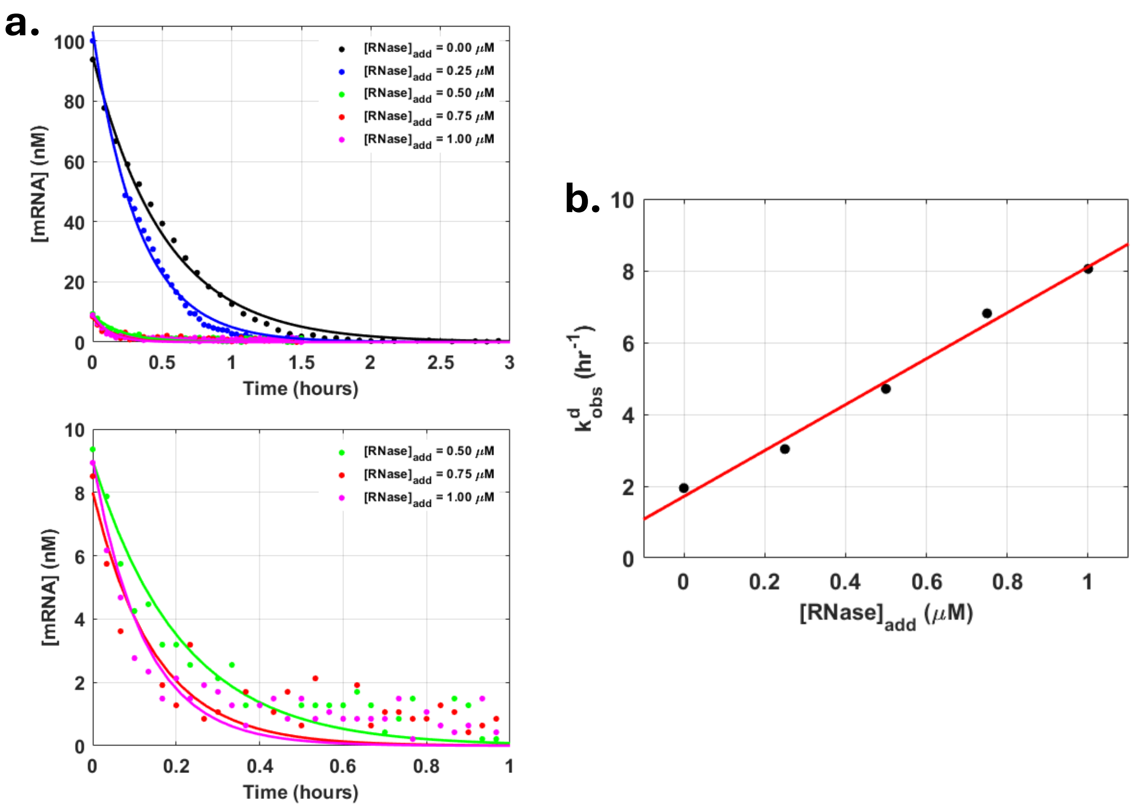

Supplementary figure 15: Degradations profiles of the decay of 100 nM mRNA in TNT extract by titrations of RNase A. a) The data was fit to a first order decay model and fitting the dependence of the observed degradation rate constant vs the added RNase A concentration to a pseudo first order model gave the native RNase A concentration in the TNT extract.

#### 12.3 Determination of translation and maturation rate constants

In order to obtain initial guesses for the translation and maturation rate constants, we modeled GdmS expression using a minimal two-step scheme in which all translation resources, including mRNA, were lumped into a single pool $\mathrm{TlR}$ that is converted to immature GdmS and then to mature GdmS*. We assumed that by the time translation picks up the mRNA concentration has peaked and the TsR concentration has declined, and therefore these variables were absorbed into $[\mathrm{TlR}]_0$ as constants:

$$\mathrm{TlR} \xrightarrow{k_p} \mathrm{GdmS} \xrightarrow{k_{\mathrm{mat}}} \mathrm{GdmS}^*.$$

The corresponding differential equations are

$$\frac{d[\mathrm{TlR}]}{dt} = -k_p[\mathrm{TlR}], \tag{50}$$

$$\frac{d[\mathrm{GdmS}]}{dt} = k_p[\mathrm{TlR}] - k_{\mathrm{mat}}[\mathrm{GdmS}], \tag{51}$$

$$\frac{d[\mathrm{GdmS}^*]}{dt} = k_{\mathrm{mat}}[\mathrm{GdmS}]. \tag{52}$$

The solution for $[\mathrm{TlR}](t)$ is

$$[\mathrm{TlR}](t) = [\mathrm{TlR}]_0\, e^{-k_p t}. \tag{53}$$

The corresponding solution for $[\mathrm{GdmS}](t)$ is

$$[\mathrm{GdmS}](t) = \frac{k_p[\mathrm{TlR}]_0}{k_p - k_{\mathrm{mat}}}\left(e^{-k_{\mathrm{mat}}t} - e^{-k_p t}\right). \tag{54}$$

Finally, the solution for the mature protein $[\mathrm{GdmS}^*](t)$ is

$$[\mathrm{GdmS}^*](t) = \frac{k_p k_{\mathrm{mat}}[\mathrm{TlR}]_0}{k_p - k_{\mathrm{mat}}}\left(\frac{e^{-k_p t}}{k_p} - \frac{e^{-k_{\mathrm{mat}}t}}{k_{\mathrm{mat}}}\right) + [\mathrm{TlR}]_0. \tag{55}$$

Using a maturation half-time for mScarlet-I of $t_{1/2}^{\mathrm{mat}} = 31.4 \pm 2.49\ \mathrm{min}$ averaged over three literature sources ([6, 7, 8]), the maturation rate constant becomes

$$k_{\mathrm{mat}} = \frac{\ln 2}{t_{1/2}^{\mathrm{mat}}} = 1.34\ \mathrm{h}^{-1}. \tag{56}$$

The expression for $[\mathrm{GdmS}^*](t)$ with this fixed maturation rate was fit to the protein expression curve for the condition without exogenous T7 RNA polymerase. This yielded ballpark estimates of the initial translation-resource pool and the translation rate constant: $[\mathrm{TlR}]_0 = 370.4 \pm 7.8\ \mathrm{nM}$ and $k_p = 0.45 \pm 0.01\ \mathrm{h}^{-1}$. Simple estimates for all rate constants and initial resource levels are summarized in Table 8.

These values were used as initial guesses for parameters in the full coupled homogeneous TXTL model, which was solved numerically and fit to the mRNA and GdmS kinetic traces.

#### 12.4 Parameter values from the kinetic model fits

The coupled differential equations for the resource limited mass action model for a one phase system (section 11.1 Transcription-translation scheme) were solved using the ode45 solver and fit using the lsqnonlin function in MATLAB 2022a (Mathworks, USA), to the mRNA and GdmS data over time for 5 different T7 RNA polymerase concentrations (470, 970, 1420, 1720, and 2040 nM), and 14 different DNA concentrations (1.4, 2.8, 4.2, 5.6, 7.0, 8.4, 9.8, 11.2, 13.9,

| Parameter | Symbol | Estimate | Units |
|---|---|---|---|
| Transcription rate constant | $k_r$ | $0.013 \pm 0.002$ | $\mathrm{nM}^{-2}\,\mathrm{h}^{-1}$ |
| Translation rate constant | $k_p$ | $0.45 \pm 0.01$ | $\mathrm{nM}^{-1}\,\mathrm{h}^{-1}$ |
| mRNA degradation rate constant | $k_d$ | $0.007 \pm 0.001$ | $\mathrm{h}^{-1}$ |
| mScarlet-I maturation rate constant (literature) | $k_{\mathrm{mat}}$ | $1.34 \pm 0.13$ | $\mathrm{h}^{-1}$ |
| Initial transcription resources | $[\mathrm{TsR}]_0$ | $320.1 \pm 2.6$ | nM |
| Initial translation resources | $[\mathrm{TlR}]_0$ | $370.4 \pm 7.8$ | nM |
| Native T7 RNA polymerase | $[\mathrm{T7}]$ | $470 \pm 0.07$ | nM |
| Native RNase | $[\mathrm{RNase}]$ | $270 \pm 0.05$ | nM |

Supplementary table 8: Parameter estimates used as initial guesses for the coupled homogeneous transcription-translation model.

| Symbol | Parameter | Value | Unit |
|---|---|---|---|
| $[\mathrm{TsR}]_0$ | Initial transcription resource concentration | 168.6 ± 77.02 | $nM$ |
| $[\mathrm{TIR}]_0$ | Initial translation resource concentration | 778.1 ± 127. | $nM$ |
| $k_r$ | Transcription rate constant | 0.012 ± 0.005 | $nM^{-2}h^{-1}$ |
| $k_d$ | Degradation rate constant | 0.022 ± 0.018 | $nM^{-1}h^{-1}$ |
| $k_p$ | Translation rate constant | 0.081 ± 0.048 | $nM^{-2}h^{-1}$ |
| $k_{\mathrm{mat}}$ | Maturation rate constant | $0.769 \pm 0.13$ | $h^{-1}$ |
| $k_b$ | DFHB1 binding rate constant | 2.490 ± 1.262 | $h^{-1}$ |
| $k_{ub}$ | DFHB1 unbinding rate constant | 0.044 ± 0.044 | $h^{-1}$ |
| $k_1$ | First maturation rate constant | 28.503 ± 5.602 | $h^{-1}$ |
| $k_2$ | Second maturation rate constant | 1.752 ± 0.206 | $h^{-1}$ |
| $k_3$ | Third maturation rate constant | 3.864 ± 1.907 | $h^{-1}$ |
| $k_4$ | Fourth maturation rate constant | 3.968 ± 2.01 | $h^{-1}$ |
| $k_p^{\mathrm{overall}}$ | Overall translation rate constant | 0.074 ± 0.037 | $h^{-1}$ |
| $\tau_{\mathrm{mat}}^{1/2}$ | Maturation half-life | 55.80 ± 9.743 | min |

Supplementary table 9: Optimal values of all parameters obtained from the one phase kinetic model.

16.7, 19.5, 22.3, 25.1, and 27.9 nM). Estimated values mentioned in table 8 were used as initial guesses. The fits were done individually as well as globally for the T7 RNA polymerase titration set and the DNA titration set. The optimal values of all parameters are given in table 9.

The intrinsic transcription rate constant was $k_r$ = 0.012 ± 0.005 $nM^{-2}h^{-1}$ and the intrinsic degradation rate constant was $k_d$ = 0.022 ± 0.018 $\mathrm{nM}^{-1}\mathrm{h}^{-1}$, suggesting similar rates of mRNA production and degradation. Building upon these values, the rates could be modulated by doping in extra T7 or RNase. The miscellany of other parameters not particularly important for phase separation were as follows: DFHB1 binding rate constant was $k_b$ = 2.490 ± 1.262 $h^{-1}$ and the unbinding rate constant was $k_{ub}$ = 0.044 ± 0.044 $h^{-1}$ suggesting DFHB1 binding was not rate-limiting. The translation rate constant for GdmS production was $k_p$ = 0.081 ± 0.048 $h^{-1}$ and the overall maturation half-life of mScarlet-I to be 55.80 ± 9.743 minutes, which was comparable to the literature average of 31.4 ± 2.49 minutes, from several sources (([6, 7, 8])). Together with the maturation rate constant, the overall expression rate constant for GdmS* was 0.074 ± 0.037 $h^{-1}$. Finally, the initial resource concentrations, and were 168.6 ± 77.02 nM and 778.1 ± 127.2 nM respectively suggesting greater resources for translation. This however was inconsequential since because in the phase separated regime, the translation pathway is negligibly contributive as the doped concentration of GdmS is much higher.

Apart from the fitting results, a secondary validation of the model came by way of the fol-

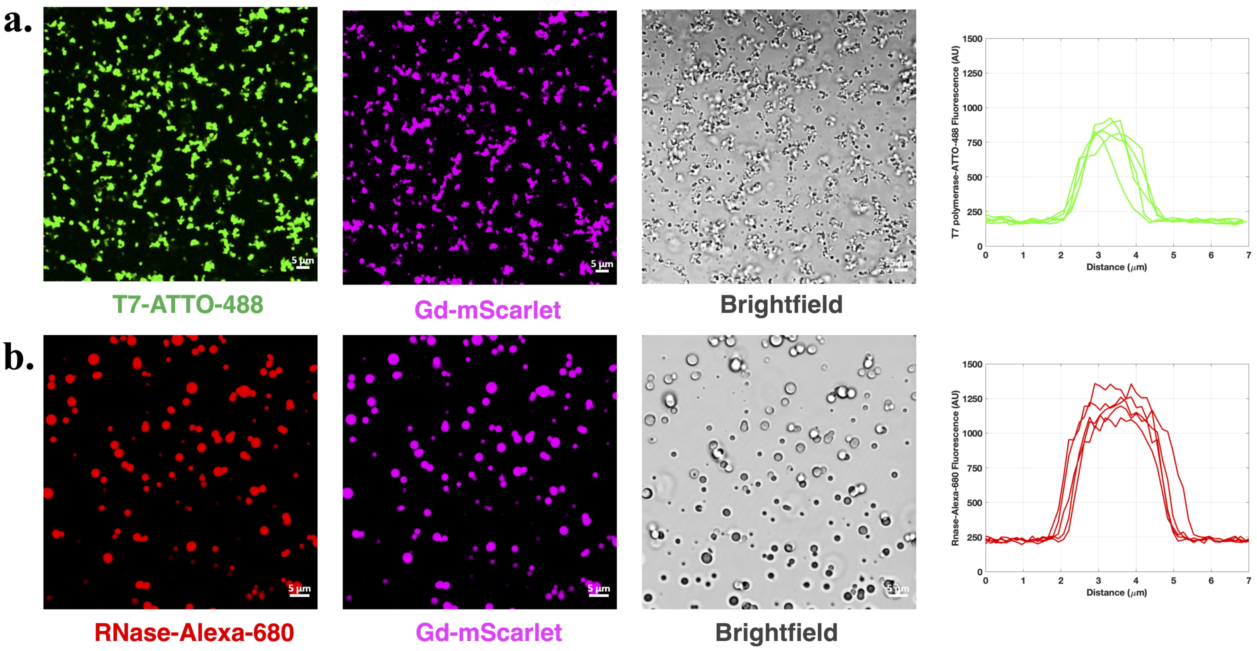

Supplementary figure 16: Confocal micrographs of GdmS-mRNA condensates doped with labelled T7 polymerase and RNase. a) Shows the partitioning of T7 polymerase labelled with ATTO-488 dye. b) Shows the partitioning of RNase labelled with Alexa Fluor 680. The partition coefficients were determined from the corresponding profiles on the right for both giving partition coefficients of 4.15 $\pm$ 0.05 (T7 polymerase) and 4.81 $\pm$ 0.04 (RNase A).

lowing experimental trend: Increasing the transcription rate by doping higher T7 concentrations led to elevated mRNA production but a concomitant decrease in GdmS expression, suggesting a shared demand on the transcriptional resource (TsR). Incorporating this dependency into the model by making the translation rate contingent on both translational (TIR) and transcriptional (TsR) resources enabled the model to reproduce this effect. Moreover, DNA titration experiments revealed an optimal DNA concentration for maximal GdmS expression. Using the same formulation with TsR-dependent translation, the model predicted a peak GdmS concentration at 9.4 nM DNA, closely matching the commercially recommended value of 5.6 nM. Together, these results reinforced the validity and predictive capability of the model.

#### 12.5 Experimental determination of Partition Coefficients of T7 polymerase and RNase A

Folded proteins are often known to be clients of condensates made up of RNA and/or IDPs [9], [10]. Therefore the partition coefficient of the enzymes T7 and RNase would be a necessary factor in the rates of transcription and degradation in the GdmS rich phase. For this reason labeled T7 and RNase, fluorescent in confocal microscopy detection channels exclusive to the mScarlet-I channel, were doped into mRNA and GdmS condensates prepared in the extract. Higher fluorescent intensity of their respective channels for both labelled T7 and RNase inside the condensates suggested that the condensates are enrich both enzymes (Figure 16). Condensates were then formed by mixing 10 $\mu$M GdmS with 0.1 $\mu$M mRNA in buffer with 1 $\mu$M ATTO-488-labeled T7 RNA polymerase (T7-ATTO488) or 5 $\mu$M Alexa Fluor 680-labeled RNase A (RNase-AF680) included in the mix. For T7-ATTO-488 detection, excitation was performed using a 488 nm laser and fluorescence was collected in the 500–550 nm emission window. RNase-AF680 was excited using a 633 nm laser and detected in the 650–700 nm range. In both cases, a sequential

| Parameter | Mean | Model error |
|---|---|---|
| $p_1$ ($\mu M^{-1}$) | 0.109 | 0.002872 |
| $p_2$ ($\mu M^{-1}$) | 0.3066 | 0.079 |
| $p_3$ ($\mu M^{-1}$) | 0.109 | 0.002872 |
| $p_4$ ($\mu M^{-1}$) | 0.6826 | 0.3423 |
| $[\text{TsR}]_0(\mu M)$ | 1.1917 | 0.02329 |

Supplementary table 10: Optimal parameter values obtained from the two phase kinetic model

| Doped GdmS ($\mu$M) | $k_r^{\text{out}}$ ($\mu M^{-2}h^{-1}$) | $k_d^{\text{out}}$ ($\mu M^{-1}h^{-1}$) | $k_r^{\text{in}}$ ($\mu M^{-2}h^{-1}$) | $k_d^{\text{in}}$ ($\mu M^{-1}h^{-1}$) |
|---|---|---|---|---|
| 4.5 | 1224.64 | 1.83455 | 1224.64 | 0.3378 |
| 5 | 1159.68 | 1.5738 | 1159.68 | 0.2401 |
| 6 | 1039.92 | 1.1582 | 1039.92 | 0.1213 |
| 10 | 672.43 | 0.3398 | 672.43 | $7.9 \cdot 10^{-3}$ |
| 20 | 226.08 | 0.01584 | 226.08 | $8.6 \cdot 10^{-6}$ |

Supplementary table 11: Rate constants evaluated using the two phase kinetic model for different doping concentrations of GdmS.

scan was performed for the mScarlet-tagged IDP (GdmS) using a 561 nm excitation laser and 579–650 nm emission detection range. Especially for the RNase-AF680 case, imaging was executed fast before the condensates dissolved away. Next, the fluorescence intensities within and outside condensates were calculated by regions of interest (ROIs) analysis of multiple droplets and surrounding GdmS poor phases. Mean fluorescence intensities were extracted for each region and the partition coefficient was approximated to be the ratio of the mean fluorescence intensity inside the condensate to that outside (Supplementary figure 16). The partition constant K was thereupon used in two phase model to convert the T7 RNA polymerase concentration and RNase concentration inside as $[\text{T7}]_{\text{in}} = P_{\text{T7}}[\text{T7}]_{\text{tot}}$ with the approximation that $[\text{T7}]_{\text{tot}} \approx [\text{T7}]_{\text{out}}$ And, $[\text{RNase}]_{\text{in}} = P_{\text{RNase}}[\text{RNase}]_{\text{tot}}$ with the approximation that $[\text{RNase}]_{\text{tot}} \approx [\text{RNase}]_{\text{out}}$.

#### 12.6 Obtaining rate constants from the two phase kinetic model

The parameters $p_1$ through $p_4$ and initial transcription resource concentration, $[\text{TsR}]_0$ , were used as tunable fitting parameters to match the condensate volume over time $V_d(t)$ to experimentally obtained data as given by the section 'Quantification of condensate volume dynamics'. Since the GdmS concentration remained fairly constant due to the excessive amount of doped concentration compared to that produced by expression, it was accepted that the condensation dynamics was purely governed by mRNA production and degradation. Thus, the dependence of $k_r^\alpha$ and $k_d^\alpha$ on the doped GdmS concentration was determined as described in section 11.6. A global fit of the droplet-phase volume was performed across all eight datasets, viz. with condition sets ($[\text{GdmS}]_{\text{add}}$ = 4.5, 5, 6, 10 and 20 $\mu$M with no additional T7 or RNase added, and $[\text{GdmS}]_{\text{add}}$ = 5, 6, and 10 $\mu$M with $[\text{T7}]_{\text{add}}$ = 1 $\mu$M and $[\text{RNase}]_{\text{add}}$ = 0.1 $\mu$M), simultaneously optimizing the five parameters $p_1, p_2, p_3, p_4$ and $[\text{TsR}]_0$. We used 'optimize.curvefit' package from the scipy library in python to obtain the global fit.

The optimized parameter values from the two phase model are given in tables 10 and 11.

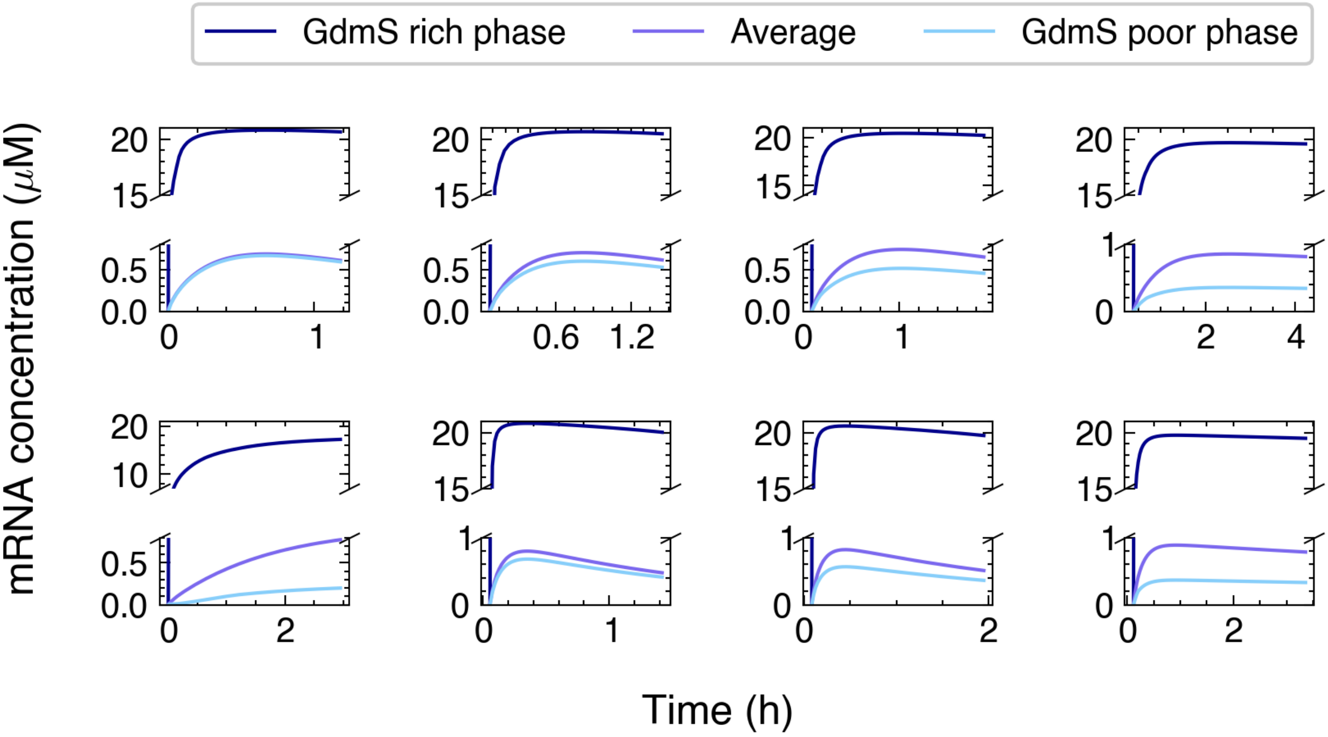

Supplementary figure 17: Time traces for [mRNA] inside the GdmS rich phase, in the GdmS poor phase and the average concentration for different concentrations of GdmS, T7 polymerase and RNase A obtained from the simulation.

### 13 RNase A based dissolution assay

For the RNase A based condensate dissolution assay, preformed condensates were assembled by mixing 0.1 $\mu$M mRNA with 10 $\mu$M GdmS in TnT extract. RNase A was added at final concentrations of 1, 2, 5, or 10 $\mu$M. After mixing, the samples were loaded and sealed as described above, and 3D Z-stacks were collected over the course of 1 hour, which was enough time for the condensates to dissolve completely. The 4 data sets were plotted with volume fraction vs time multiplied by the corresponding RNase A concentration. The data, now adjusted for RNase A concentration was fit to a first order decay curve with the RNase A concentration as a known input parameter:

$$V_d(t) = V_d(t=0)\exp\left(-k_d[\text{RNase}]t\right) . \quad (57)$$

The $k_d$ values obtained for each decay curve was averaged and the standard error was calculated.

### 14 RNase activity dependence on condensate volume

#### 14.1 RNAse activity in the supernatant

To investigate the dependence of mRNA degradation kinetics on condensate volume, a 20 µL solution of GdmS-mRNA condensates made in 80% v/v TNT extract with 0.1 $\mu$M mRNA and 10 and 17.4 $\mu$M GdmS with 5 $\mu$M RNase A added were prepared and centrifuged at 2000 ´g for 10 minutes. 10 µL of the supernatant was carefully aspirated out without disturbing the condensate pellet. Additional mRNA and the DFHBI dye was added to the supernatant to final concentration

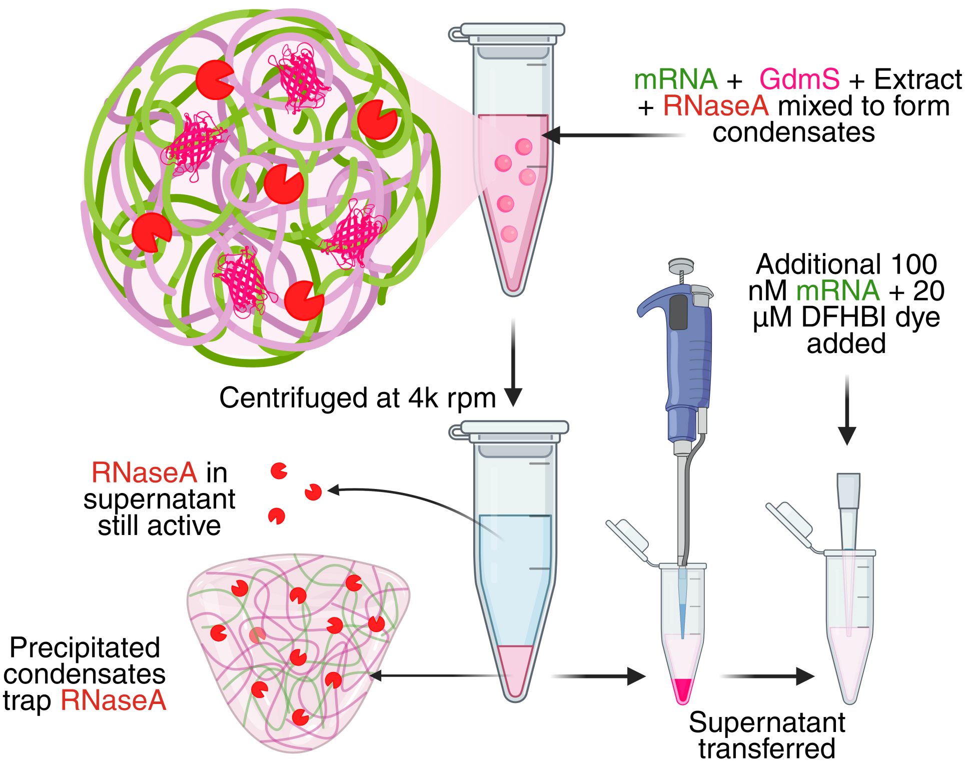

Supplementary figure 18: Schematic describing the mRNA degradation assay by centrifugation of condensates.

of 0.1 $\mu$M and 20 $\mu$M respectively. The mRNA degradation was monitored using the TECAN micro-wellplate reader with settings as described before in section 9.2 (Cell free expression and quantification of mRNA and GdmS (A schematic of the assay is given in Supplementary figure 18). One control sample was done where there was no condensate added. Additionally, the condensate volumes for the two conditions were estimated by previously described 3D confocal Z-stack analysis using Imaris (Supplementary figure 19).

The mRNA concentrations vs time were fit to a first order decay as:

$$[\text{mRNA}](t) = [\text{mRNA}]_0 \exp\left(-k_d^{\text{obs}} t\right) . \tag{58}$$

The observed rate constant $k_d^{\text{obs}}$ was then plotted against the corresponding condensate volume fractions and fit using the following logic.

We devised a theoretical relationship between $k_d^{\text{obs}}$ and $V_f$ with total RNAse A with the parameters ($[\text{RNase}]_{\text{tot}}$) and RNAse A partition coefficient ($K_{\text{RNase}}$).

Given that the native RNase concentration was 0.27 $\mu$M as determined from the section 'Simple model for transcription, mRNA degradation and translation for estimating parameter guesses', and the added RNase concentration was 0.10 $\mu$M, the total RNase concentration in the 20 µL sample was 0.37 $\mu$M. In order to interpret the inverse correlation between condensate volume and decay rate we noted that the RNase present in the supernatant is responsible for degrading free mRNA; thus, its effective concentration is expected to decline as increasing amounts partition into growing droplets. Assuming equilibrium partitioning, the RNase concentration in the supernatant or GdmS poor phase could be expressed in terms of the total RNase concentration $[\text{RNase}]_{\text{tot}}$, the partition coefficient $K_{\text{RNase}}$ and the volume fraction $V_f$:

$$[\text{RNase}]_{\text{sup}} = \frac{[\text{RNase}]_{\text{tot}}}{1 + (K_{\text{RNAse}} - 1)V_f} . \tag{59}$$

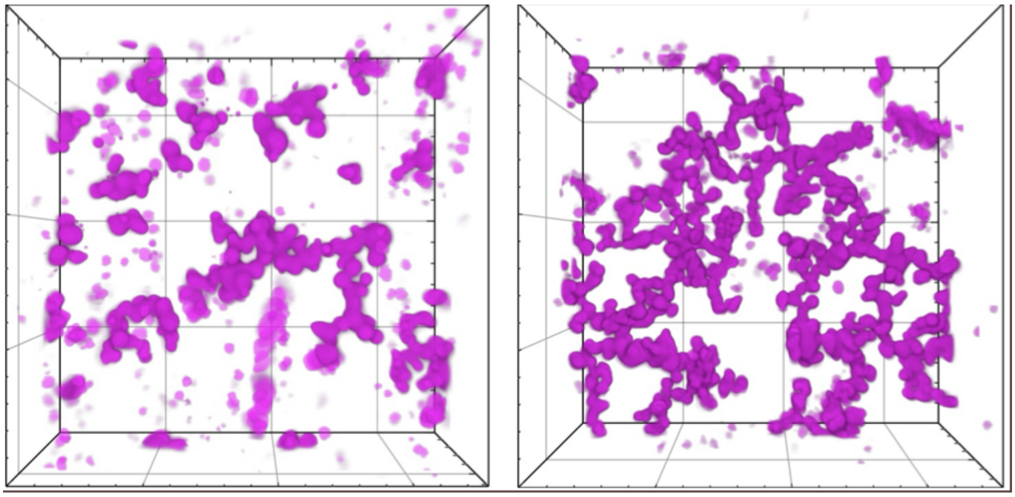

Supplementary figure 19: 3D confocal micrographs of condensates prepared by mixing $100\,$nM mRNA with $10\,\mu$M GdmS (left) and $17\,\mu$M GdmS (right). The dimensions of each micrograph is $106\,\mu$m $\times$ $106\,\mu$m.

Given that the observed degradation rate constant of mRNA degradation in the supernatant is simply $k_d^{\text{obs}} = k_d[\text{RNase}]_{\text{sup}}$, we get

$$k_d^{\text{obs}} = k_d \frac{[\text{RNase}]_{\text{tot}}}{1 + (K_{\text{RNAse}} - 1)V_f} \,. \tag{60}$$

Fitting data to a relation of the form $k_d^{\text{obs}} = \frac{A}{1+BV_f}$ gave an intrinsic degradation rate constant of $k_d$ = $(5.33 \pm 0.01) \times 10^{-3}$ nM$^{-1}$h$^{-1}$ which was comparable (within 20%) to the degradation rate constant at $(4.29 \pm 0.05) \times 10^{-3}$ nM$^{-1}$h$^{-1}$ obtained from the homogenous model. The difference in the degradation rate constants obtained from two different routes could be attributed to differences in mRNA and GdmS concentrations in the GdmS poor phase. Despite this the results further confirm that the majority of mRNA degradation occurs in the GdmS poor phase despite the high partitioning of RNAse A into the condensate. It is pertinent to note that $P_{\text{RNAse}}$ (41.2 ± 7.0) obtained from this theoretical approach was an order of magnitude greater than our experimentally measured $P_{\text{RNAse}}$ (4.81 ± 0.04). This discrepancy could be attributed to limitations of using fluorescence intensity to measure partition coefficients via effects from the condensate on the fluorophore's quantum efficiency or changes to molecular partitioning driven by the fluorophore. In addition, the partition coefficient could be underestimated if the system has not fully equilibrated.

### 15 Refractive Index Imaging of Condensate Degradation by Holotomography

In order to get insight into the density of the condensates during RNase A degradation, a holotomographic microscope (Tomocube) was used to investigate specific solutions. All reactions were carried out in 80% (v/v) TnT extract diluted in nuclease-free water. Experiments were performed at $30^0$C using an environmental incubation chamber integrated into the microscope stage. Phase-separated condensates were prepared by mixing 10 $\mu$M GdmS with 0.1 $\mu$M mRNA. For the control experiments, two condensate suspensions made with 10 $\mu$M GdmS and 0.1 $\mu$M standard mRNA (2.8 knt) and separately with a shorter variant (1.8 knt) were also imaged and analyzed. The refractive indices and the densities calculated as described before, inside and outside the condensates were plotted (Supplementary figure 20). RNase A was added to a final concentration of 5 $\mu$M to induce enzymatic degradation. The samples were loaded onto slides with 0.17 mm coverslip thickness. 2D time-lapse sequences were then acquired using the Tomocube HT-X1 system for approximately 20 minutes. The microscope captures holograms at multiple illumination angles via a digital micromirror device (DMD) and reconstructs these into refractive index maps using a proprietary inverse light-scattering algorithm. The reconstructed RI distribution reflects the spatial variation in macromolecular concentration within and around condensates. A jet colormap was applied to the RI maps to visually encode refractive index values, enabling intuitive visualization of density changes over time. The refractive index (n) is linearly proportional to the mass density ($\rho$) of macromolecules in solution according to the relation, with being the refractive index increment of the solution, itself being a ratio between change in refractive index, dn, with respect to change in mass concentration of the solute dc:

$$n = n_0 + \alpha\, \rho\,, \tag{61}$$

where $\alpha = \frac{dn}{dc}$.

Segmented volumes of condensates were obtained by applying RI thresholding across the reconstructed 3D tomograms. Within each segmented volume, the system calculated dry mass using a refractive index increment (RII) assumption corresponding to the biochemical nature of the material: 0.19 fL/pg for protein-dominated. The total dry mass and volume were thus extracted from the full segmented 3D object giving the density of each condensate.

All images were generated with the RI visualized via jet colormaps set at a range from 1.335 to 1.365.

### 16 Transcription and degradation kinetics with unlimited transcriptional resource (TsR) supply

When treating the system as an open system, i.e., with resource input permitted, two TsR supply regimes were considered: continuous (constant) supply and periodic (oscillatory) supply. In both cases, the primary output analyzed was the time-averaged mRNA concentration after oscillatory behavior reached steady-state. We considered both phase-separating and non-phase-separating (homogeneous) conditions. For each condition, both the mRNA concentration and the relative amplitude of oscillations (normalized to the mean) were evaluated, with corresponding analyses extended to the condensate volume fraction.

In the constant supply regime, TsR was added to the system at a fixed supply rate , and its temporal evolution was governed by the equation:

$$\frac{d[\text{TsR}]}{dt} = \frac{V_c r^{\text{in}}_{\text{TsR}} + (V - V_c) r^{\text{out}}_{\text{TsR}}}{V_c} + S\,. \tag{62}$$

where $r^{\text{in}}_{\text{TsR}}$ and $r^{\text{out}}_{\text{TsR}}$ are the consumption rates of TsR in the GdmS rich and GdmS poor phases, respectively (see section 11.2), and $V_c$ and $V$ are the volumes of the GdmS rich phase and compartment respectively.

In the oscillatory regime, a square wave supply profile was implemented with period . During the first half of the cycle (0 to $\tau/2$), the rate at which TsR was added to the system was S$-A$:

$$\frac{d[\text{TsR}]}{dt} = \frac{V_c r^{\text{in}}_{\text{TsR}} + (V - V_c) r^{\text{out}}_{\text{TsR}}}{V} + S - A\,. \tag{63}$$

In the second half of the cycle ($\tau/2$ to $\tau$), TsR was supplied at a rate S+$A$, resulting in:

$$\frac{d[\text{TsR}]}{dt} = \frac{V_c r^{\text{in}}_{\text{TsR}} + (V - V_c) r^{\text{out}}_{\text{TsR}}}{V} + S + A\,. \tag{64}$$

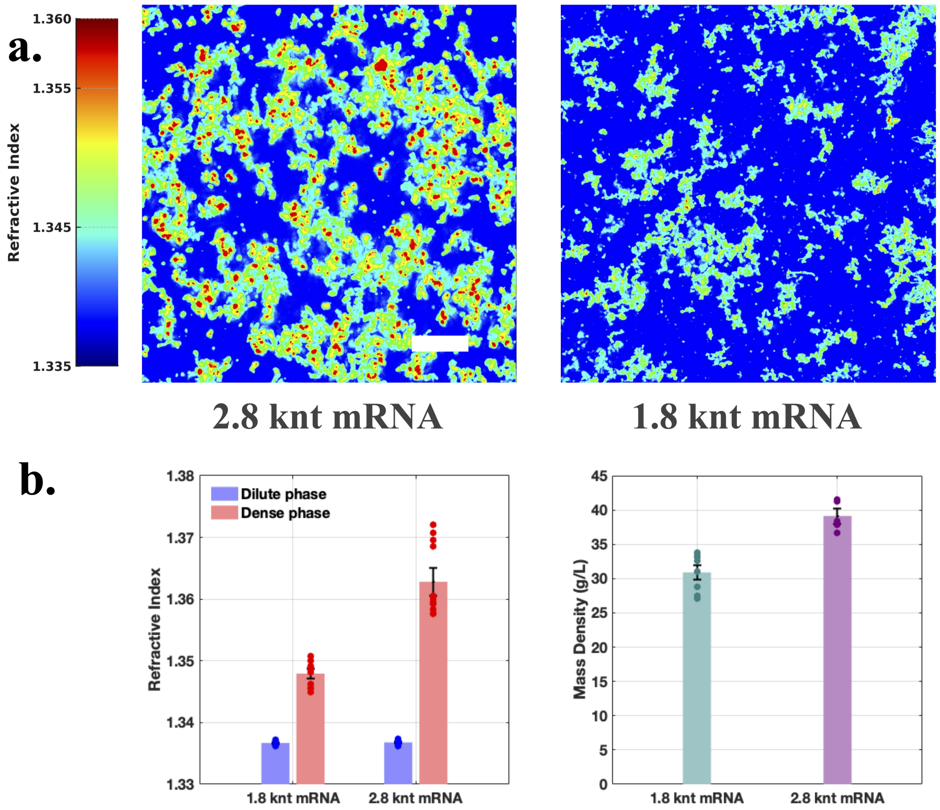

Supplementary figure 20: a) Holotomographic micrographs of GdmS-mRNA condensates made by mRNA of two different lengths, 1.8 kilonucleotides and 2.8 kilonucleotides. The color bar represents the refractive index value. b) Region of interest analysis inside the condensate and outside for both systems gave disparate overal local refractive indices from which c) the mass density was obtained demonstrating that shorter mRNA would cause the refractive index inside the condensats to be lower and therefore density too.

Our model was always initialized with zero mRNA and TsR. As TsR accumulated, mRNA production was initiated. Upon reaching supersaturation, phase separation occurred (if permitted), splitting the system into GdmS poor and GdmS rich phases with phase-dependent kinetic parameters (see section 11.3). In contrast, under non-phase separating conditions, the system was constrained to remain homogeneous regardless of supersaturation, and kinetic rates corresponding to the GdmS poor phase were applied throughout. The [TsR] at the next time step can then be determined according to Euler integration :

$$[\text{TsR}](t+dt) = [\text{TsR}](t) + \frac{d[\text{TsR}]}{dt}dt \ , \tag{65}$$

where $\frac{d[\text{TsR}]}{dt}$ is given by either Eq.(62), Eq.(63) or Eq.(64), depending on the type of external supply of TsR.

All rate constants and phase diagram parameters were taken from global fits previously described. Depending on the amount of TsR present at any given moment, the rate of change of [mRNA] was determined as described in section 11.4. We compared [mRNA] for no external TsR supply, constant supply and oscillating supply. For each condition we compared the scenarios where the system was allowed to phase separate, and where it was forced to remain homogeneous.

For a representative value of frequency and supply amplitude, we plotted [mRNA] as a function of time for the phase separated and homogeneous scenarios, and compare the cases of no TsR input, constant input and oscillating input. In case of a oscillating supply, both the frequency $\tau^{-1}$ and amplitude $A$ of the TsR supply rate were systematically varied to characterize the system's kinetic filtering behavior. After [mRNA] reached steady oscillations around a constant mean, we took the time average of [mRNA]

$$\overline{[\text{mRNA}]} = \frac{\int_t^{t+\tau} [\text{mRNA}](t)dt}{\tau} \tag{66}$$

where $\tau$ is the period of oscillations. We evaluated the quantity $\overline{[\text{mRNA}]}$ as a function of oscillation frequencies $\tau^{-1}$ and amplitudes $A$. The quantity depicted in Figure 5A1-A3, B2-B3 is $\overline{[\text{mRNA}]}$.

### 17 Supplementary note: Determination of GdmS-mRNA condensate material properties using optical tweezer based passive microrheology

Given that previous studies have shown that condensate density can influence primitive ribozyme activity [3], we measured the viscoelastic properties of the condensate environment. To do this, we conducted passive microrheology by optically trapping carboxylate-coated beads within condensates prepared in extract, following the previously established pMOT protocol [11]. Two additional control conditions were tested: extract alone and extract supplemented with GdmS only.

Passive microrheology with optical tweezers was used to estimate the frequency-dependent viscoelastic moduli and zero-frequency viscosity of GdmS–mRNA condensates in TNT extract. Samples were prepared by diluting TNT extract to 80% (v/v) in nuclease-free water, with and without phase-separated condensates. Carboxylate-coated polystyrene beads with a diameter of $2\ \mu\text{m}$ were added to each sample. Condensates were allowed to settle, and individual beads

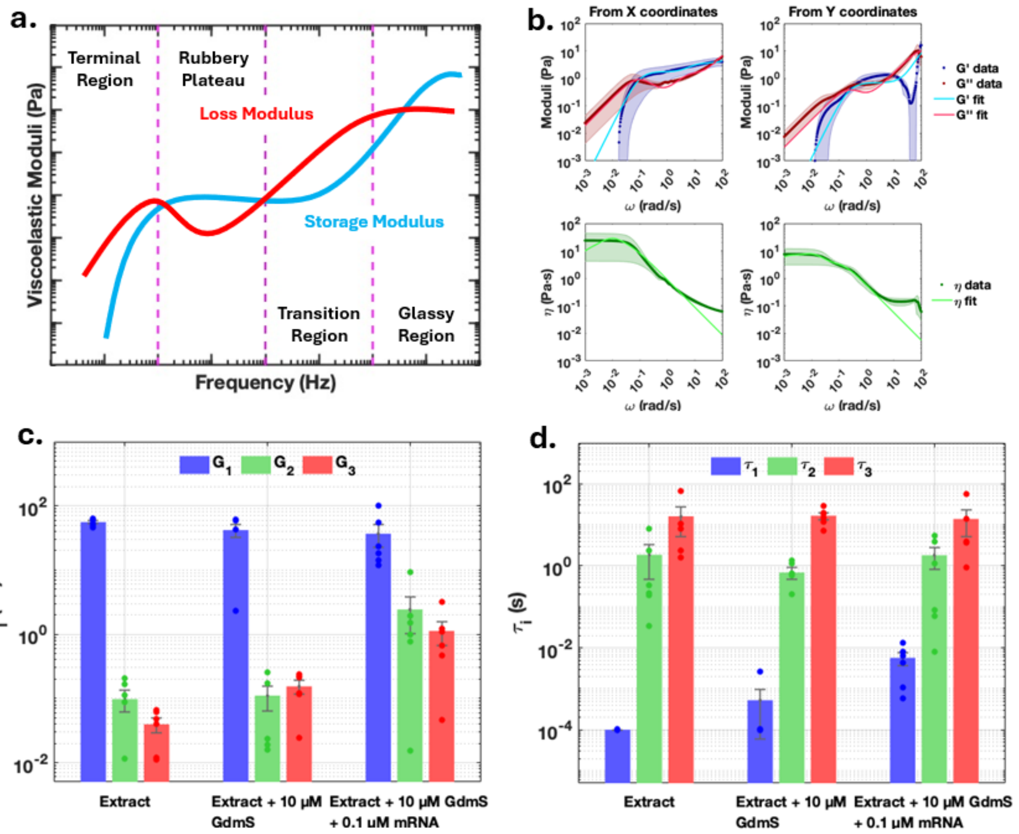

Supplementary figure 21: Passive bead microrheology experiments with optical tweezers (pMOT). (a) Representative frequency-dependent elastic (storage) and viscous (loss) moduli of a gel-like material, illustrating distinct mechanical response regimes (adapted from [12]). (b) Frequency-dependent viscoelastic moduli obtained from the $x$- and $y$-coordinate fluctuations of trapped beads, fitted with a Prony-series model for extracts containing GdmS–mRNA condensates. (c) Elastic (Prony) moduli $G_i$ extracted from the fits. (d) Corresponding relaxation times $\tau_i$.

embedded within large condensates were identified and trapped using a Lumicks C-Trap optical tweezer system (Lumicks C-Trap, Netherlands). The trap laser power was maintained at approximately 5% to ensure stable confinement without excessive heating.

High-speed imaging of bead motion was performed for durations of 10–20 minutes per replicate and condition. Frame acquisition rates were optimised by adjusting camera cropping, pixel clock, and exposure time, typically reaching $\sim 200$ frames per second. Using the bead-position recognition feature in the Lumicks BlueLake operational software, the bead $x$–$y$ trajectory over time was exported as `.hdf` files and analysed using a custom Python script as described by Alsareedah et al. The frequency-dependent storage modulus $G'(\omega)$ and loss modulus $G''(\omega)$ were computed from the $x$–$y$ position data over a frequency range spanning $0.001$ to $100$ Hz. These viscoelastic spectra were fit using a three-component Prony series (generalised Maxwell model), which describes the complex shear modulus $G^*(\omega)$ as

$$G^*(\omega) = G_\infty + \sum_{j=1}^{3} \frac{G_j \, i\omega\tau_j}{1 + i\omega\tau_j} \,. \tag{67}$$

Here, $G_\infty$ is the high-frequency plateau modulus, and each Maxwell element is defined by a modulus $G_j$ and a characteristic relaxation time $\tau_j$. The model captures viscoelastic behaviour over multiple time scales by summing three such elements in parallel.

Separating into real and imaginary components yields the elastic (storage) and viscous (loss) moduli, $G'(\omega)$ and $G''(\omega)$:

$$G'(\omega) = G_\infty + \sum_{j=1}^{3} \frac{G_j\, \omega^2 \tau_j^2}{1 + \omega^2 \tau_j^2}\,, \tag{68}$$

$$G''(\omega) = \sum_{j=1}^{3} \frac{G_j\, \omega \tau_j}{1 + \omega^2 \tau_j^2}\,, \tag{69}$$

where each viscoelastic component represents a distinct physical relaxation process. $G_1$ and $\tau_1$ represent fast local rearrangements and short-range interactions, $G_2$ and $\tau_2$ capture intermediate-scale elastic behaviour, potentially linked to network rearrangements or local restructuring, and $G_3$ and $\tau_3$ characterise slow processes including macromolecular rearrangement and long-time fluid relaxation.

The zero-frequency (long-time) viscosity $\eta_0$ was calculated from the low-frequency asymptotic behaviour of $G''(\omega)$ as

$$\eta_0 = \lim_{\omega \to 0} \frac{G''(\omega)}{\omega} = \sum_{j=1}^{3} G_j \tau_j\,. \tag{70}$$

This effective viscosity quantifies the drag experienced at slow deformation rates and serves as a key parameter for comparison with alternative viscosity measurements such as those derived from FRAP recovery kinetics.

The fit consisted of three distinct relaxation processes modelled as three time constants, $\tau_1$, $\tau_2$, and $\tau_3$, with corresponding moduli $G_1$, $G_2$, and $G_3$. The fastest component, $\tau_1 = 5.73 \pm 1.98$ ms, likely corresponds to rapid local rearrangements such as side-chain fluctuations or solvent-mediated motions, and accordingly carried the largest modulus ($G_1 = 36.87 \pm 14.11$ Pa). An intermediate process, characterized by $\tau_2 = 1.81 \pm 0.98$ s, reflects mesoscale dynamics, possibly associated with network restructuring or local domain reorganization. The slowest process, $\tau_3 = 14.02 \pm 8.93$ s, is indicative of long-timescale rearrangements such as intermolecular sliding or fusion-like events. Both long-timescale processes, being on the order of seconds, indicate that the condensates retain structural memory over extended periods, a hallmark of viscoelastic or gel-like materials. Consistently, the condensate-containing samples exhibited higher values of moduli $G_2$ and $G_3$ (associated with $\tau_2$ and $\tau_3$) than both extract alone and extract with GdmS only, suggesting that long-timescale viscoelastic processes are more prominent in the dense environment of condensates.

We also obtained the zero-frequency viscosity of the condensates, $10.92 \pm 7.42$ Pa s, which was of the same order of magnitude as that estimated by FRAP ($9.455 \pm 1.551$ Pa s). Such high viscosities are consistent with gel-like, shear-thickening condensates as reported previously [13]. The material properties of GdmS–mRNA coacervates could therefore contribute to the attenuation of reaction kinetics within the coacervate interior, despite enzyme partitioning into the condensate, where slow diffusion and high viscoelasticity may underlie the slowdown in reaction rates. It is interesting to note that previous studies have shown that phase-separated GdmS and mRNA form liquid droplets in buffer [14]. We attribute the difference in material states between our system and previous studies to the macromolecular environment of the cell-free expression system, which is a complex cocktail rich in folded proteins and RNA. Previous literature has demonstrated that different physicochemical conditions, such as decreased salt content, increased structural content of condensates, and molecular crowding, can lead to more gel-like condensates [15].

### 18 Supplementary note: Diffusion coefficient determination by Fluorescence Recovery after Photobleaching (FRAP)

In all instances, FRAP measurements on the GdmS-mRNA condensates were carried out on the same Zeiss LSM 880 confocal microscope using the same objective and detection channel for GdmS as previously described. Two pre-bleach frames were acquired prior to photobleaching and 100% laser power at 591 nm (targeting the GdmS molecule) was used for bleaching, while 1% laser power was used for detection of the recovery.

Fluorescence recovery curves were fit to a single exponential function of the form

$$I(t) = I_\infty \left(1 - e^{-t/\tau_{\mathrm{FR}}}\right), \tag{71}$$

where $\tau_{\mathrm{FR}}$ is the fluorescence recovery time constant. From these fits, the effective diffusion coefficient $D$ of GdmS within the condensate was calculated using the Soumpasis relation

$$D = \frac{0.224\, r_B^2}{\ln 2\, \tau_{\mathrm{FR}}}, \tag{72}$$

where $r_B$ is the radius of the bleached region.

In order to convert the FRAP-derived diffusion coefficients into an apparent microviscosity, we estimated the hydrodynamic radius $R_h$ of the GdmS. First, the hydrodynamic radius of the folded mScarlet core, $R_{h,\mathrm{core}}$, was obtained directly from its crystal structure (PDB: 5LK4) using a solvent-accessible surface (SAS) construction. Atomic coordinates (excluding waters) were read, and van der Waals radii were assigned according to Bondi. A SAS volume $V_{\mathrm{SAS}}$ was computed by voxelizing the union of atomic spheres inflated by a 1.4 Å water probe. The equivalent-sphere radius associated with this SAS volume was

$$R_{\mathrm{eq,SAS}} = \left(\frac{3V_{\mathrm{SAS}}}{4\pi}\right)^{1/3}, \tag{73}$$

and the solvent-excluded radius was approximated as

$$R_{\mathrm{SES}} = R_{\mathrm{eq,SAS}} - 1.4\ \text{Å}. \tag{74}$$

Finally, a hydration shell of 3 Å was added to yield the hydrodynamic radius of the folded mScarlet section,

$$R_{h,\mathrm{mS}} = R_{\mathrm{SES}} + 3\ \text{Å} \approx 2.3\ \mathrm{nm}. \tag{75}$$

The intrinsically disordered GdmS region (425 residues) was treated as a polymer coil. Its radius of gyration was approximated using a standard scaling relation for intrinsically disordered proteins,

$$R_{g,\mathrm{IDP}} = aN^{\nu}, \tag{76}$$

with $N = 425$ residues, $\nu = 0.6$ and $a = 0.19$ nm. The choice of $a$ is based on polymer-scaling analyses of unfolded proteins and IDPs from SAXS and single-molecule FRET, which consistently report

$$R_g \approx 0.2\ \mathrm{nm} \times N^{0.6} \tag{77}$$

for polypeptide chains in good-solvent conditions. We therefore adopted $a = 0.19$ nm as a representative prefactor within this empirically established range. For the IDP Gd, this yields

$$R_{g,\mathrm{Gd}} \approx 7.2\ \mathrm{nm}. \tag{78}$$

We then converted the radius of gyration $R_g$ into a hydrodynamic radius $R_h$ using

$$R_h \approx \frac{R_g}{1.5}. \tag{79}$$

This proportionality is a standard result from polymer physics for flexible coils in good or $\theta$-solvent conditions. The hydrodynamic radius of the disordered region was then taken as

$$R_{h,\mathrm{Gd}} \approx \frac{R_{g,\mathrm{Gd}}}{1.5} \approx 4.8\ \mathrm{nm}. \tag{80}$$

The total hydrodynamic radius of the fusion protein was approximated by combining the contributions of the core and the disordered tail as

$$R_h \approx \sqrt{R_{h,\mathrm{mS}}^2 + R_{h,\mathrm{Gd}}^2} \approx 5.3\ \mathrm{nm}. \tag{81}$$

Using this hydrodynamic radius, the apparent microviscosity $\eta$ experienced by GdmS within the condensate was obtained from the Stokes–Einstein relation,

$$\eta = \frac{k_B T}{6\pi R_h D}, \tag{82}$$

where $k_B$ is the Boltzmann constant, $T = 303\ \mathrm{K}$ (30 ℃), and $R_h = 5.3\ \mathrm{nm}$. Diffusion coefficients $D$ were obtained from FRAP fits. The resulting diffusion coefficients and apparent microviscosities are summarized in Table 12.

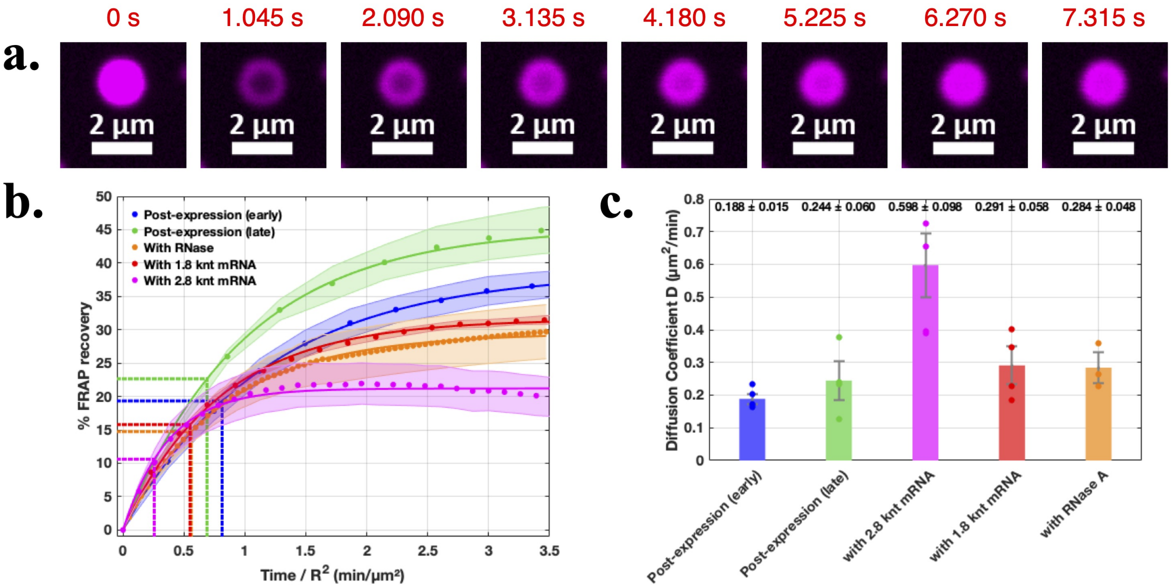

Supplementary figure 22: Fluorescence Recovery after Photobleaching of GdmS-mRNA condensates under different conditions. a) An example of a FRAP experiment on sample 2.8knt mRNA (100 nM), 10 uM for GdmS. b) Fluorescence recovery curves with normalised time. Dotted lines show the position of the half life in the time axis shows that the recovery time is fasters with the longer length mRNA. c) Diffusion coefficients obtained by the Soumpasis equation

.

| Condition | $D$ ($\mu\mathrm{m}^2\,\mathrm{min}^{-1}$) | $\eta$ (Pa s) |
|---|---|---|
| Post-expression (early) | $0.1884 \pm 0.0151$ | $13.3 \pm 1.1$ |
| Post-expression (late) | $0.2445 \pm 0.0596$ | $10.3 \pm 2.5$ |
| with 2.8 knt mRNA | $0.5976 \pm 0.0980$ | $4.2 \pm 0.7$ |
| with 1.8 knt mRNA | $0.2910 \pm 0.0583$ | $8.6 \pm 1.7$ |

Supplementary table 12: Diffusivities from FRAP data and the derived coefficients of viscosity under different conditions.
